# Tuning siRNA Specificity through Seed Region Incorporation of Deoxyribonucleotide Stereoisomers

**DOI:** 10.64898/2026.01.13.699368

**Authors:** Mehran Nikan, Thazha P. Prakash, Guillermo Vasquez, Graeme C. Freestone, Marie Annoual, Michael Tanowitz, Hongda Li, Sagar Damle, Rodrigo Galindo-Murillo, Stephanie K. Klein, Audrey Low, Clare Quirk, Colleen Y. Heller, Dorothy T. Ta, Andrew T. Watt, Michael T. Migawa, Eric E. Swayze

**Affiliations:** Ionis Pharmaceuticals Inc., 2855 Gazelle Court, Carlsbad, CA 92010, USA

## Abstract

Precise chemical design continues to drive advances in RNA-based therapeutics. Here, we report the synthesis and site-specific incorporation of four canonical phosphoramidites (U, C, A, and G), each bearing non-natural nucleoside configurations: β-D-2′-deoxyxylonucleosides and α-L-2′-deoxyribonucleosides. These stereochemically distinct nucleoside analogs were introduced at positions 6 and 7 within siRNA seed regions. When applied to siRNAs targeting *Ttr*, *ACTN1*, and *Marc1*, these modifications reduced off-target gene repression in functional assays and, in several cases, in transcriptome-wide differential expression analyses, while preserving robust on-target activity. *In vivo*, *Marc1*-targeting siRNAs containing these modified nucleosides showed decreased hepatotoxicity, as evidenced by reduced serum ALT and AST levels. Collectively, these findings establish β-D-2′-deoxyxylonucleoside and α-L-2′-deoxyribonucleoside analogs as promising chemical tools for enhancing the specificity and safety of siRNA therapeutics. This work underscores the power of integrating rational nucleoside design with comprehensive functional and *in vivo* evaluation to advance drug development based on RNA interference (RNAi).

**GRAPHICAL ABSTRACT:** 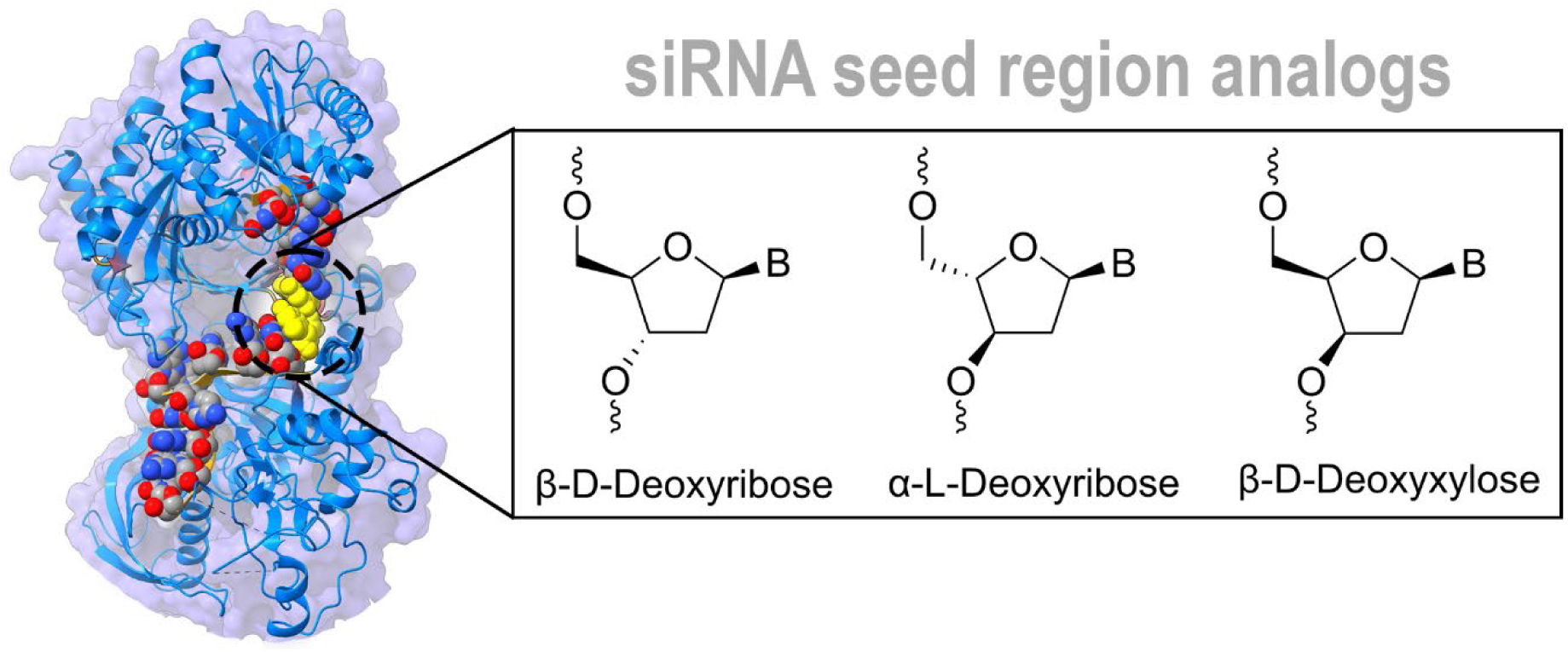

## INTRODUCTION

The chirality of DNA arises from its deoxyribose sugar, which contains three chiral centers. While eight stereoisomers are theoretically possible, DNA exclusively adopts the β-D configuration (Figure 1). This stereochemistry underpins the right-handed helices and consistent structural features observed in all living organisms. Nature’s stringent selection underscores the fundamental role of sugar configuration in determining nucleic acid structure and function and is a critical consideration when designing modified nucleotides for therapeutic applications.

**Figure 1.**
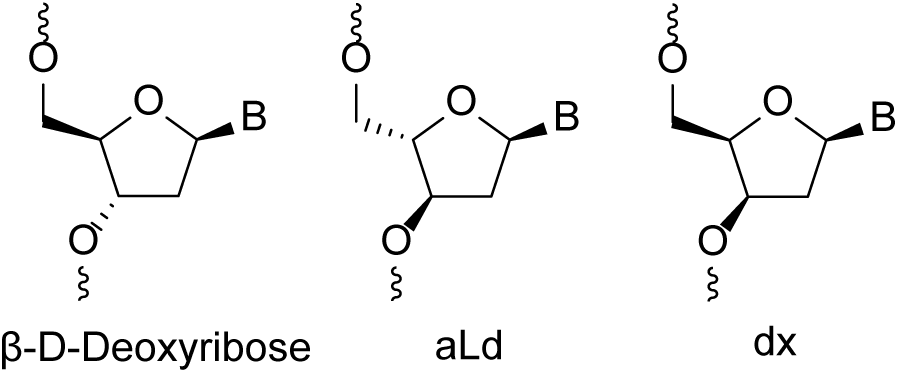
Chemical structures of β-D-deoxyribonucleoside, α-L-2′-deoxyribonucleoside (aLd) and β-D-2′-deoxyxylonucleoside (dx).

In oligonucleotide therapeutics such as small interfering RNAs (siRNAs), chemical modifications to nucleotide building blocks are essential for enhancing efficacy and minimizing adverse effects.^1^ A central challenge in siRNA design is mitigating off-target effects arising from unintended interactions between the guide strand’s seed region (nucleotides 2 to 8) and partially complementary sequences in non-target mRNAs, a phenomenon known as the miRNA-like effect.^2–4^ These interactions can lead to widespread dysregulation of gene expression and potential toxicity.^5^ Accordingly, sugar modifications that selectively modulate seed pairing to reduce off-target activity are highly desirable.^6^ While various chemical strategies have been explored to improve siRNA specificity, including our recent demonstration that a single alkyl phosphonate modification in the seed region reduces off-target activity,^7^ most sugar modifications focus on altering the overall scaffold,^8–12^ as seen in unlocked nucleic acids (UNA)^13^ or glycol nucleic acids (GNA).^14–16^ In contrast, sugar stereochemistry has received relatively little attention.

To address this gap, we selected two atypical nucleoside analogs, β-D-2′-deoxyxylonucleosides (dx) and α-L-2′-deoxyribonucleosides (aLd), (Figure 1) to investigate their impact on siRNA specificity and seed region tolerability. dx differs from natural β-D-2′-deoxyribonucleosides by inversion at the 3′ carbon, resulting in a diastereoisomeric backbone.^17^ Nucleic acids constructed from this nucleoside, known as deoxyxylonucleic acid (dXNA), have been extensively characterized, and are known to adopt unusual structures such as left-handed or ladder-like helices.^17–19^ Despite alterations in sugar pucker and backbone geometry, dXNA retains the ability to support Watson-Crick base pairing, exhibits high nuclease resistance, and shows limited cross-pairing with natural DNA or RNA, effectively functioning as an orthogonal self-pairing system.^17, 19^ While single incorporations of dx are poorly tolerated in DNA duplexes, likely due to changes in sugar pucker and torsion angles, they are better accommodated in RNA hybrids, which favor an N-type sugar pucker.^17, 20^

aLd (Figure 1) is a rare stereoisomer defined by both its L-configuration and α-anomeric attachment of the nucleobase.^21^ Homooligomers of aLd T and aLd A have been synthesized and show substantial nuclease resistance but poor hybridization with natural nucleic acids.^22^ In contrast, single incorporations of aLd remain largely underexplored, owing to the limited availability of a complete set of canonical phosphoramidites. As a result, most prior studies have focused on related analogs such as α-D-2′-deoxyribonucleosides,^21^ β-L-2′-deoxyribonucleosides,^23, 24^ or α-L-ribo-configured locked nucleic acids (α-L-LNA).^25^ Notably, single incorporations of β-L-2′-deoxyribonucleosides have been reported to enhance nuclease resistance, particularly against exonucleases, but can reduce duplex thermal stability in a sequence-dependent manner.^22^

Thus, we hypothesized that these atypical nucleoside configurations could uniquely influence siRNA function and target specificity, particularly within the seed region. To systematically assess sequence- and position-dependent effects, we synthesized all four canonical phosphoramidites for both aLd and dx, enabling their precise incorporation into siRNAs for comprehensive evaluation. This approach provides a versatile toolkit to explore the impact of sugar stereochemistry on siRNA function and establishes a foundation for future efforts to tailor siRNA performance through stereochemical modification.

## RESULTS AND DISCUSSION

Having identified dx and aLd as promising candidates, we next developed efficient routes to their canonical phosphoramidites and assessed their functional impact in siRNA.

### Synthesis of dx and aLd phosphoramidites

As outlined in Scheme 1, the synthesis of dx and aLd phosphoramidites for U, C, A, and G followed distinct, but convergent strategies, each grounded in established methodologies. For the aLd series, previous approaches typically relied on multistep transformations from protected L-arabinose or its derivatives, as L-arabinose is the only naturally abundant L-pentose.^22^ Alternative strategies based on inversion of D-sugars required full stereochemical inversion at multiple centers and often resulted in low yields or difficult-to-separate anomeric mixtures.^21, 26^ Despite these challenges, the full set of five unprotected canonical aLd nucleosides (U/T, C, A, G) was systematically synthesized by Imbach et al. as early as 1991.^27^ Their pioneering work adapted conventional nucleoside methodologies, originally optimized for the corresponding β-D-sugars, to yield unprotected monomeric nucleosides for initial structural and antiviral evaluation. These monomers, however, generally showed little antiviral activity. Nevertheless, a general and scalable synthesis providing all four canonical aLd phosphoramidites necessary for automated oligonucleotide synthesis has remained unavailable. In our approach, the synthesis began from 1-O-acetyl-2,3,5-tri-O-benzoyl-L-arabinofuranose, a compound readily accessible from L-arabinose using established methods.^22, 28^ Selective 2′-deoxygenation was carried out using the Barton-McCombie reaction,^29^ affording protected aLd nucleosides suitable for phosphitylation. This unified route provided all four canonical aLd nucleosides and their corresponding phosphoramidites in good yield and purity. To our knowledge, this represents the first complete, consistent, and scalable synthesis of all four canonical aLd phosphoramidites, enabling their systematic evaluation in siRNA and oligonucleotide therapeutics.

**Scheme 1.**
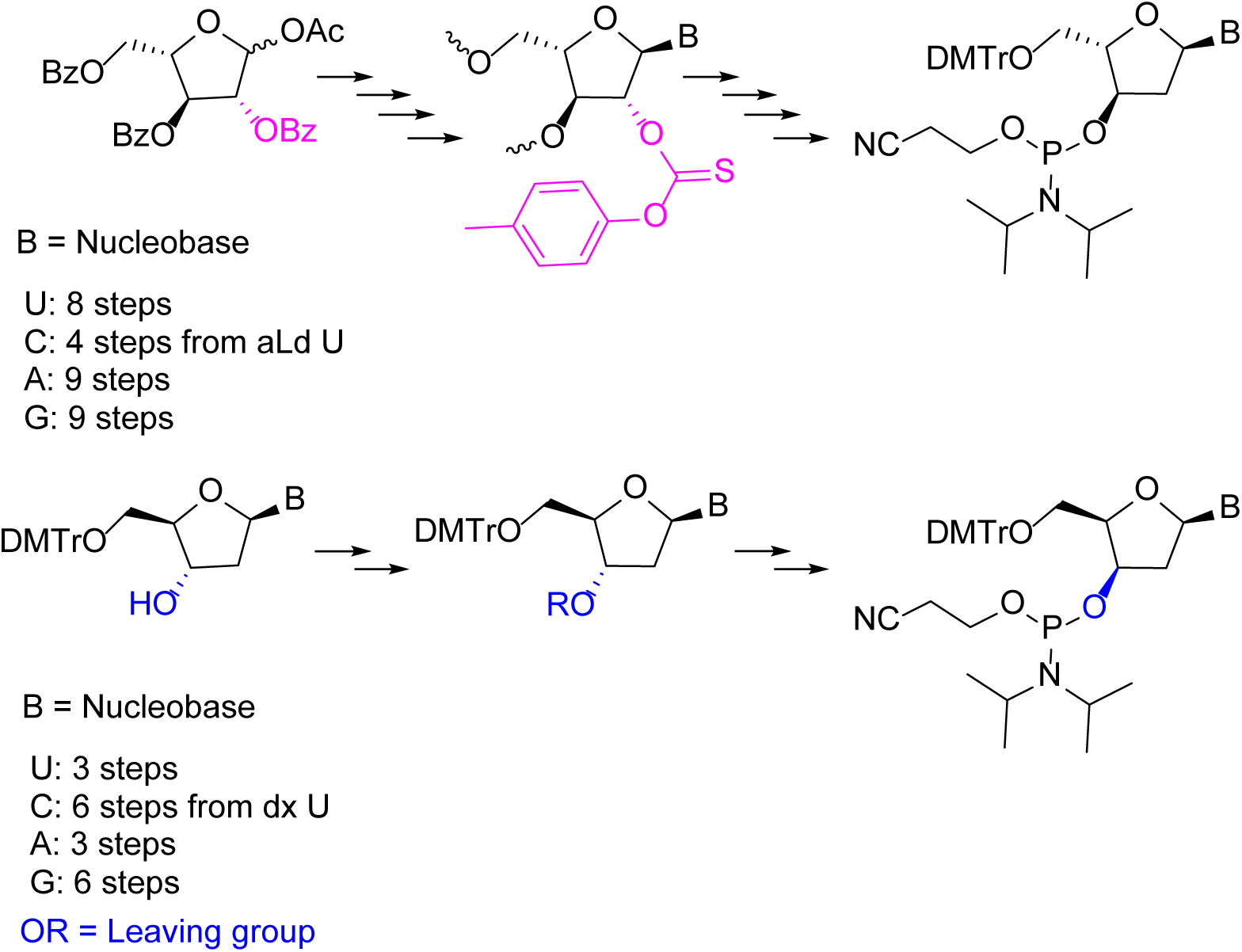
General synthetic routes to aLd (top) and dx (bottom) phosphoramidites of U, C, A, and G. Top: From 1-O-acetyl-2,3,5-tri-O-benzoyl-L-arabinofuranose via Barton-McCombie deoxygenation. Bottom: From β-D-deoxyribonucleosides via installation of a leaving group and nucleophilic displacement to effect selective 3′-inversion. Phosphitylation affords the corresponding phosphoramidites (B = U, C, A, or G). Detailed procedures are in the Supporting Information.

In designing the dx series, we selected β-D-2′-deoxyribonucleosides as common starting materials to access the corresponding phosphoramidites for all four canonical bases (U, C, A, and G). Previous work by Seela and co-workers reported the individual preparation of these monomers using base-specific synthetic strategies.^30–34^ For purines, earlier approaches such as deoxygenative [1,2]-hydride shifts^35–37^ required harsh conditions and sometimes employed pyrophoric reagents (e.g., lithium triethylborohydride, LTBH), making them difficult to handle and scale. Other purine inversion routes, such as triflate-mediated inversions, were also employed but frequently gave isomeric mixtures arising from acyl migration, which complicated purification.^38, 39^ While purification of guanosine derivatives synthesized via these approaches was often tedious due to aggregation, these methods nevertheless represented important foundational chemistry.

For pyrimidines, SN2-type displacement at the 3′-position through anhydronucleoside intermediates^40^ provided access to dx T and dx C but generally resulted in modest yields.^31, 32^ In Seela’s early work, the protected dx C phosphoramidite was obtained in only 17% yield and described as “accessible with difficulty.”^32^ Subsequent modifications to this route, often involving nucleophilic opening of anhydronucleosides, improved the efficiency of the key xylo intermediate formation and the final phosphitylation step, enabling higher overall yields.^41^

In contrast, Wang and co-workers adapted the in-situ oxidation-reduction procedure of Eisenhuth and Richert^34^ to prepare dx phosphoramidites and their precursors.^42^ This approach employed Dess–Martin periodinane oxidation at C3′ followed by stereoselective NaBH_4_ reduction under low-temperature conditions (−60 °C), minimizing depurination and providing xylo-configured nucleosides in high yield.

Recognizing the need for a unified, efficient, and scalable synthetic strategy applicable to both purine and pyrimidine nucleosides, we developed a streamlined approach. Our route begins with β-D-2′-deoxyribonucleosides and proceeds via selective inversion at the 3′-position to generate the xylo configuration through nucleophilic displacement. Subsequent phosphitylation afforded the corresponding purine and pyrimidine phosphoramidites, which were fully compatible with automated solid-phase oligonucleotide synthesis. Complete synthetic procedures for all eight phosphoramidites are provided in the Supporting Information.

With the synthetic routes established, we next evaluated how aLd and dx modifications influence siRNA specificity by introducing these nucleosides into defined positions within the siRNA guide strand.

### Dual luciferase-based off-target screening of siRNAs targeting *Ttr* and *ACTN1* functionalized with dx and aLd

To systematically investigate the influence of sugar stereochemistry on siRNA specificity, we selected guide sequences targeting transthyretin (*Ttr*) and alpha-actinin 1 (*ACTN1*), both of which have been extensively characterized in the literature for their pronounced off-target effects mediated by seed region interactions (Table 1).^7, 15, 43^ Mindful of previous work analyzing positional effects, we incorporated dx and aLd at guide-strand positions 6 and 7 in both constructs.^10, 13, 15^

**Table 1.**
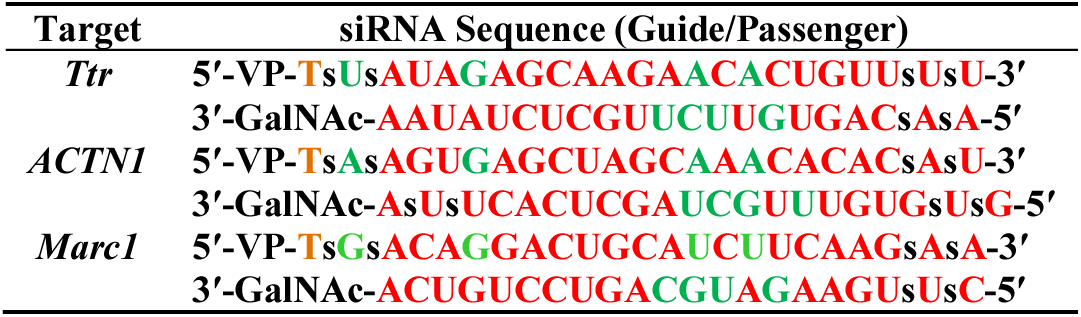
siRNA sequences and chemical modifications: red: 2′-O-methyl, green: 2′-fluoro, black: 2′-deoxy, orange: 2′-O-methoxyethyl; GalNAc: N-acetylgalactosamine; VP: 5′-(*E*)-vinylphosphonate; all linkages are phosphodiester unless marked with “s” (phosphorothioate).

We employed a Dual-Glo Luciferase Reporter Assay (Promega) to evaluate the relative effects of these modifications on both on-target and off-target silencing.^7, 13, 43, 44^ In this assay, cells were co-transfected with the modified siRNAs and two distinct reporter constructs: one containing a perfectly complementary site in the 3′ untranslated region (UTR) of Renilla luciferase to measure on-target activity, and another featuring four tandem seed-complementary sites to sensitively capture miRNA-like off-target effects. Firefly luciferase served as an internal normalization control, enabling reliable comparison within each experiment (see Experimental Section for details).

Our results revealed that both dx and aLd, when introduced at seed positions 6 and 7, consistently and substantially reduced off-target reporter repression for both *Ttr* (Figure 2) and *ACTN1* (Figure 3a-b) siRNAs, relative to the unmodified parent within each experiment. Notably, the greatest improvements were observed when aLd was incorporated at position 7 (hereafter aLd A7) and dx at position 6 (hereafter dx G6). This was evidenced by a decreased ability of the modified siRNAs to repress the off-target reporter, indicating improved specificity. These effects were highly reproducible across independent experiments. However, due to inherent variability between runs, the dual luciferase assay is best suited for assessing relative differences in silencing specificity within a single experiment, rather than for determining precise IC₅₀ values across separate assays. The observed trends should therefore be interpreted as consistent improvements in off-target discrimination rather than as quantitative measures of potency. Importantly, across all tested configurations and replicates, the modified siRNAs maintained on-target knockdown efficiency, exhibiting comparable levels of target mRNA suppression to their unmodified counterparts. This demonstrates that improved specificity was achieved without compromising the intended silencing activity.

**Figure 2.**
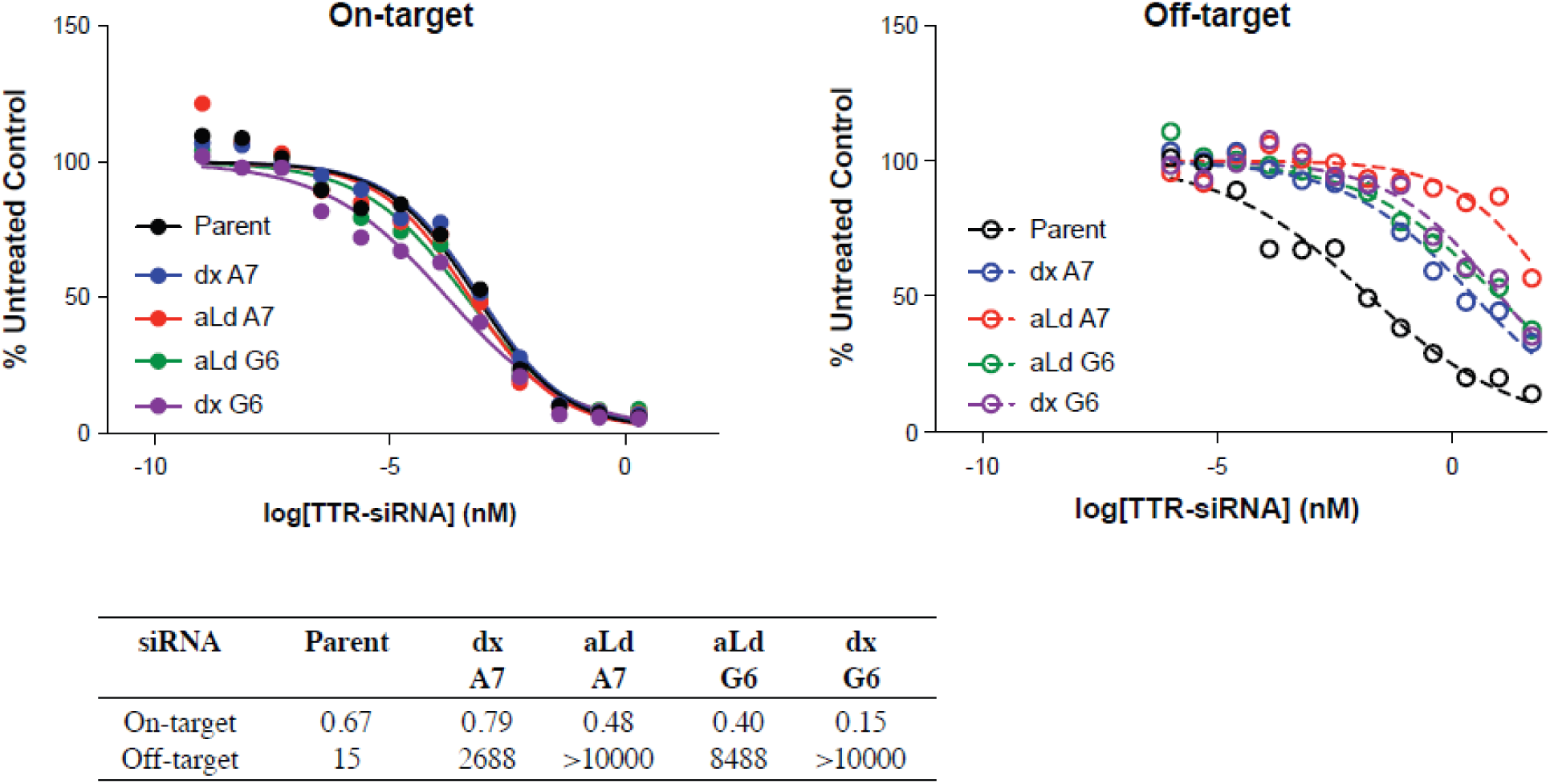
Dual-Luciferase Reporter Assay results for *Ttr* siRNAs modified with dx or aLd at positions 6 and 7 of the guide strand. Solid line: on-target; dashed line: off-target. IC_50_ values are reported in pM.

**Figure 3.**
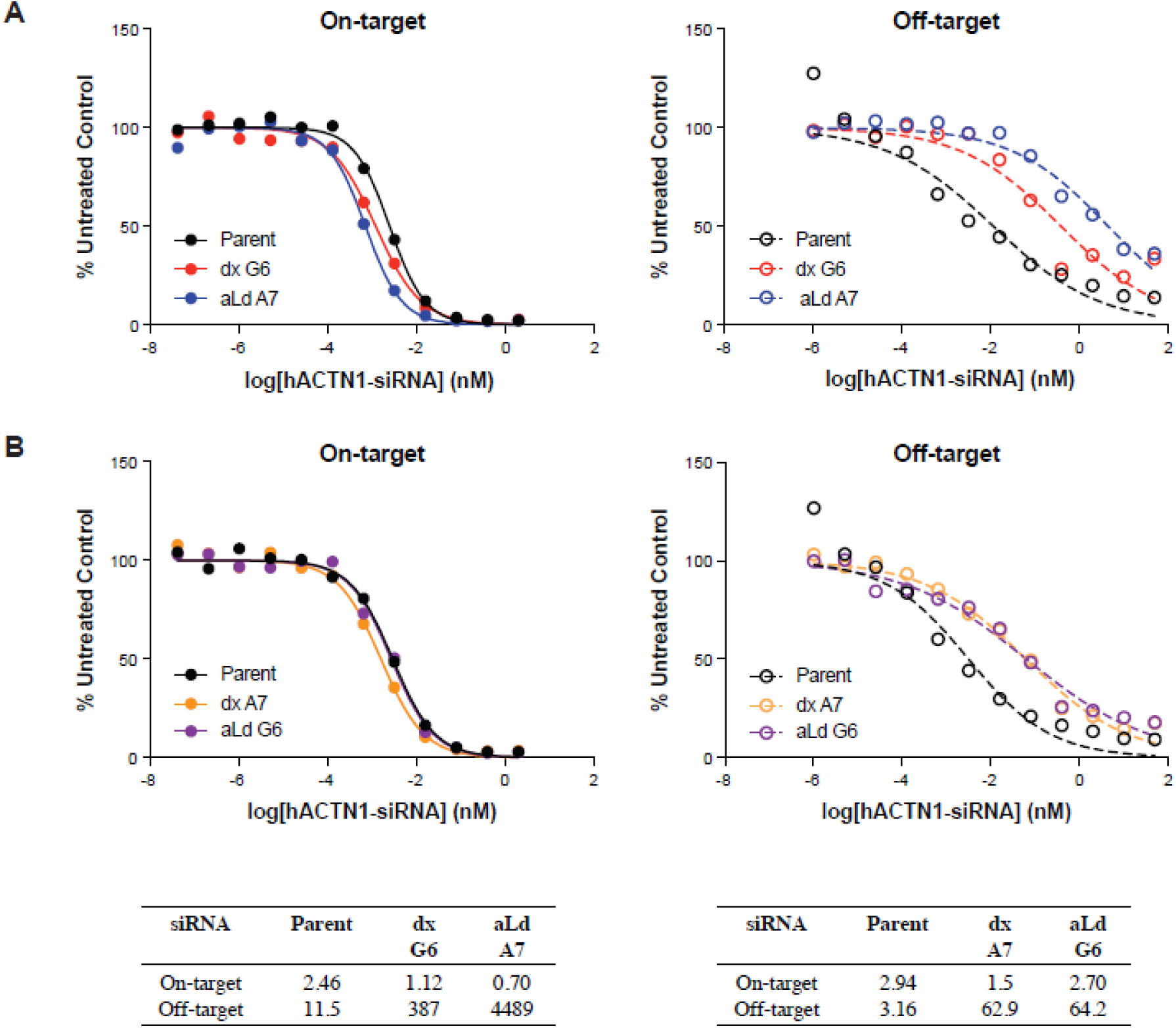
Dual-Luciferase Reporter Assay results for *ACTN1* siRNAs modified with dx or aLd at guide strand positions 6 and 7. (a) dx at position 6 (dx G6) and aLd at position 7 (aLd A7); (b) dx at position 7 (dx A7) and aLd at position 6 (aLd G6). Solid line: on-target; dashed line: off-target. IC₅₀ values are reported in pM.

Encouraged by these promising cell-based findings, we next sought to evaluate the performance of these modified siRNAs in a physiologically relevant context. To this end, we assessed the *in vivo* silencing efficacy of *Ttr*-targeting siRNAs containing dx or aLd at guide strand positions 6 and 7 in mice.

### *In vivo* functional and transcriptomic profiling of *Ttr* siRNAs modified with dx and aLd

One of the key considerations in developing siRNA therapeutics is balancing enhanced specificity with effective *in vivo* silencing efficacy. To assess whether sugar stereochemistry can help achieve this, we evaluated *Ttr*-targeting siRNAs containing dx or aLd at guide strand positions 6 and 7 (Table 1), sites within the seed region critical for both target engagement and off-target interactions. Modified and unmodified siRNAs were administered subcutaneously at doses ranging from 0.1 to 3 mg/kg. Hepatic *Ttr* mRNA levels were quantified by qPCR one week after dosing to assess gene silencing efficacy. All dx- and aLd-modified siRNAs retained robust *in vivo* potency, with ED₅₀ values approximately 1.1- to 1.8-fold higher than the unmodified parent sequence (Figure 4). These modest differences demonstrate that dx and aLd are well tolerated at central seed positions and preserve gene-silencing potency *in vivo*.

**Figure 4.**
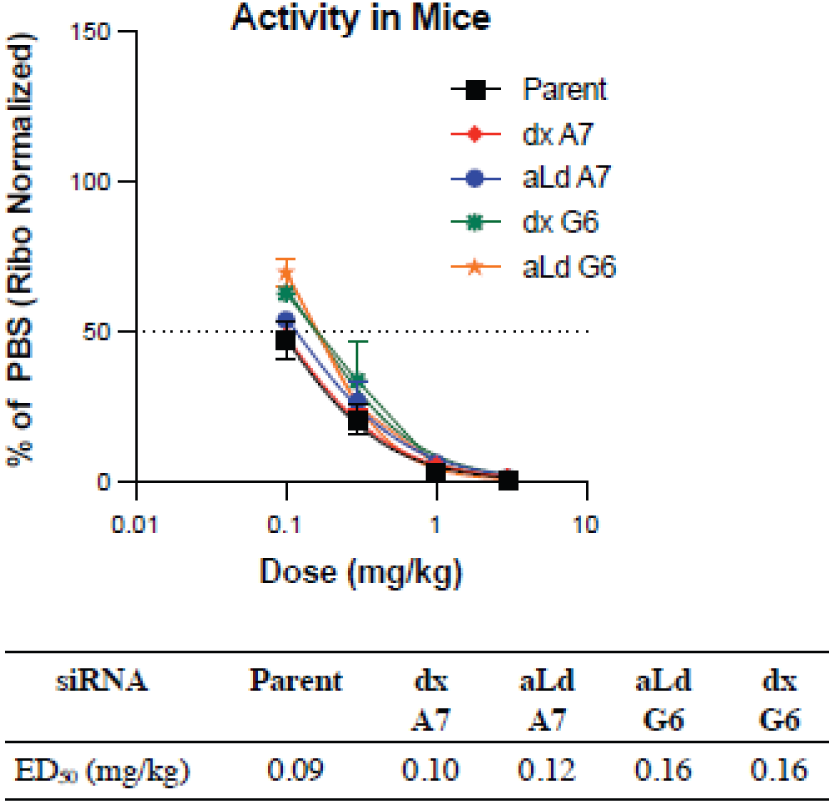
*In vivo* activities of the *Ttr* siRNAs modified with dx or aLd at guide strand positions 6 and 7 in mice, at doses of 3, 1, 0.3, or 0.1 mg/kg.

With *in vivo* potency established, we next evaluated whether these sugar modifications also mitigate transcriptome-wide off-target activity. To assess this, mice were dosed with the parent siRNA and the best-performing modified *Ttr*-targeting siRNAs identified in the dual luciferase screen: dx G6 and aLd A7. Liver tissues were collected one week after subcutaneous administration across an extended dose range (0.1 to 100 mg/kg), and transcriptomic profiling was performed. Differential gene expression (DGE) analysis was performed as outlined in the Materials and Methods, using criteria established previously to define differentially expressed genes (DEGs).^45^ Specifically, genes showing a fold change greater than twofold (either up or down), a p-value below 0.01, and a q-value under 0.1 were classified as the DEGs. This identification was made by comparing expression levels between saline-treated control mice and the siRNA-treated groups. Volcano plots (color coding defined in Figure 5 legend) were used to visualize the DEGs, where red points signify significant upregulation, blue points represent significant downregulation, and grey points indicate no substantial change in expression. The parent siRNA induced a dose-dependent increase in the DEGs. In contrast, both dx- and aLd-modified siRNAs showed a clear reduction in the DEGs across the entire dose range, particularly at higher concentrations (Figure 5), indicating effective suppression of off-target effects *in vivo*. In addition to reducing the DEGs, the modified siRNAs also decreased subthreshold gene expression changes, providing further evidence that sugar stereochemistry reduces unintended transcriptomic perturbations.

**Figure 5.**
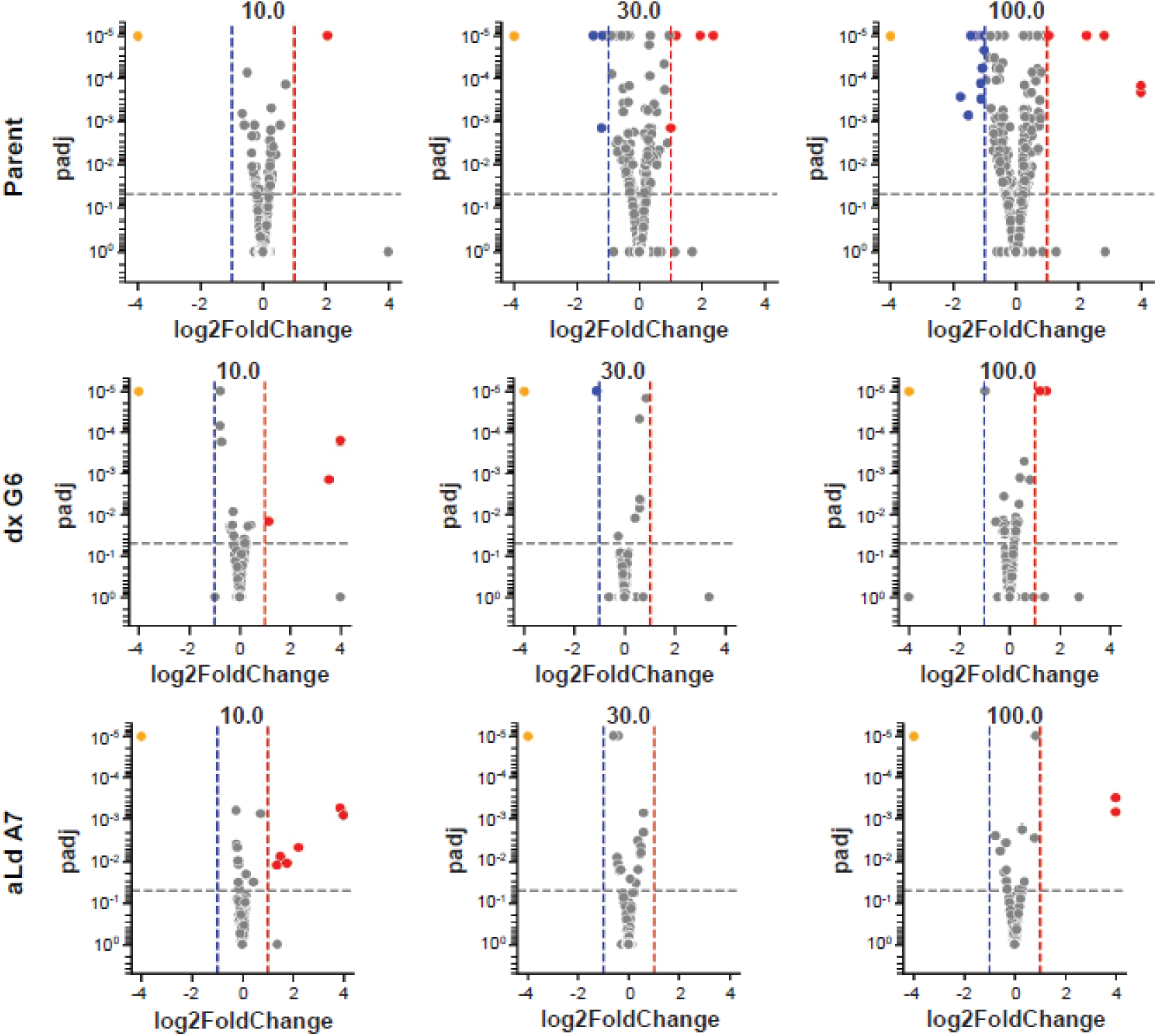
Volcano plots depicting DEGs in mouse liver after *Ttr* siRNA treatment at the top three doses (10, 30, and 100 mg/kg) over 7 days. Top row: parent siRNA; middle row: dx G6-modified siRNA; bottom row: aLd A7-modified siRNA. Colors indicate on-target (orange, *Ttr*), upregulated (red), downregulated (blue), and non-significant genes (grey).

These results demonstrate that dx and aLd modifications at central seed positions preserve potent on-target silencing while substantially improving *in vivo* specificity at the transcriptomic level. We next examined whether these improvements extend to a second siRNA sequence.

### *In vitro* functional and transcriptomic profiling of *ACTN1* siRNAs modified with dx and aLd

To further validate the findings from dual-luciferase reporter assays on how sugar stereochemistry influences siRNA specificity, we performed comprehensive *in vitro* functional and transcriptomic profiling using *ACTN1* siRNAs (Table 1) in A431 epidermoid carcinoma cells. Because *ACTN1* is a human-specific siRNA sequence, these experiments were conducted exclusively in human cells. The *ACTN1* siRNAs were modified with either dx G6 or aLd A7, as these positions had previously shown the most significant off-target mitigation in reporter assays. On-target silencing activity was assessed following a 96-hour incubation after transfection. Both modified siRNAs maintained comparable silencing, with IC₅₀ values closely matching those of the unmodified parent sequence, indicating that these sugar modifications preserved intended potency (Figure 6a). To evaluate transcriptome-wide off-target effects, DGE analysis was performed as outlined previously. Volcano plots were used to visualize the DEGs, using the color coding defined in the Figure 5 legend. The parent *ACTN1* siRNA induced a dose-dependent increase in the DEGs. In contrast, the dx G6 siRNA exhibited a clear reduction in the DEGs across the dose range, particularly at higher concentrations, indicating effective mitigation of off-target activity at the transcriptomic level (Figure 6b). A different pattern was observed for the aLd-modified *ACTN1* siRNA (aLd A7). Although this modification showed pronounced off-target suppression in the dual-luciferase assay, the DGE results revealed little to no reduction in the number of DEGs relative to the parent (Figure 6b). This discrepancy highlights a disconnect between the dual-luciferase assay and transcriptome-wide profiling for this modification, suggesting that certain sugar isomers may behave differently depending on the assay context or cellular environment. Together, these results indicate that dx and aLd modifications can differentially influence off-target activity depending on the sequence and experimental context.

**Figure 6.**
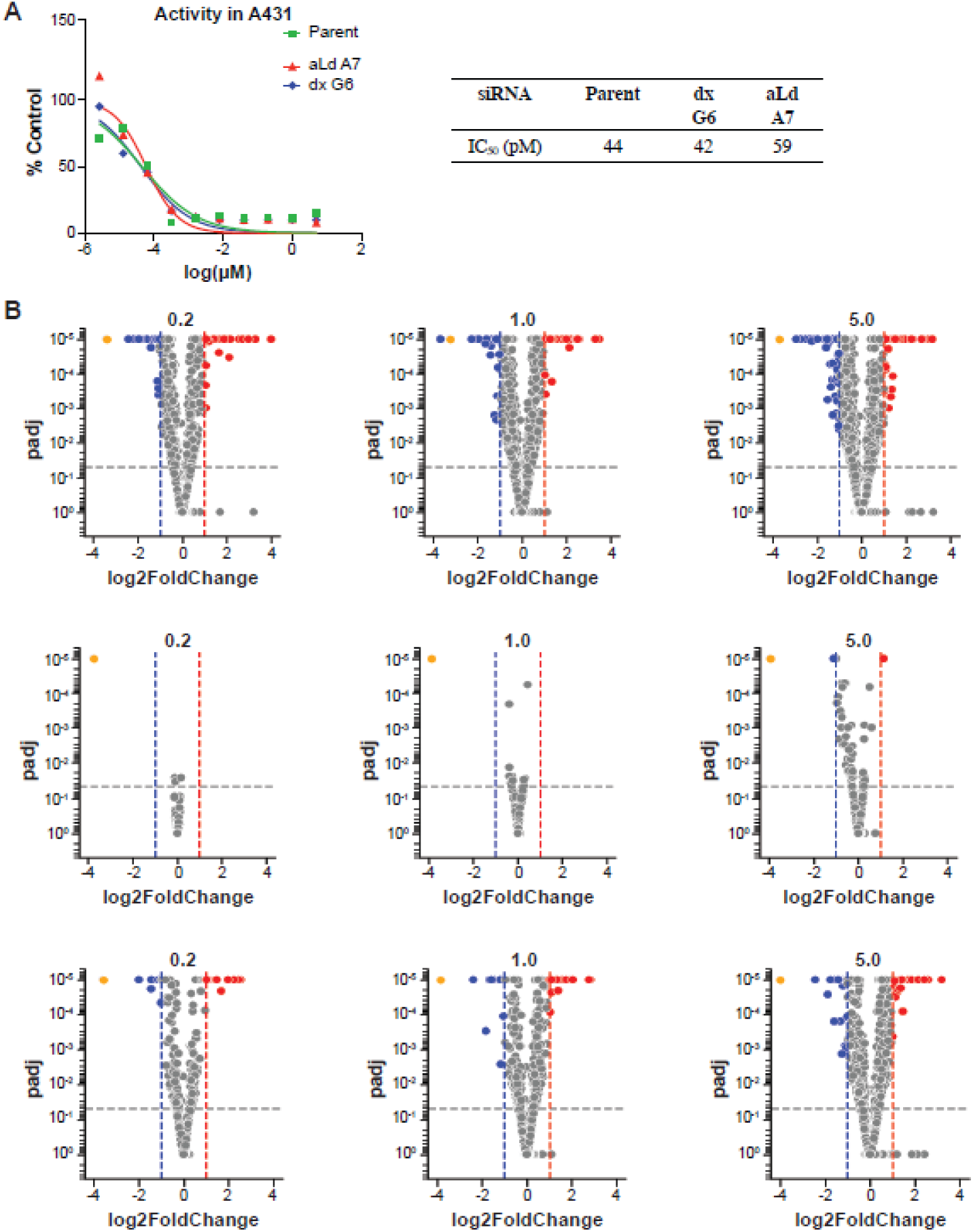
(a) *In vitro* activities of *ACTN1* siRNAs with dx or aLd modifications at guide positions 6 and 7 in A431 cells (96 h, lipid transfection). (b) Volcano plots of DEGs after *ACTN1* siRNA treatment (0.2, 1, 5 µM). Top row: parent siRNA; middle row: dx G6-modified siRNA; bottom row: aLd A7-modified siRNA. Colors indicate on-target (orange, *ACTN1*), upregulated (red), downregulated (blue), and non-significant genes (grey).

### Toxicity evaluation of *Marc1* siRNAs functionalized with dx and aLd

Building on the improved specificity observed for *Ttr* and *ACTN1* siRNAs, we next evaluated whether dx and aLd modifications could also mitigate sequence-driven toxicity in a physiologically relevant model. To test this, we applied the same modification strategy to a previously characterized *Marc1*-targeting siRNA that induces hepatotoxicity at submilligram doses and causes measurable liver toxicity within three days.^7^ The DGE study was therefore designed as a short-term assessment to capture early transcriptomic changes while minimizing secondary effects of toxicity. *Marc1* siRNAs were synthesized with dx or aLd at seed region positions 6 and 7, and their impact on off-target activity and toxicity was evaluated through two complementary *in vivo* studies using transcriptomic (DGE) and hepatotoxicity endpoints.

In the first experiment, *Marc1* siRNAs were administered at 0.4, 2, and 10 mg/kg in Balb/c mice (n = 4/group), and animals were sacrificed after three days for liver tissue analysis by DGE and qPCR. qPCR demonstrated dose-dependent on-target knockdown, with hepatic *Marc1* mRNA reduced by ∼80–90% at 2 and 10 mg/kg (Figure S1). Volcano plots summarizing the transcriptomic data (Figure 7a-b) showed that the parent siRNA induced widespread off-target changes in differentially expressed genes (DEGs), whereas the modified siRNAs, particularly dx G6 and aLd G6, substantially reduced the number of DEGs across all doses. Cumulative distribution function (CDF) analyses (Figure 7c) showed pronounced left-shifts for the parent siRNA (Δ ≈ –0.20 to –0.12) that were markedly reduced for the dx G6 and aLd G6 modifications (Δ ≈ –0.03 to –0.08), confirming mitigation of seed-mediated repression (see Supporting Information, RNA Sample Processing and DGE Analysis).

**Figure 7.**
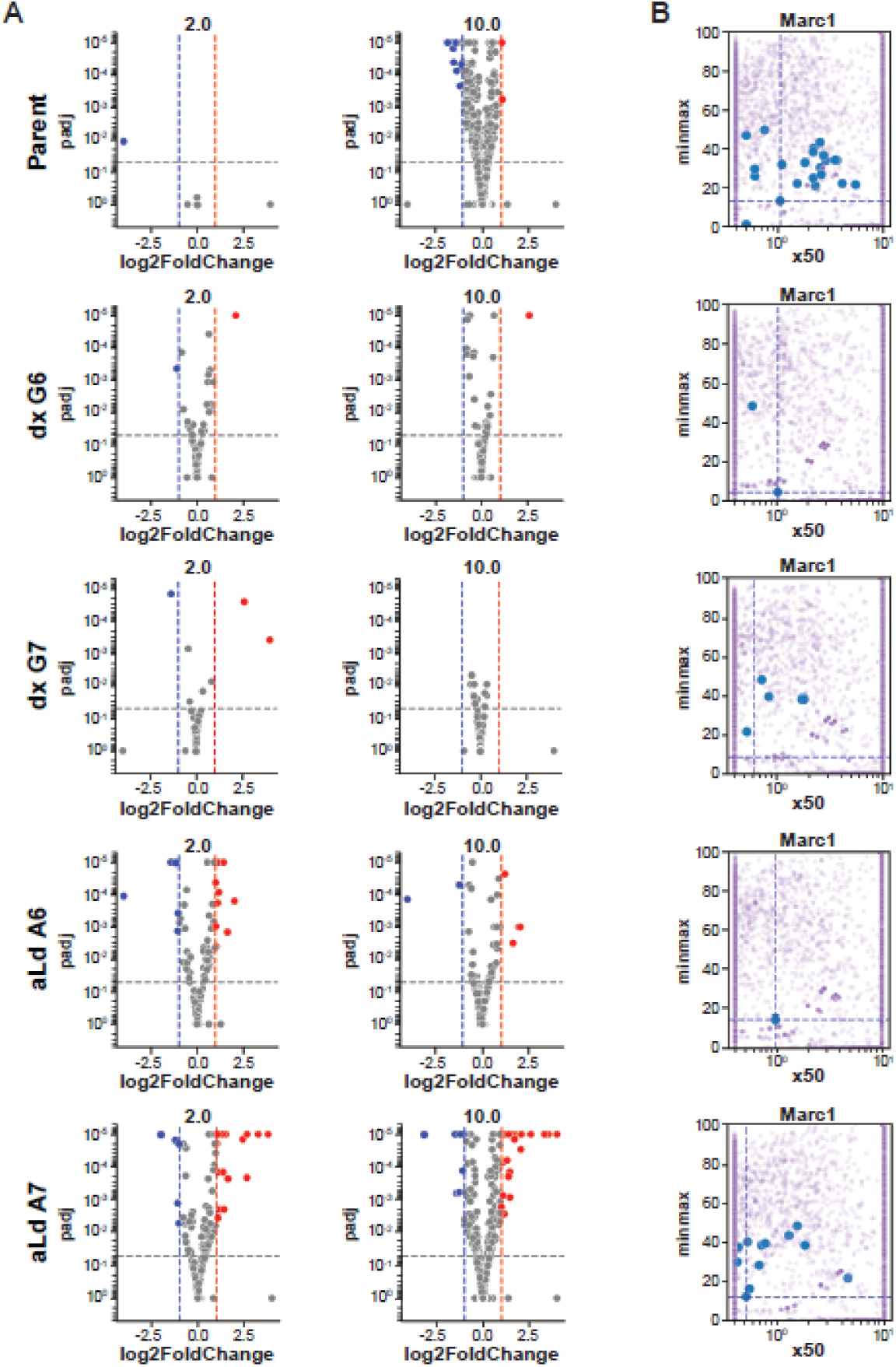
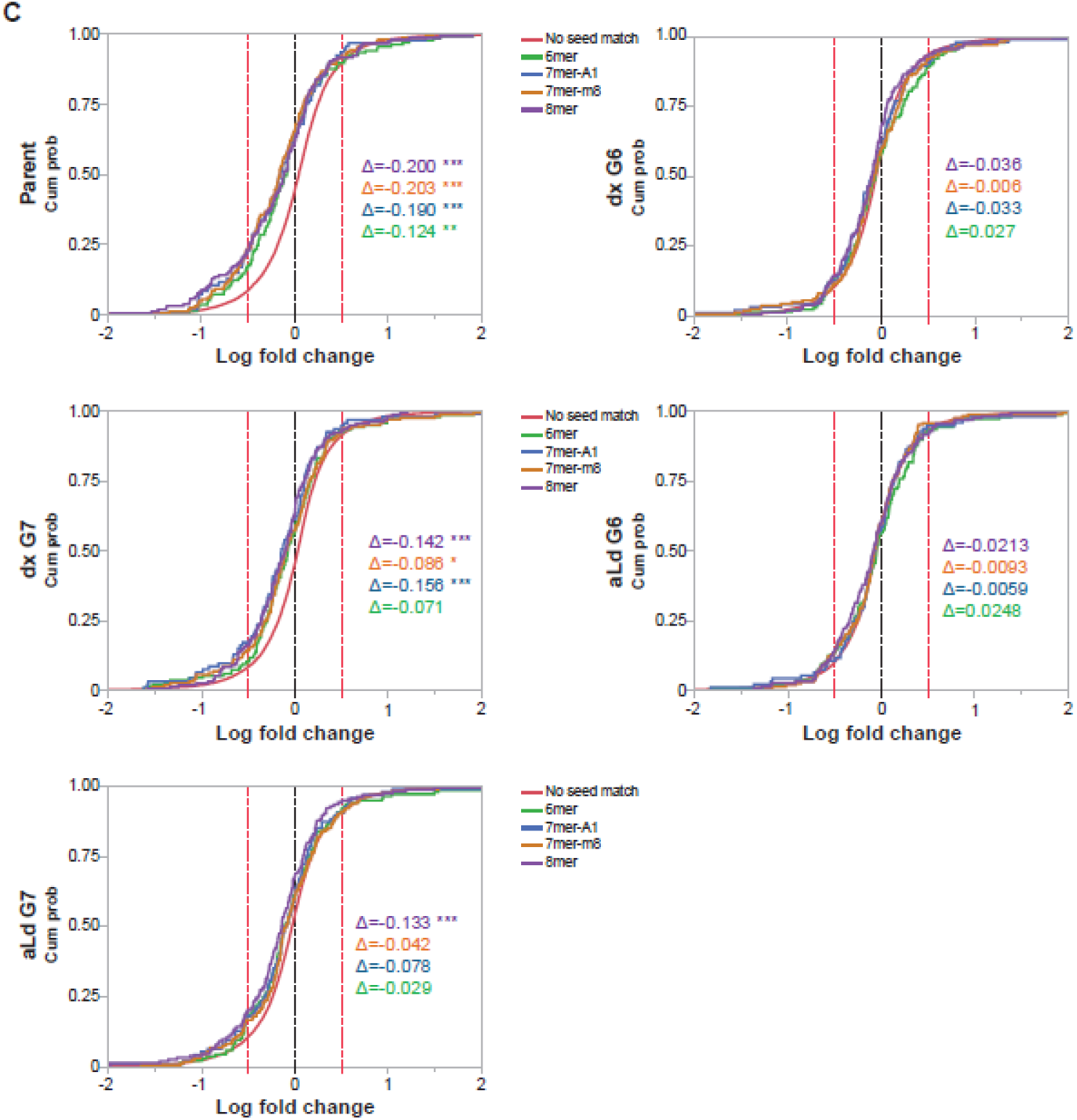
(a) Volcano plots depicting DEGs in mouse liver after *Marc1* siRNA treatment at the top two doses (2 and 10 mg/kg) over 3 days. From top to bottom, plots correspond to parent siRNA, dx G6, dx G7, aLd G6, and aLd A7 siRNAs. Colors indicate upregulated (red), downregulated (blue), and non-significant genes (grey). (b) Plot of relative IC_50_ values of down-regulated genes relative to on-target IC_50_ values. Colors indicate blue for genes exhibiting concentration-responsive knockdown and purple for genes with non-significant expression changes. (c) Comparative expression analysis of transcripts containing 8mer or 6/7mer seed matches versus those with no seed match following *Marc1* siRNA treatment at 10 mg/kg over a 3-day period. Delta values represent cumulative distribution function (CDF) shifts, where negative values indicate downregulation. Statistical significance is denoted as follows: *p < 0.05, **p < 0.01, ***p < 0.001; absence of asterisks indicates non-significant differences (p ≥ 0.05).

On-target knockdown efficiency was maintained across all modified siRNAs, yielding target mRNA suppression comparable to the unmodified parent (Figure S1). Notably, dx at position 7 enhanced on-target activity, lowering the ED_50_ from 0.985 mg/kg for the parent to 0.480 mg/kg (Figure S1). This reproducible trend was also observed across additional siRNA sequences evaluated during clinical candidate screening and is currently under investigation (data not shown). For the *Marc1* siRNA, these results suggest a modest trade-off between specificity and potency across different modification positions.

In a second *in vivo* study (Figure 8), mice received weekly subcutaneous doses of 10 mg/kg and were analyzed after one week for serum chemistry and liver histopathology. The parent siRNA caused severe hepatotoxicity, with ALT values reaching approximately 14,800 U/L, AST about 3,000 U/L, and GLDH greater than 600 U/L, accompanied by hepatomegaly (liver-to-body weight approximately 5.4%; data not shown). Histological examination revealed moderate single-cell necrosis, cytoplasmic degeneration, mild fibrosis, and inflammatory infiltrates. In contrast, dx G6 and aLd G6 completely normalized serum enzyme levels (ALT less than 100 U/L, AST less than 70 U/L, GLDH about 5 U/L) and showed no detectable histopathological alterations, indistinguishable from PBS controls. dx G7 exhibited partial improvement (ALT about 1,300 U/L, GLDH about 250 U/L) with minimal hepatocyte necrosis and mild fibrosis, while aLd G7 displayed near-baseline enzyme levels and only minimal centrilobular hypertrophy without degenerative findings. Liver-weight changes correlated closely with biochemical recovery, reinforcing the link between sequence-driven off-target activity and hepatocellular injury.

**Figure 8.**
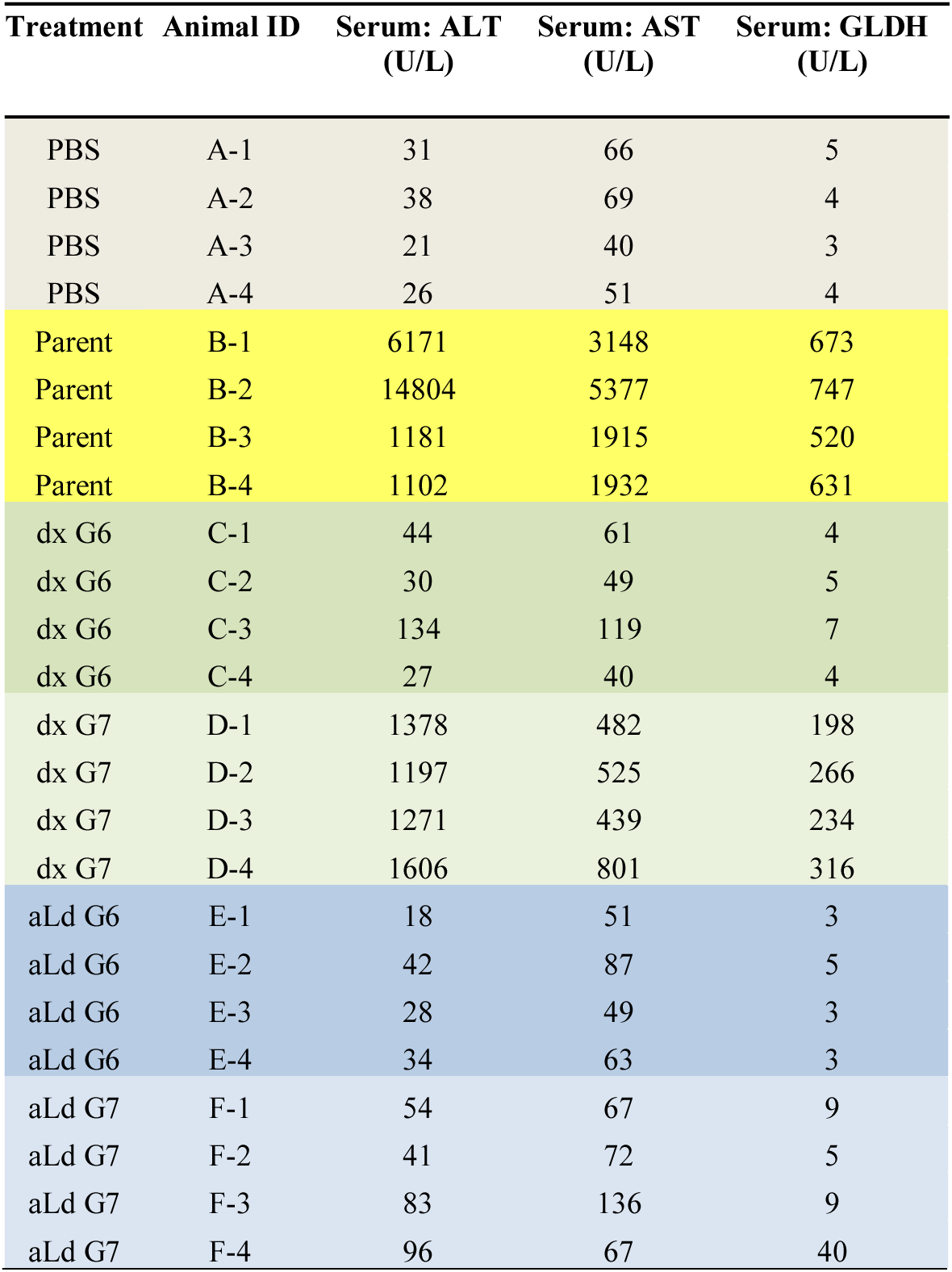
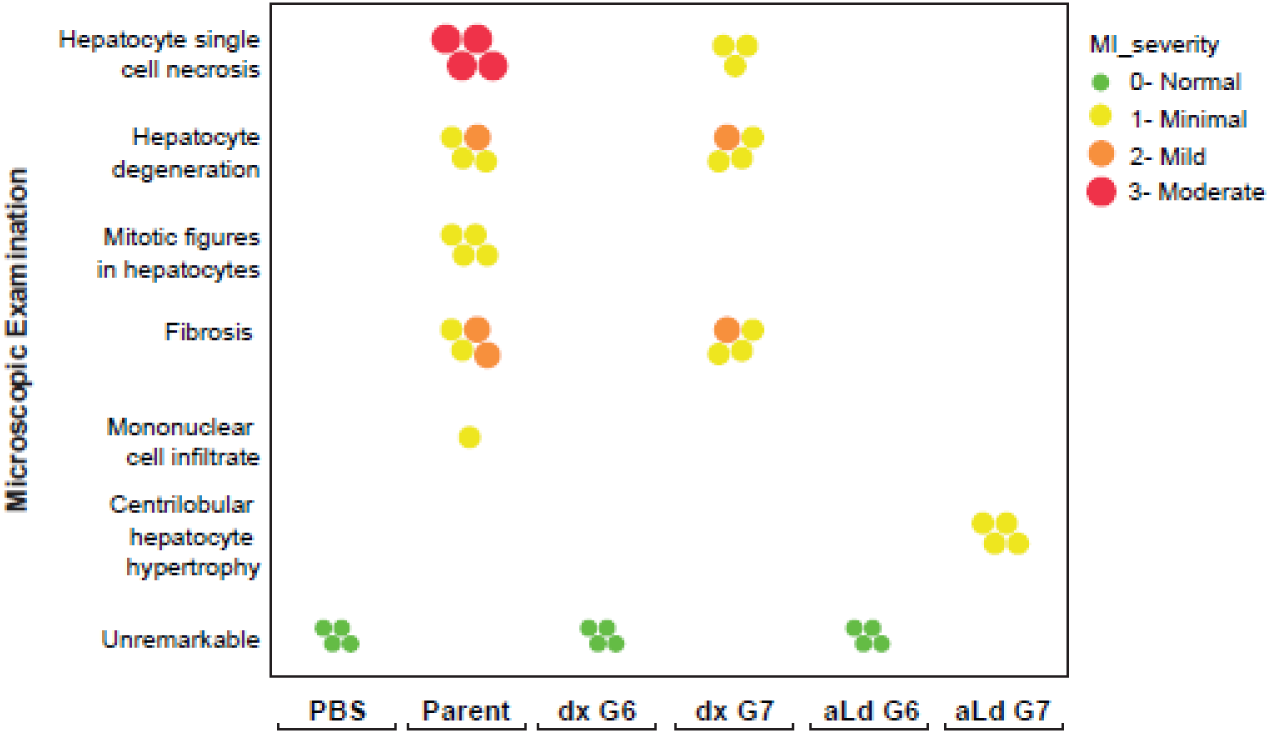
Toxicity evaluation of parent and modified *Marc1* siRNAs in mice. Animals received a single 10 mg/kg subcutaneous dose, and liver enzyme levels along with histopathology were assessed one week later. (a) The parent compound induced marked increases in serum liver enzymes, accompanied by (b) moderate hepatocellular necrosis, degeneration, and proinflammatory findings including mononuclear infiltrates and sinusoidal fibrosis. In contrast, the dx G6 and aLd G6 siRNAs produced no detectable enzyme elevations or pathological alterations.

### Molecular dynamics (MD) simulations

To explore how stereoisomeric sugar modifications influence RNAi specificity, molecular dynamics (MD) simulations were performed on *Ttr* siRNA guide strands incorporating dx and aLd at seed positions 6 or 7. These modified guide strands were modeled in complex with hAgo2 to assess their structural and energetic effects on guide-target duplex formation and interactions with the protein. Both dx and aLd analogs were stably accommodated within the hAgo2 binding pocket, preserving canonical Watson-Crick base pairing with the complementary target RNA throughout the simulations. The guide-target seed duplex maintained an A-form-like geometry, with the modified nucleotides adopting C3′-endo sugar puckers (average pseudorotation angles of 21°-25°). Notably, this conformation contrasts with the intrinsic preferences of the free sugars, indicating that hAgo2’s binding environment imposes structural constraints that override their natural sugar flexibility.

Structural studies by MacRae et al. identified a pronounced kink between nucleotides 6 and 7 in the seed region of the guide strand in hAgo2, created by the intercalation of Ile365 from α-helix 7, which disrupts A-form stacking in the unbound guide RNA.^46^ Target binding triggers a ∼4 Å shift of helix 7 into the minor groove, relieving the kink and restoring continuous A-form geometry beyond nucleotide 5. This movement stabilizes base pairing at the seed-central junction and allows the duplex to extend further along the target. Helix-7 functions as a dynamic structural element in this process: in the absence of target pairing, it inserts between nucleotides 6 and 7, breaking base stacking to create the kink; upon seed pairing, it pivots to dock into the minor groove, relieving the kink and stabilizing the duplex.^46^ This repositioning not only promotes extension of the guide-target pairing but also accelerates both target binding and release, the latter being critical for minimizing dwell time on partially matched targets and enabling rapid off-target dismissal.^47^ Importantly, α-helix 7 also serves as a molecular gatekeeper, actively favoring canonical Watson-Crick base pairs in the seed region while discriminating against non-canonical interactions including G:U wobbles and mismatches at nucleotides 6 and 7. This property enables hAgo2 to rapidly dismiss off-targets and maintain high specificity during the cellular target search. The severity of the 6-7 kink correlates with the extent of guide-target pairing, acting as an internal sensor for duplex fidelity and forming a central part of Argonaute’s target interrogation mechanism.^47^ Recent cryo-EM and kinetic data reveal that the kink also plays a second, catalytic role: by repositioning the nucleotide 6 phosphate, it opens space for Lys709 to swing into the active site alongside Asp669 to engage the scissile phosphate at t11. Without this geometry, K709 stays out of position and Ago2-mediated cleavage is impaired.^48^ The 6-7 kink therefore acts as a dual-function element, coordinating catalytic activation and stabilizing the guide-target duplex upon target binding.

In our simulations, dx at position 6 maintained the native phosphate-backbone interactions with Lys709 and Arg761, preserving the electrostatic network of the unmodified guide-protein interface (Figure 9). In functional assays with *Ttr* siRNAs, this placement produced the greatest improvement in specificity, maintaining strong knockdown activity while substantially mitigating unintended repression. MD analysis indicated that while kink geometry was preserved in perfect matches, imperfect duplexes exhibited modest increases in 6-7 backbone angular variability, consistent with reduced stability of mismatched seeds. These subtle changes in geometry may enhance helix-7’s ability to disengage from imperfect targets, improving mismatch discrimination without compromising on-target catalysis. This increased angular variability is consistent with helix-7’s role in minimizing dwell time on off-targets, allowing hAgo2 to rapidly bind and dissociate from imperfectly paired RNAs. In this way, the modification directly exploits helix-7’s discriminatory power to accelerate the rejection of off-targets and improve siRNA specificity. aLd at position 6 formed new attractive electrostatic contacts with Lys709 and Ser798, replacing interactions with Gln757, Thr759, and Ser760 seen in the unmodified guide strand (Figure 9). Because nucleotide 6 lies in the kink, these contacts stabilized the licensed geometry in perfect matches, promoting α-helix 7 movement and K709 activation, but in imperfect seeds produced little conformational change, which may explain its somewhat lower discrimination compared to dx at the same position, despite preserved activity. At position 7, which forms the downstream arm of the kink, dx established new interactions with Arg370 but produced minimal changes in kink geometry or local backbone conformation in either duplex context; its off-target suppression was more modest than the best-performing placements (Figure 9). In contrast, aLd at position 7 formed additional contacts with Thr759, altering phosphate positioning and backbone conformation more noticeably in imperfect seeds than in perfect matches. This remodeling likely sharpens helix-7’s sensitivity to mismatches in the 3′ end of the seed region, making it more effective at rejecting imperfect duplexes while maintaining high-fidelity binding to perfect matches (Figure 9).

**Figure 9.**
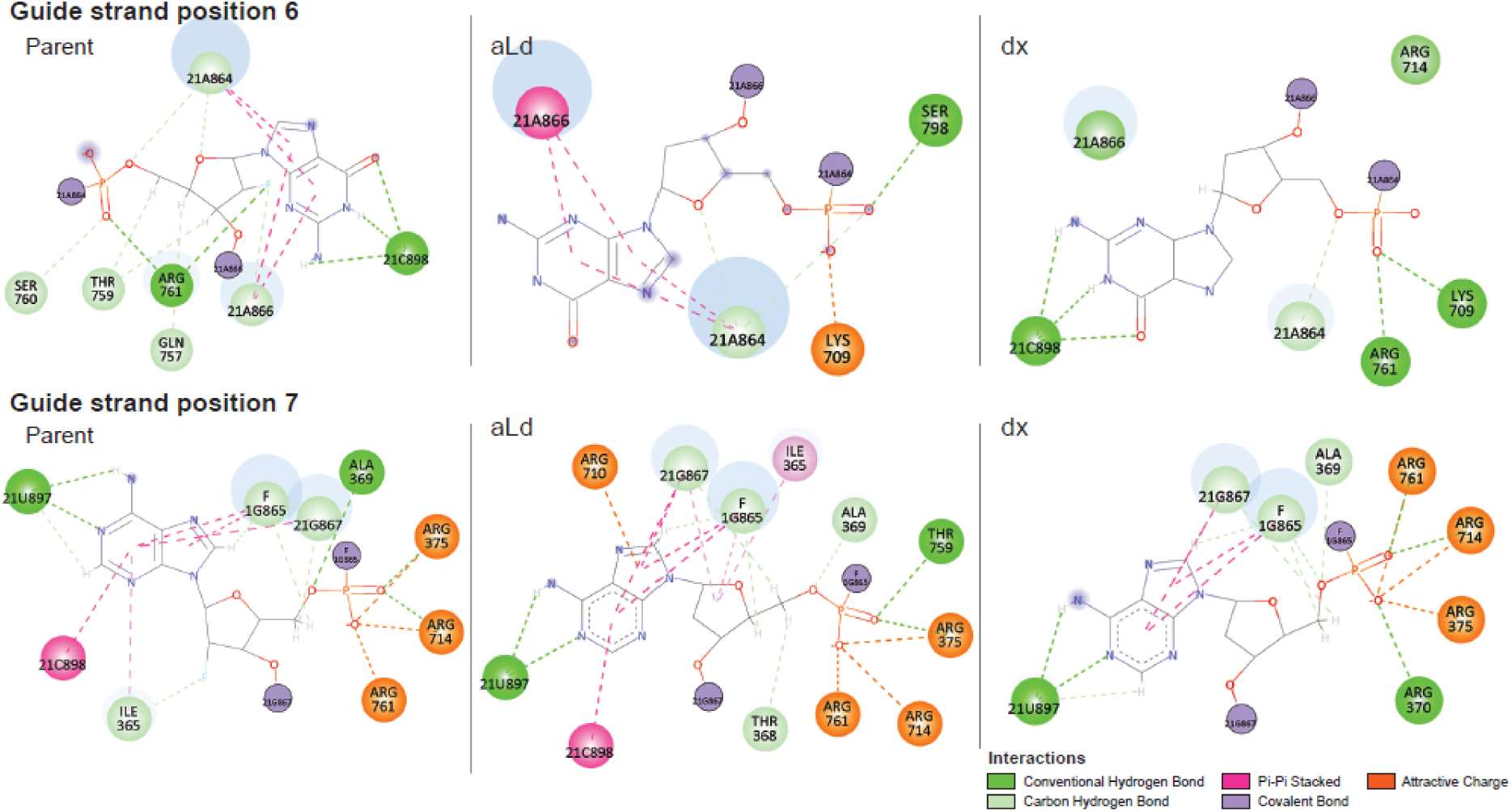
Interaction network of dx and aLd at positions 6 and 7 of the *Ttr* guide within hAgo2. Representative structure from the most populated cluster of ∼15 µs MD simulations showing key contacts between modified sugars and hAgo2 residues.

Collectively, these results indicate that dx at position 6 of *Ttr* siRNA is optimal for preserving native kink geometry in perfect matches while subtly destabilizing imperfect seeds, and that aLd at position 7 of *Ttr* siRNA excels by remodeling downstream kink geometry to penalize imperfect seeds without impairing on-target licensing. Both leverage the kink’s dual role in stabilization and catalysis to enhance specificity while maintaining efficacy and safety, with position and stereochemistry determining the balance between these effects. Notably, these findings underscore how helix-7’s dual function as both a catalytic activator and a molecular discriminator can be harnessed by local sugar modifications to maximize siRNA efficacy and specificity, facilitating rapid, accurate engagement with fully matched targets and swift rejection of imperfect duplexes in the complex cellular environment.

## CONCLUSIONS

In summary, this study reports the synthesis and systematic evaluation of all eight stereoisomeric phosphoramidites of β-D-2′-deoxyxylonucleoside (dx) and α-L-2′-deoxyribonucleoside (aLd), which were incorporated into the seed region of siRNAs to assess their impact on off-target activity, on-target efficacy, and safety. Using *Ttr* and *ACTN1* siRNAs with validated off-targets, dual-luciferase reporter assays demonstrated that introducing dx or aLd at guide strand positions 6 or 7 significantly reduced off-target effects. These findings were substantiated by transcriptome-wide differential gene expression (DGE) analyses performed *in vitro* and *in vivo*, which revealed a consistent decrease in unintended gene modulation without compromising silencing potency.

These results highlight the necessity of employing complementary assay platforms to provide a comprehensive evaluation of off-target activity, reflecting the influence of sequence context and cellular environment. Such variability emphasizes that chemical modifications should be carefully optimized for each siRNA to maximize efficacy and specificity.

Importantly, *in vivo* toxicity studies revealed that *Marc1*-targeting siRNAs containing dx or aLd modifications exhibited reduced hepatotoxicity, as indicated by lower serum ALT and AST levels, demonstrating an improved safety profile. Molecular dynamics simulations provided mechanistic insight, showing that dx and aLd sugars adopt a C3′-endo sugar pucker compatible with the hAgo2 seed-binding pocket. These modifications subtly alter local electrostatic interactions, which are proposed to modulate conformational dynamics and binding kinetics, thereby enhancing discrimination against partially complementary off-target sequences while maintaining on-target engagement. In addition to preserving potency, dx at position 7 enhanced on-target activity (Figure S1). This reproducible trend was also observed across multiple siRNAs during internal candidate screening, and we are now investigating its underlying mechanism.

Together, these results establish stereochemically engineered sugar modifications as valuable tools for fine-tuning siRNA specificity and safety. However, the breadth of this study also highlights the importance of evaluating these modifications across a large and diverse panel of siRNAs to fully characterize their performance and generalizability. Notably, a key strength of the dx and aLd modifications identified here is their ability to preserve potent on-target silencing comparable to unmodified siRNAs, which is a vital consideration for therapeutic translation.

In closing, this work exemplifies how integrating systematic chemical synthesis, multifaceted functional assays, *in vivo* validation, and molecular modeling can drive the rational design of next-generation siRNA therapeutics. By precisely tailoring the molecular ‘key,’ these modifications enable siRNAs to engage selectively with their intended mRNA ‘lock,’ while minimizing interactions with off-target sequences

## EXPERIMENTAL SECTION

### General Information

All reagents and solvents were obtained from commercial suppliers and used as received unless otherwise noted. General synthetic and analytical procedures (TLC, flash chromatography, NMR, LC-MS) followed previously reported methods.^7^ Complete synthetic, analytical, and biological details are provided in the Supporting Information.

### Synthesis of dx and aLd phosphoramidites

The dx and aLd phosphoramidites of uridine, cytidine, adenosine, and guanosine were synthesized according to the procedures described in the Supporting Information.

### Oligonucleotide Synthesis

Guide and passenger strands were synthesized on an Applied Biosystems 394 DNA/RNA Synthesizer (2 µmol scale) and on an ÄKTA OligoPilot 10 (40 µmol scale) using VIMAD UnyLinker or pre-loaded GalNAc solid supports. Purification by ion-pair reversed-phase and strong-anion-exchange chromatography was performed as previously described.^7^ Full experimental details are provided in the Supporting Information.

### Biological Assays

Dual-luciferase reporter assays, A431 cell-culture experiments, *in vivo* studies of *Ttr* and *Marc1* siRNAs, and histological analyses were performed as previously reported,^7^ unless otherwise indicated. Concentrations, dosing regimens, primer/probe sequences, and statistical analyses are described in the Supporting Information.

### Differential Gene-Expression (DGE) and Data Analysis

DGE profiling was carried out using QuantSeq 3′ mRNA-Seq on the Illumina platform following our established procedures.^7, 45^ Threshold criteria, software parameters, and the full computational workflow are detailed in the Supporting Information.

### Molecular Dynamics Simulations

Modified residues of the *Ttr* sequence (vinyl phosphonate T, 2′-fluoro nucleosides, 2′-O-methyl nucleosides, and the aLd and dx analogs) were built using the modXNA methodology^49^ within hAgo2 (PDB 6N4O). Simulation setup, force-field parameters, and trajectory analyses are described in the Supporting Information.

## ASSOCIATED CONTENT

Supplementary data are available with this preprint.

## AUTHOR INFORMATION

### Author Contributions

Organic syntheses: G.V., G.C.F., M.T.M.; Oligonucleotide syntheses: M.N., M.A.; Dual-luciferase reporter assays: M.T.; DGE experiments: S.D.; Cell culture assays: C.Q., A.T.W.; Animal studies: A.L.; Toxicity studies: H.L., C.Y.H., D.T.T.; Histology: S.K.; Molecular dynamics simulations: R.G.-M.; Supervision: T.P.P., E.E.S.; Writing and project management: M.N.; Review and editing: all authors.

### Notes

All authors are employees of Ionis Pharmaceuticals, Inc.

## ACKNOWLEDGMENT

We thank Dr. Hans Gaus for acquiring high-resolution LC–MS data and Raul Alonzo for assistance with the graphics.

## ABBREVIATIONS

Dx: β-D-2′-deoxyxylonucleoside
aLd: α-L-2′-deoxyribonucleoside
VP: vinylphosphonate
SAX: strong anion exchange
*Ttr*: transthyretin
*ACTN1*: alpha-actinin 1
*Marc1*: mitochondrial amidoxime reducing component 1
DGE: differential gene expression
hAgo2: human Argonaute 2
ALT: alanine aminotransferase
AST: aspartate aminotransferase

## DATA AVAILABILITY

The differential gene expression (DGE) data generated in this study have been deposited in the NCBI Gene Expression Omnibus (GEO) under accession number GSE313756. All other data supporting the findings of this study are available within the article.

