## Supplementary Data for "Tuning siRNA Specificity through Seed Region Incorporation of Deoxyribonucleotide Stereoisomers"

**List of contents**

| | |
|---|---|
| Synthesis | S2-S51 |
| Methods | S52-S57 |
| Table S1 | S57 |
| Figure S1 | S58 |
| Figure S2 | S58 |
| References | S59-S60 |
| NMR Spectra | S61-S92 |

**Synthesis of aLd nucleosides:** The systematic synthesis of the free, unprotected aLd nucleosides of uracil/thymine (U/T), cytosine (C), adenine (A), and guanine (G) was first documented in 1991 (shown below).[1] These early synthetic routes employed classical glycosylation and deoxygenation procedures for the preparation of A, T, G, and U nucleosides, while site-specific chemical modifications were applied for cytosine derivatives.

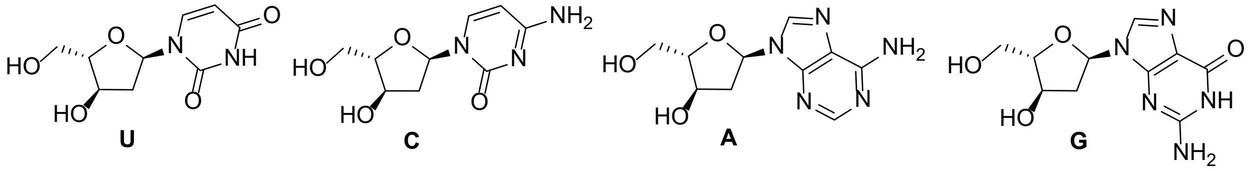


To the best of our knowledge, the synthesis reported herein represents the first unified and comprehensive route to the corresponding protected phosphoramidites, with extensive characterization of final compounds and key intermediates. The foundational prior work underlying these synthetic strategies has been appropriately cited at the beginning of each respective section.

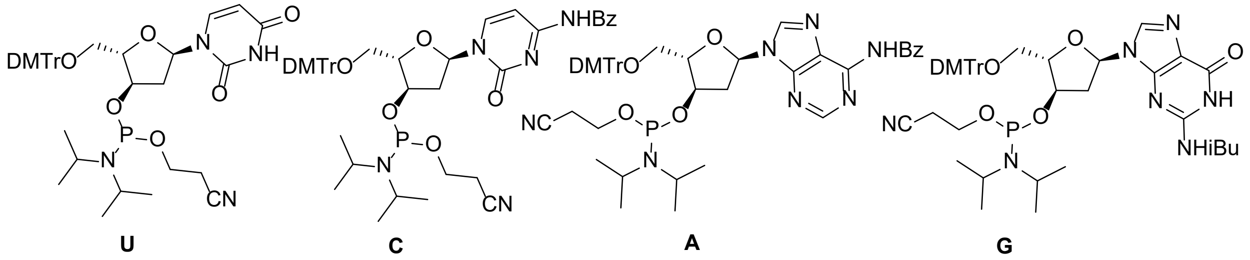


**Synthesis of aLd U phosphoramidite.** The syntheses of aLd T and aLd U phosphoramidites have been reported previously,[2, 3] with the aLd U phosphoramidite described in a patent.[3] Here, we follow similar procedures with modifications and provide full characterization of the final nucleosides and phosphoramidites, as well as most intermediates, several of which lacked analytical data in prior reports

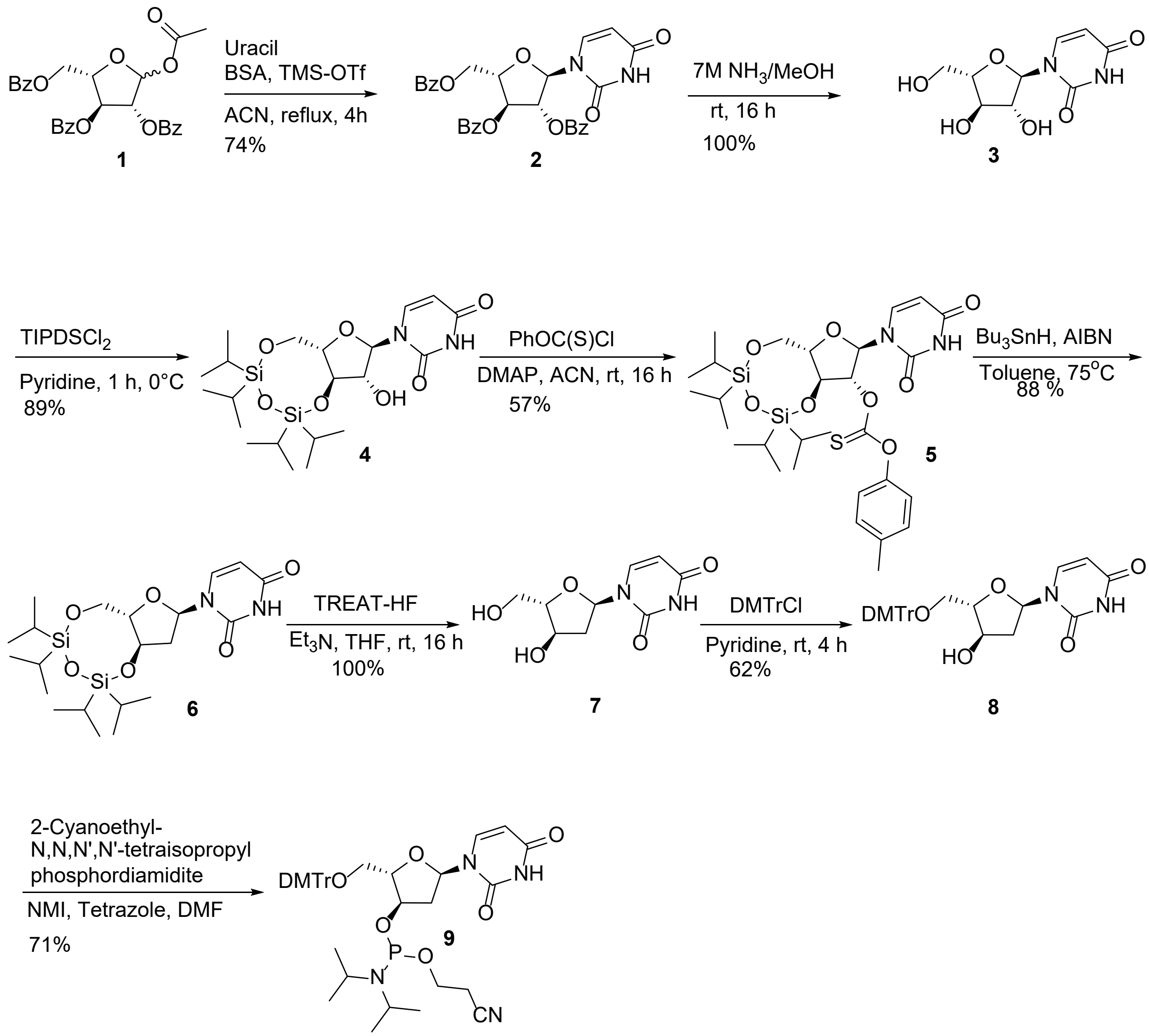


**(2S,3S,4R,5R)-2-((Benzoyloxy)methyl)-5-(2,4-dioxo-3,4-dihydropyrimidin-1(2H)-yl)tetra hydrofuran-3,4-diyl dibenzoate *(2)***

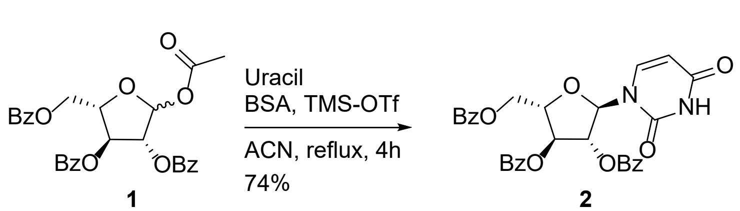


Uracil (47.10 g, 420.23 mmol) and N,O-bis(trimethylsilyl)acetamide (205.50 mL, 840.45 mmol) were added to a solution of **1**[2, 4] in acetonitrile (800 mL). The mixture was heated at 40 °C for 30 min to obtain a clear solution, then cooled in an ice bath. Trimethylsilyl trifluoromethanesulfonate

(74.72 mL, 336.18 mmol) was added, and the reaction was stirred at 80 °C for 4 h. The mixture was concentrated under reduced pressure and diluted with ethyl acetate. The organic layer was washed successively with saturated sodium bicarbonate solution and brine, dried, and concentrated to an oil. Purification by silica gel chromatography (Si, 1.5 kg column, 0–50 % ethyl acetate/hexanes) afforded the desired product as a white solid.

**Yield:** 87 g (156.33 mmol, 74%). **$^1$H NMR** (300 MHz, DMSO-$d_6$) δ 11.51 (s, 1H), 7.92–8.07 (m, 7H), 6.26 (d, J = 4.04 Hz, 1H), 6.08 (t, J = 4.08 Hz, 1H), 5.84–5.97 (m, 1H), 5.71 (d, J = 7.99 Hz, 1H), 5.10 (q, J = 4.61 Hz, 1H), 4.66 (d, J = 4.49 Hz, 2H). **$^{13}$C NMR** (75 MHz, DMSO-$d_6$) δ 165.9, 165.5, 165.3, 163.7, 151.1, 142.8, 134.4, 134.3, 134.0, 130.0, 129.7, 129.3, 129.2, 129.0, 102.3, 90.9, 81.9, 80.1, 77.1, 64.6. **LC-MS:** m/z 555.1 $[M–H]^-$.

**1-((2R,3R,4R,5S)-3,4-Dihydroxy-5-(hydroxymethyl)tetrahydrofuran-2-yl)pyrimidine-2,4 (1H,3H)-dione *(3)***

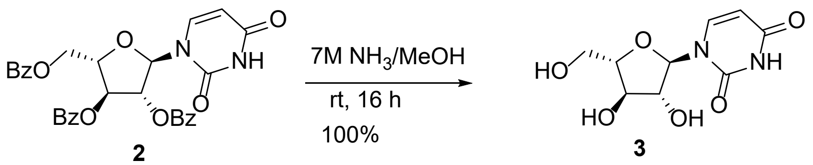


Ammonia in methanol (7.00 M, 150 mL) was added to a solution of **2** (87 g, 156.43 mmol) in methanol (80 mL). The reaction mixture was heated at 40 °C for 16 h and then stirred at rt for 72 h. The solvent was evaporated under reduced pressure to yield an oil, which was purified by Biotage chromatography (Si, 220 g column, 0–20 % methanol/dichloromethane) to afford the desired product as a white solid.

**Yield:** 38 g (155.69 mmol, 100 %). **$^1$H NMR** (300 MHz, DMSO-$d_6$) δ 11.26 (br s, 1H), 7.70 (br d, J = 7.72 Hz, 1H), 5.58–5.75 (m, 3H), 5.41 (br s, 1H), 4.92 (br s, 1H), 4.09 (br s, 2H), 3.94 (br s,

1H), 3.32–3.60 (m, 2H). $^{13}$**C NMR** (75 MHz, DMSO-d$_6$) δ 163.7, 151.1, 142.0, 101.7, 91.0, 87.5, 80.1, 75.6, 61.7. **LC-MS:** m/z 245.1 [M+H]$^+$.

**1-((6aS,8R,9R,9aR)-9-hydroxy-2,2,4,4-tetraisopropyltetrahydro-6H-furo[3,2-f][1,3,5,2,4]trioxadisilocin-8-yl)pyrimidine-2,4(1H,3H)-dione** ***(4)***

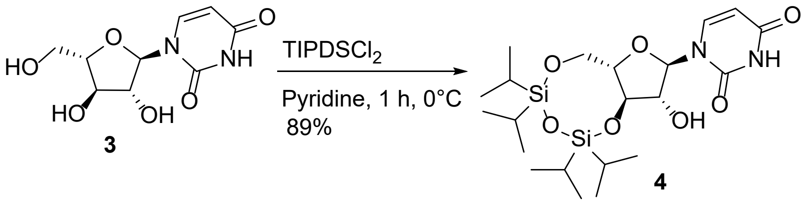


Compound **3** (56.24 g, 230.30 mmol) was dissolved in pyridine (100 mL) and evaporated to dryness under reduced pressure at 60 °C three times to dry the starting material. The residue was dissolved again in pyridine (700 mL), and 1,3-dichloro-1,1,3,3-tetraisopropyldisiloxane (69.99 mL, 218.79 mmol) was added at 0 °C under ice-bath cooling. The reaction mixture was stirred for 1 h, quenched with water, and concentrated under reduced pressure to an oil. The oil was dissolved in ethyl acetate, and the organic layer was washed sequentially with 10 % aqueous HCl, water, saturated sodium bicarbonate solution, water, and brine. The combined organic layers were dried and concentrated under reduced pressure to afford a crude oil. Purification by Biotage chromatography (Si, 220 g column, 0–60 % ethyl acetate/hexanes) afforded the desired product as a white solid.

**Yield:** 99.80 g (205.05 mmol, 89 %). $^{1}$**H NMR** (300 MHz, DMSO-d$_6$) δ 11.38 (br d, J = 9.96 Hz, 1H), 7.81–8.00 (m, 1H), 5.82 (br dd, J = 5.83, 10.59 Hz, 1H), 5.57–5.73 (m, 2H), 4.37–4.60 (m, 1H), 4.02–4.34 (m, 2H), 3.88 (br s, 2H), 3.28–3.44 (m, 2H), 0.92–1.21 (m, 25H). $^{13}$**C NMR** (75 MHz, DMSO-d$_6$) δ 163.6, 151.2, 143.0, 102.4, 89.0, 81.6, 77.0, 75.3, 61.9, 17.8, 17.6, 17.4, 17.3, 13.3, 13.0, 12.6, 12.5. **LC-MS:** m/z 485.2 [M–H]$^-$.

**O-((6aS,8R,9R,9aS)-8-(2,4-Dioxo-3,4-dihydropyrimidin-1(2H)-yl)-2,2,4,4-tetraisopropyl tetrahydro-6H-furo[3,2-f][1,3,5,2,4]trioxadisilocin-9-yl) O-(p-tolyl) carbonothioate *(5)***

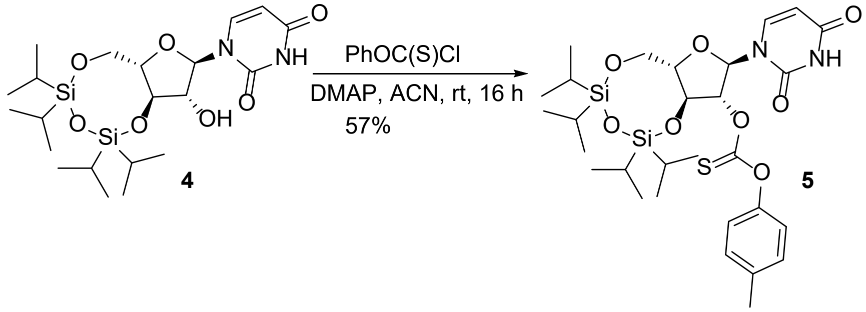


Compound **4** (99.80 g, 205.05 mmol) and 4-dimethylaminopyridine (75.15 g, 615.15 mmol) were dissolved in anhydrous acetonitrile (800 mL), followed by slow addition of O-4-methylphenyl chlorothioformate (63.26 mL, 410.10 mmol). The mixture was stirred at rt for 16 h. The solvent was removed under reduced pressure, and the residue was partitioned between ethyl acetate and water. The aqueous phase was extracted with ethyl acetate, and the combined organic extracts were washed sequentially with 10 % aqueous HCl, water, saturated sodium bicarbonate solution, water, and brine. The organic layer was dried over magnesium sulfate, filtered, and concentrated under reduced pressure. Purification by Biotage chromatography (Si, 350 g column, 0–40 % ethyl acetate/hexanes) afforded the desired product as a white solid.

**Yield:** 74 g (116.19 mmol, 57 %). **$^1$H NMR** (300 MHz, $CDCl_3$) δ 8.34 (br s, 1H), 7.13–7.37 (m, 4H), 6.95 (br d, J = 7.00 Hz, 2H), 6.28 (t, J = 5.33 Hz, 1H), 5.91 (d, J = 4.04 Hz, 1H), 5.76 (br d, J = 8.08 Hz, 1H), 4.75 (t, J = 7.14 Hz, 1H), 4.24–4.47 (m, 1H), 4.01 (br s, 2H), 2.36 (s, 3H), 0.94–1.18 (m, 29H).

**1-((6aS,8R,9aR)-2,2,4,4-tetraisopropyltetrahydro-6H-furo[3,2-f][1,3,5,2,4]trioxadisilocin-8-yl)pyrimidine-2,4(1H,3H)-dione *(6)***

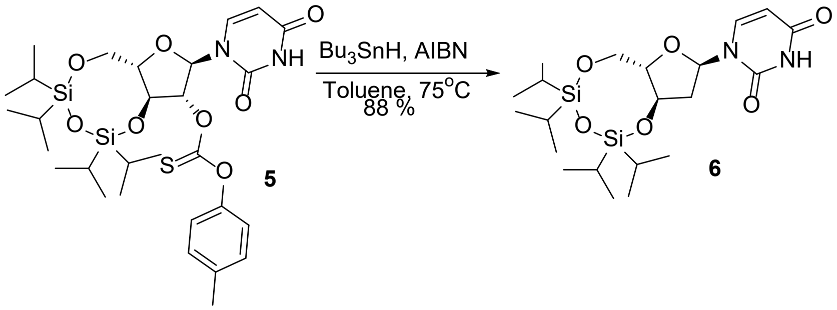


Azobisisobutyronitrile (3.79 g, 23.06 mmol) and tributyltin hydride (335.62 g, 115.0 mmol) in toluene (120 mL) were added dropwise to a degassed (nitrogen) solution of **5** (73.44 g, 115.31 mmol) in toluene (120 mL) at 80 °C. The reaction mixture was maintained at 80 °C for 1 h, then cooled to rt. The solvent was removed under reduced pressure, and the residue was purified by Biotage chromatography (Si, 350 g column, 0–30 % ethyl acetate/hexanes) to afford the desired product as a white solid. The product was used in subsequent steps without further characterization

**Yield:** 48 g (101.97 mmol, 88 %).; no spectra are available for this batch.

**1-((2R,4R,5S)-4-Hydroxy-5-(hydroxymethyl) tetrahydrofuran-2-yl)pyrimidine-2,4(1H,3H)-dione *(7)***

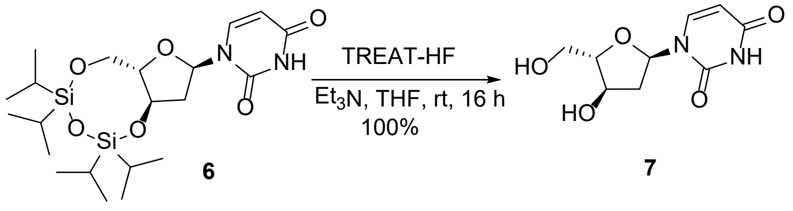


Triethylamine (34.60 mL, 249.62 mmol) was added to a solution of **6** (47 g, 99.85 mmol) in tetrahydrofuran (91 mL). The reaction was cooled to 0 °C under a nitrogen atmosphere using an ice bath. Triethylamine trihydrofluoride (81.88 mL, 499.24 mmol) was added slowly at 0 °C, and the reaction was allowed to warm to rt and stirred overnight. The solvent was removed under reduced pressure, and the residue was purified by Biotage chromatography (Si, 10 g column, 0–10 % methanol/dichloromethane) to afford the desired product as a white solid.

**Yield:** 22.79 g (99.87 mmol, 100 %). $^{1}$**H NMR** (300 MHz, DMSO-$d_6$) δ 11.20 (s, 1H), 7.87 (d, J = 7.91 Hz, 1H), 6.10 (dd, J = 2.56, 7.67 Hz, 1H), 5.62 (d, J = 8.08 Hz, 1H), 5.32 (d, J = 2.96 Hz, 1H), 4.83 (t, J = 5.65 Hz, 1H), 4.12–4.26 (m, 2H), 3.36–3.43 (m, 2H), 2.53–2.62 (m, 1H), 2.07 (s, 1H), 1.84–1.93 (m, 1H). $^{13}$**C NMR** (75 MHz, DMSO-$d_6$) δ 161.7, 148.9, 139.7, 99.4, 87.8, 84.1, 68.9, 60.1. **LC-MS:** m/z 229.20 $[M+H]^+$.

**1-((2R,4R,5S)-5-((bis(4-methoxyphenyl)(phenyl)methoxy)methyl)-4-hydroxytetrahydro furan-2-yl)pyrimidine-2,4(1H,3H)-dione *(8)***

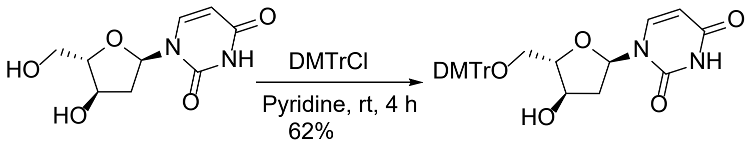


**7** **8**

4,4'-Dimethoxytrityl chloride (32.40 g, 96.40 mmol) was added to a solution of **7** (22 g, 96.40 mmol) in pyridine (192 mL) at rt. The mixture was stirred for 4 h and then quenched by addition of methanol (0.5 mL). The reaction was diluted with water and ethyl acetate, and the aqueous layer was extracted with ethyl acetate. The combined organic extracts were washed successively with water, saturated sodium bicarbonate solution, and brine, then concentrated under reduced pressure. Purification by Biotage chromatography (Si, 220 g column, 0–80 % ethyl acetate/hexanes) afforded the desired product as a white solid.

**Yield:** 31.60 g (59.56 mmol, 62 %). $^{1}$**H NMR** (300 MHz, $CDCl_3$) δ 8.96 (br s, 1H), 7.68 (d, J = 7.90 Hz, 1H), 7.35–7.45 (m, 2H), 7.19–7.33 (m, 9H), 6.80–6.87 (m, 4H), 6.20 (dd, J = 1.48, 7.32 Hz, 1H), 5.65 (dd, J = 1.80, 8.17 Hz, 1H), 4.43 (t, J = 3.81 Hz, 2H), 3.79 (s, 6H), 3.10–3.26 (m,

2H), 2.91 (br s, 1H), 2.68–2.82 (m, 1H), 2.24 (br d, J = 14.81 Hz, 1H). **$^{13}$C NMR** (75 MHz, $CDCl_3$) δ 163.8, 158.6, 150.3, 144.5, 141.4, 135.7, 135.6, 130.0, 128.1, 127.9, 127.0, 113.3, 101.2, 89.0, 88.4, 86.6, 72.7, 64.1, 55.2, 41.3. **LC-HRMS (ESI$^-$)** *m/z* 529.1979 [M−H]$^-$

**(2S,3R,5R)-2-((bis(4-methoxyphenyl)(phenyl)methoxy)methyl)-5-(2,4-dioxo-3,4-dihydropyrimidin-1(2H)-yl)tetrahydrofuran-3-yl (2-cyanoethyl) diisopropyl phosphoramidite *(9)***

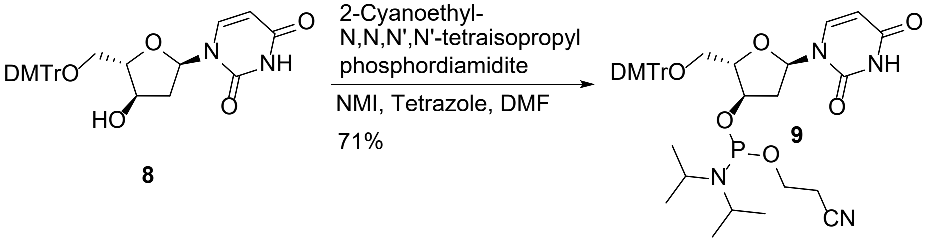


1*H*-tetrazole (478.90 mg, 6.94 mmol) and 1-methylimidazole (0.172 mL, 2.17 mmol) were added to a solution of **8** (4.60 g, 8.67 mmol) in DMF (42.0 mL) at rt under nitrogen. 2-Cyanoethyl-N,N,N',N'-tetraisopropylphosphorodiamidite (4.13 mL, 13.00 mmol) was added dropwise, and the reaction was stirred at rt for 4h. Water (1 mL) was then added to quench the reaction. A 3:1 mixture of toluene/hexanes (80 mL) was added, and the mixture was washed four times with 3:2 (v/v) DMF/water (50 mL). The upper organic phase was then washed with saturated sodium bicarbonate solution and brine, dried over sodium sulfate, and concentrated under reduced pressure to afford a white foam. Purification by Biotage chromatography (Si, 100 g column, 0–60 % ethyl acetate/hexanes) afforded the desired product as a white amorphous solid.

**Yield:** 4.50 g (6.16 mmol, 71 %). **$^1$H NMR** (300 MHz, $CDCl_3$) δ 8.46 (br s, 1H), 7.71 (d, J = 8.33 Hz, 1H), 7.37–7.44 (m, 2H), 7.19–7.35 (m, 9H), 6.80–6.88 (m, 4H), 6.28–6.36 (m, 1H), 5.69 (dd, J = 4.85, 8.17 Hz, 1H), 4.46–4.60 (m, 2H), 3.46–3.81 (m, 10H), 3.09–3.32 (m, 2H), 2.63–2.88 (m,

1H), 2.40–2.58 (m, 2H), 2.17–2.38 (m, 1H), 1.05–1.20 (m, 12H). **$^{31}$P NMR** (121 MHz, $CDCl_3$) δ 149.67 (s, 1P), 149.07 (s, 1P). **LC-HRMS (ESI$^-$)** *m/z* 729.3058 [M−H]$^-$

**Synthesis of aLd C phosphoramidite.** The synthesis of aLd C phosphoramidite has been reported previously in a patent, starting from aLd U via a 4-thiouridine intermediate.[3] This route requires harsh conditions, such as refluxing in dioxane with $P_2S_5$ and high-temperature ammonolysis. In this work, we used a more convenient approach employing a 4-triazolyl intermediate that reacts efficiently at room temperature and affords the target amidite under mild conditions.

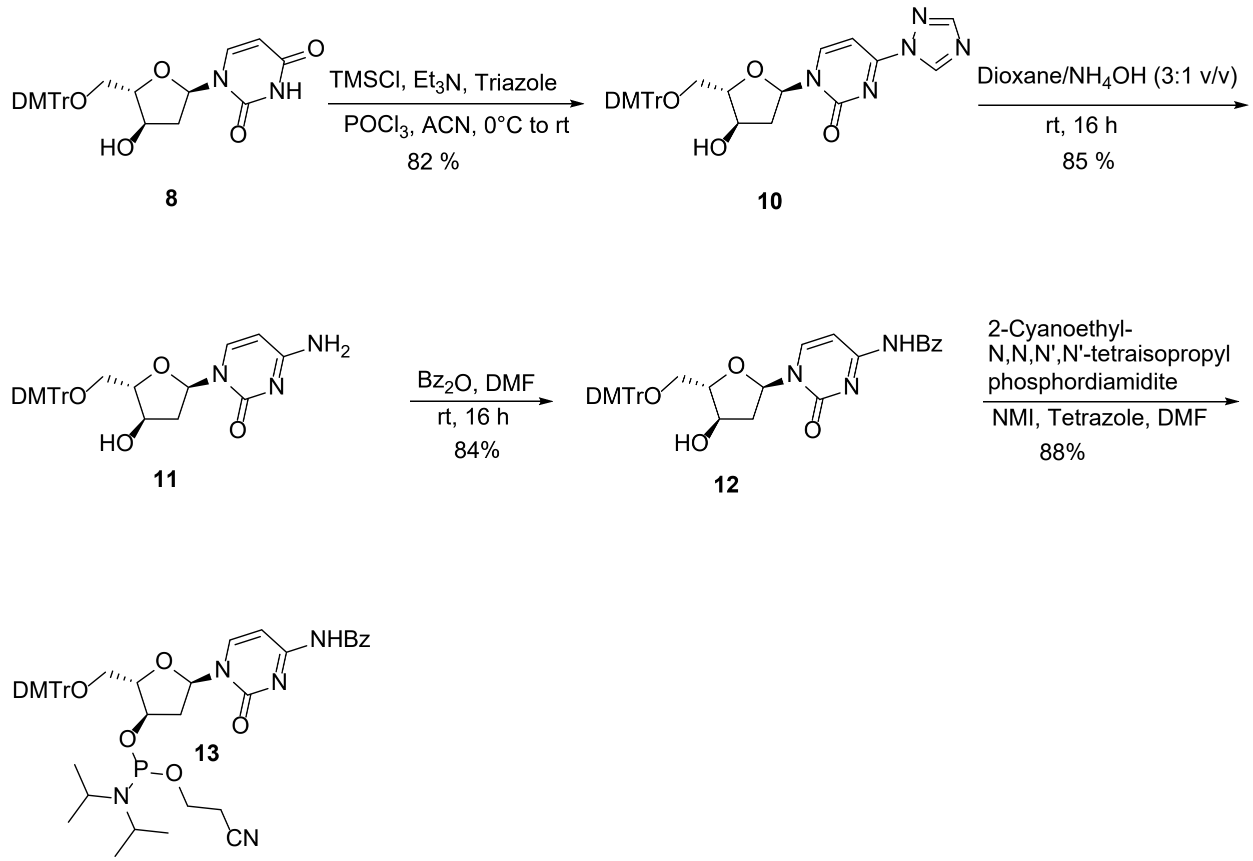

**1-((2R,4R,5S)-5-((Bis(4-methoxyphenyl)(phenyl)methoxy)methyl)-4-hydroxytetrahydro furan-2-yl)-4-(1H-1,2,4-triazol-1-yl)pyrimidin-2(1H)-one *(10)***

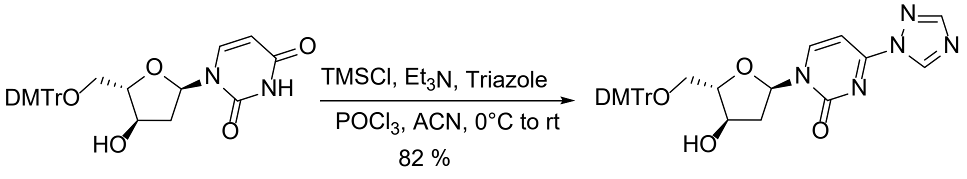


A suspension of **8** (16.72 g, 31.51 mmol) in anhydrous acetonitrile (400 mL) was treated with triethylamine (96.63 mL, 693.28 mmol) under nitrogen and stirred. The mixture was cooled in an ice bath, and chlorotrimethylsilane (20.0 mL, 157.56 mmol) was added dropwise over 15 min; stirring was continued for 20 min at rt. 1*H*-1,2,4-Triazole (32.65 g, 472.69 mmol) was added (note: temperature increased from 25 °C to 35 °C) and the mixture was stirred for 20 min. The reaction was cooled to 0 °C (ice bath), and phosphorus oxychloride (8.65 mL, 94.54 mmol) was added dropwise over 15 min. The ice bath was removed, and the reaction was stirred at rt for 2 h. The mixture was concentrated to a small volume under reduced pressure, diluted with ethyl acetate, and the organic layer was washed with saturated aqueous sodium bicarbonate (2×), water, and brine, then concentrated to a yellow oil. Purification by Biotage chromatography (Si, 100 g column, 50–100 % ethyl acetate/hexanes) afforded the desired product.

**Yield:** 15 g (25.79 mmol, 82 %). **$^1$H NMR** (300 MHz, $CDCl_3$) δ 9.27 (s, 1H), 8.27 (d, J = 7.15 Hz, 1H), 8.11 (s, 1H), 7.38–7.45 (m, 2H), 7.23–7.35 (m, 9H), 7.00 (d, J = 7.15 Hz, 1H), 6.80–6.90 (m, 4H), 6.28 (d, J = 6.28 Hz, 1H), 5.29 (s, 1H), 4.44–4.53 (m, 1H), 4.28 (d, J = 4.76 Hz, 1H), 3.80 (s, 6H), 3.20 (dd, J = 4.98, 7.94 Hz, 2H), 2.20 (d, J = 14.54 Hz, 1H), 2.04 (s, 1H). $^{13}$C NMR (75 MHz, $CDCl_3$) δ 159.4, 158.8, 154.8, 154.0, 147.8, 144.6, 143.4, 135.8, 135.7, 130.1, 128.2, 128.1, 127.2, 113.4, 93.3, 90.6, 90.1, 86.8, 72.8, 63.7, 55.4, 41.3.

**4-Amino-1-((2R,4R,5S)-5-((Bis(4-methoxyphenyl)(phenyl)methoxy)methyl)-4-hydroxytetrahydrofuran-2-yl)pyrimidin-2(1H)-one** ***(11)***

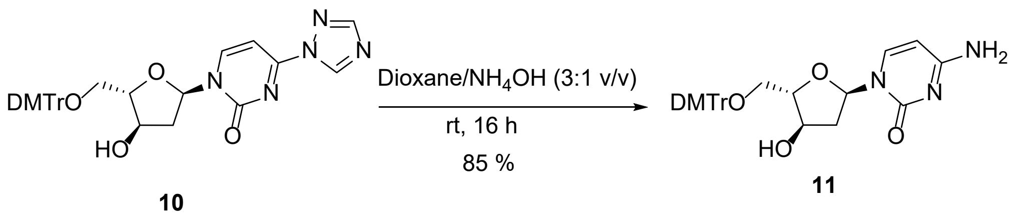


Compound **10** was dissolved in a 3:1 (v/v) mixture of dioxane/ammonium hydroxide (50 mL) and stirred at rt for 16 h. The solvent was removed under reduced pressure to give a white foam. Purification by Biotage chromatography (Si, 100 g column, 50–100 % ethyl acetate/hexanes) afforded the desired product as a white solid.

**Yield:** 11.60 g (21.9 mmol, 85 %). **$^1$H NMR** (300 MHz, DMSO-$d_6$) δ 13.84–14.22 (m, 1H), 8.17–8.74 (m, 1H), 8.08 (br d, J = 7.45 Hz, 1H), 7.79 (d, J = 7.45 Hz, 1H), 7.18–7.43 (m, 10H), 6.97–7.16 (m, 2H), 6.91 (d, J = 8.80 Hz, 4H), 6.18 (dd, J = 2.92, 7.32 Hz, 1H), 5.75 (d, J = 7.36 Hz, 1H), 5.28 (d, J = 3.14 Hz, 1H), 4.24–4.37 (m, 1H), 4.04–4.22 (m, 1H), 3.74 (s, 6H), 2.94–3.21 (m, 3H), 2.49–2.62 (m, 2H), 1.89 (br d, J = 14.09 Hz, 1H). **$^{13}$C NMR** (75 MHz, DMSO-$d_6$) δ 166.2, 158.6, 155.8, 145.3, 142.1, 136.0, 136.0, 130.2, 128.3, 128.2, 127.2, 113.7, 93.9, 87.7, 87.0, 86.1, 71.5, 64.6, 55.5, 49.1, 41.2. **LC-MS:** m/z 528.20 $[M–H]^-$.

**N-(1-((2R,4R,5R)-5-((Bis(4-methoxyphenyl)(phenyl)methoxy)methyl)-4-hydroxytetrahydrofuran-2-yl)-2-oxo-1,2-dihydropyrimidin-4-yl)benzamide** ***(12)***

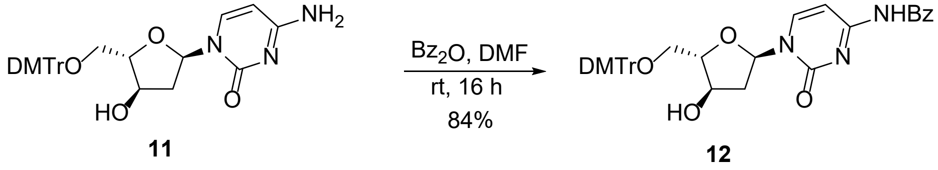

Benzoic anhydride (3.20 g, 14.12 mmol) in DMF (10 mL) was added dropwise to a solution of **11** (6.8 g, 12.84 mmol) in anhydrous DMF (40 mL) at rt. The reaction was stirred overnight. The mixture was quenched with methanol (0.5 mL), diluted with a 1:1 mixture of water/ethyl acetate (100 mL), and the layers were separated. The organic layer was washed with saturated sodium bicarbonate (1 × 50.0 mL) and brine (1 × 50.0 mL), dried over sodium sulfate, filtered, and concentrated under reduced pressure to a foam. Purification by Biotage chromatography (Si, 100 g column, 0–80 % ethyl acetate/hexanes) afforded the desired product as a white solid.

**Yield:** 6.80 g (10.73 mmol, 84 %). **$^{1}$H NMR** (300 MHz, $CDCl_3$) δ 8.59 (br s, 1H), 8.08 (dd, J = 7.45, 13.82 Hz, 1H), 7.89 (br d, J = 7.36 Hz, 2H), 7.39–7.64 (m, 6H), 7.20–7.36 (m, 10H), 6.81–6.89 (m, 4H), 6.25–6.37 (m, 1H), 4.70 (s, 1H), 4.46–4.61 (m, 1H), 3.80 (s, 6H), 3.53–3.74 (m, 2H), 3.15–3.51 (m, 4H), 2.35–2.63 (m, 3H). **$^{13}$C NMR** (75 MHz, $CDCl_3$) δ 162.2, 158.6, 145.6, 144.6, 135.8, 135.7, 133.1, 130.0, 129.0, 128.1, 127.9, 127.6, 126.9, 113.2, 90.4, 89.7, 86.5, 72.8, 64.2, 55.2, 41.5, 36.5. **LC-HRMS (ESI$^-$)** *m/z* 632.2399 $[M-H]^-$

**(2S,3R,5R)-5-(4-Benzamido-2-oxopyrimidin-1(2H)-yl)-2-((Bis(4-methoxyphenyl)(phenyl) methoxy)methyl)tetrahydrofuran-3-yl (2-cyanoethyl) diisopropylphosphoramidite** ***(13)***

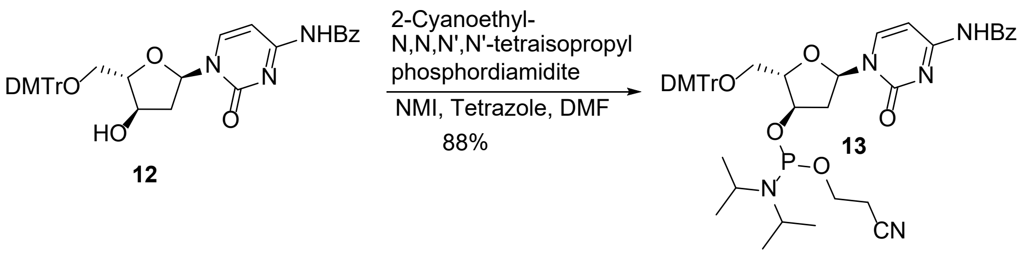


1*H*-Tetrazole (0.592 g, 8.58 mmol) and 1-methylimidazole (0.213 mL, 2.68 mmol) were added to a solution of **12** (6.80 g, 10.73 mmol) in DMF (42 mL), followed by dropwise addition of 2-cyanoethyl-N,N,N',N'-tetraisopropylphosphorodiamidite (5.11 mL, 16.10 mmol). The reaction was stirred at rt for 90 min. Water (1 mL) was added to quench the reaction. A 3:1 mixture of toluene/hexanes (80 mL) was added, and the mixture was washed four times with 3:2 (v/v) DMF/water (50 mL). The upper organic phase was then washed with saturated sodium bicarbonate solution and brine, dried over sodium sulfate, and concentrated under reduced pressure. Purification by Biotage chromatography (Si, 220 g column, 80–100 % ethyl acetate/hexanes) with the sample preloaded in a small amount of ethyl acetate, afforded the desired product as a white amorphous solid.

**Yield:** 7.83 g (9.39 mmol, 88 %). **$^{1}$H NMR** (300 MHz, $CDCl_3$) δ 8.59 (br s, 1H), 8.08 (dd, J = 7.45, 13.82 Hz, 1H), 7.89 (br d, J = 7.36 Hz, 2H), 7.39–7.64 (m, 6H), 7.18–7.36 (m, 10H), 6.81–6.89 (m, 4H), 6.25–6.37 (m, 1H), 4.70 (t, J = 3.77 Hz, 1H), 4.46–4.61 (m, 1H), 3.80 (s, 6H), 3.54–3.74 (m, 2H), 3.15–3.51 (m, 4H), 2.78–2.94 (m, 1H), 2.67–2.78 (m, 1H), 2.35–2.63 (m, 3H), 0.98–1.20 (m, 12H). **$^{31}$P NMR** (121 MHz, $CDCl_3$) δ 149.52 (s, 1P), 149.11 (s, 1P). **LC-HRMS ($ESI^-$)** *m/z* 832.3480 $[M{-}H]^-$

**Synthesis of aLd A phosphoramidite.** The synthesis of aLd A has been reported previously starting from aLd U via a transglycosylation reaction, as direct construction of the purine ring on the L-sugar was considered difficult.[2] The corresponding phosphoramidite has also appeared in a patent without synthetic or characterization details.[3] In this work, we start from 1-O-acetyl-2,3,5-tri-O-benzoyl-L-arabinofuranose (compound **1**) and construct the purine ring directly on the L-sugar.

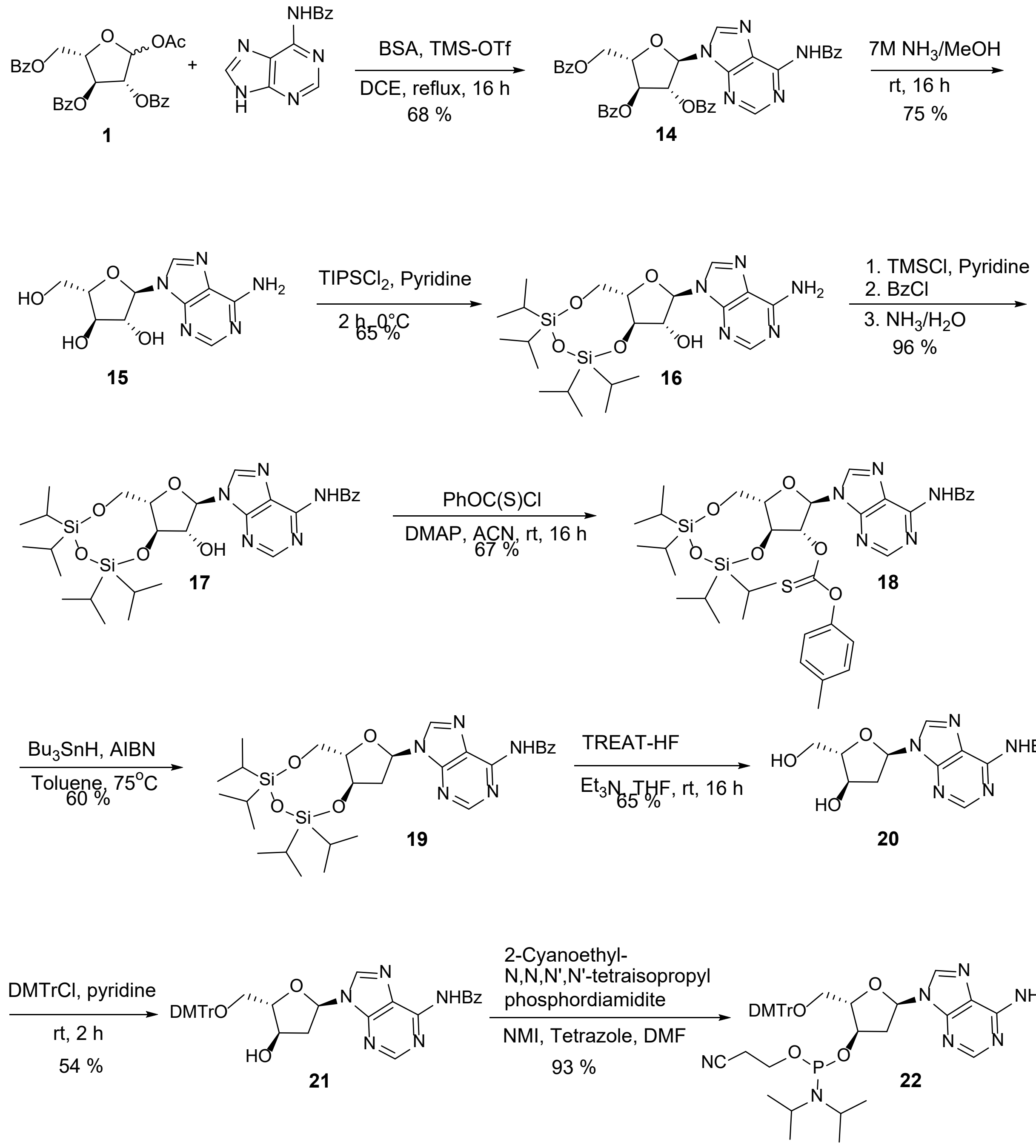


**(2R,3R,4S,5S)-2-(6-Benzamido-9H-purin-9-yl)-5-((benzoyloxy)methyl)tetrahydrofuran-3,4-diyl dibenzoate *(14)***

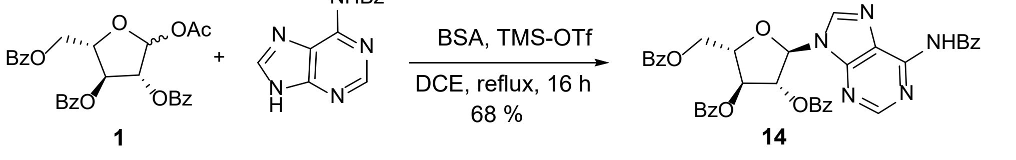

$N^6$-Benzoyladenine (23.40 g, 97.30 mmol, 1.30 equiv) and **1** (38.0 g, 75.3 mmol) were co-evaporated with toluene (4 × 50 mL) at 60 °C. The residue was suspended in anhydrous 1,2-dichloroethane (800 mL) and N,O-bis(trimethylsilyl)acetamide (73.7 mL, 301 mmol, 4.0 equiv) was added. After refluxing at 80 °C for 1 h to obtain a clear solution, the mixture was cooled in an ice bath to 5 °C, trimethylsilyl trifluoromethanesulfonate (21.80 mL, 121 mmol, 1.6 equiv) was added, and the reaction was heated at reflux overnight. The mixture was concentrated under reduced pressure, diluted with ethyl acetate (200 mL), and the organic phase was washed with deionized water (200 mL), then saturated aqueous sodium bicarbonate to pH 7, and brine. The organic layer was dried over sodium sulfate, filtered, and concentrated under reduced pressure. Purification by Biotage chromatography (Si, 320 g column, 0–5 % methanol/dichloromethane) afforded the product as a white solid.

**Yield:** 35.2 g (51.5 mmol, 68 %). **$^1$H NMR** (300 MHz, $CDCl_3$) δ 9.04 (s, 1H), 8.83 (s, 1H), 8.30 (s, 1H), 8.00–8.12 (m, 6H), 7.88–7.98 (m, 2H), 7.35–7.65 (m, 12H), 6.52–6.59 (m, 2H), 5.93 (dd, J = 2.78, 4.49 Hz, 1H), 5.12 (q, J = 4.67 Hz, 1H), 4.80 (d, J = 4.94 Hz, 2H).**$^{13}$C NMR** (75 MHz, $CDCl_3$) δ 166.1, 165.5, 165.4, 164.5, 153.1, 151.7, 149.7, 141.5, 134.0, 133.6, 133.3, 132.9, 130.1, 129.8, 129.4, 128.9, 128.7, 128.5, 128.3, 127.9, 123.5, 89.3, 83.2, 80.6, 63.6. **LC-MS:** m/z 684.2 $[M+H]^+$.

**(2R,3R,4R,5S)-2-(6-Amino-9H-purin-9-yl)-5-(hydroxymethyl)tetrahydrofuran-3,4-diol** ***(15)***

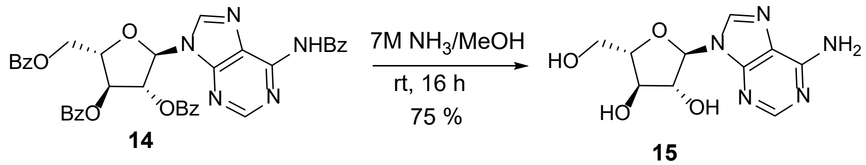

Compound **14** (43.0 g, 58.50 mmol) was suspended in methanol (50 mL) and cooled to −20 °C. A solution of ammonia in methanol (7 N, 150 mL) was added, and the mixture was heated at 45 °C overnight. The mixture was concentrated to an oil, suspended in ethyl acetate (100 mL) to precipitate a white solid, which was collected by filtration, rinsed with fresh ethyl acetate, and dried under high vacuum.

**Yield:** 11.7 g (43.8 mmol, 75 %). **$^{1}$H NMR** (300 MHz, DMSO-$d_6$) δ 8.32 (s, 1H), 8.17 (s, 1H), 7.26 (s, 2H), 5.86 (dd, J = 3.19, 5.07 Hz, 2H), 5.73 (d, J = 5.03 Hz, 1H), 5.00 (t, J = 5.52 Hz, 1H), 4.66 (q, J = 5.30 Hz, 1H), 4.13–4.21 (m, 1H), 3.80–4.05 (m, 2H), 3.36–3.55 (m, 2H). **$^{13}$C NMR** (75 MHz, DMSO-$d_6$) δ 156.4, 153.0, 149.6, 140.5, 119.5, 88.9, 85.7, 79.7, 75.7, 61.6. **LC-MS:** m/z 268.1 $[M+H]^+$.

**(6aS,8R,9R,9aR)-8-(6-Amino-9H-purin-9-yl)-2,2,4,4-tetraisopropyltetrahydro-6H-furo[3,2-f][1,3,5,2,4]trioxadisilocin-9-ol** ***(16)***

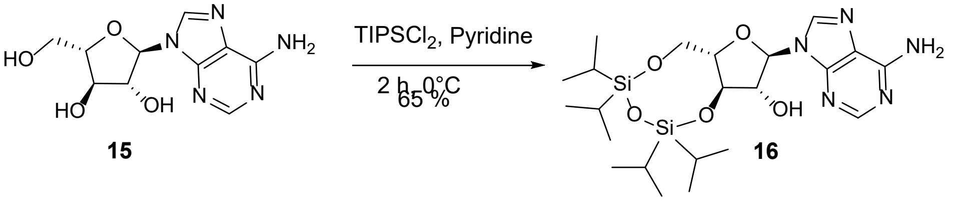


Compound **15** (11.76 g, 43.78 mmol) was dissolved in pyridine (400 mL) under nitrogen, cooled in an ice bath to 0 °C, and 1,3-dichloro-1,1,3,3-tetraisopropyldisiloxane (12.66 mL, 39.60 mmol, 0.90 equiv) was added dropwise. The reaction was allowed to warm to ~10 °C and stirred for 2 h (TLC in ethyl acetate/hexanes 8:2 indicated completion). The mixture was cooled to 0 °C and quenched by slow addition of water (20 mL), then concentrated to an oil under reduced pressure. The residue was dissolved in ethyl acetate and the organic layer was washed with 10 % aqueous

HCl, water, saturated sodium bicarbonate solution, water, and brine, then concentrated to give a colorless oil. The crude oil was suspended in hexanes to induce precipitation.

**Yield:** 14 g (27.5 mmol, 65 %). **$^1$H NMR** (300 MHz, DMSO-$d_6$) δ 8.13 (s, 1H), 7.93 (s, 1H), 7.00 (s, 2H), 5.73 (br d, J = 5.92 Hz, 1H), 5.56 (d, J = 6.10 Hz, 1H), 4.81 (br d, J = 6.19 Hz, 1H), 3.92–4.16 (m, 2H), 3.58–3.79 (m, 2H), 2.26 (br s, 1H), 0.67–0.93 (m, 25H). **$^{13}$C NMR** (75 MHz, DMSO-$d_6$) δ 156.5, 153.1, 149.8, 141.1, 119.8, 87.5, 81.7, 77.2, 75.5, 61.3, 17.7, 17.6, 17.5, 17.4, 17.3, 13.4, 13.0, 12.7, 12.5. **LC-MS:** m/z 510.3 $[M+H]^+$.

**N-(9-((6aS,8R,9R,9aR)-9-Hydroxy-2,2,4,4-tetraisopropyltetrahydro-6H-furo[3,2-f][1,3,5,2,4]trioxadisilocin-8-yl)-9H-purin-6-yl)benzamide** ***(17)***

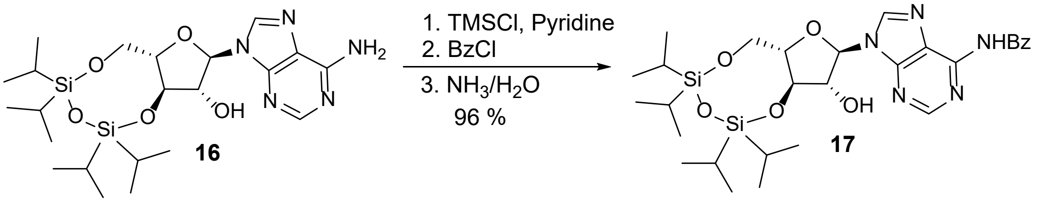


Compound **16** (7.90 g, 15.50 mmol) was dissolved in pyridine (100 mL) under nitrogen, cooled in an ice bath, and trimethylsilyl chloride (13.80 mL, 108 mmol, 5.0 equiv) was added dropwise. The ice bath was removed, and the mixture was stirred for 1 h at rt. The reaction was re-cooled and benzoyl chloride (9.00 mL, 77.50 mmol, 5.0 equiv) was added dropwise; the mixture was warmed to rt and stirred overnight. The reaction was cooled, and water (150 mL) was added dropwise, keeping the temperature below 7 °C, then stirred at rt for 1 h. The mixture was cooled again, and ammonium hydroxide (100 mL) was added dropwise and stirred for 30 min. Volatiles were partially evaporated at rt, the mixture was diluted with ethyl acetate and washed with water, saturated sodium bicarbonate, and brine; the organic layer was dried over sodium sulfate, filtered, and concentrated. Purification by Biotage chromatography (Si, 100 g column, 0–5 % methanol/dichloromethane) afforded a white solid.

**Yield:** 9.20 g (15.0 mmol, 96 %). **$^1$H NMR** (300 MHz, $CDCl_3$) δ 9.31, 8.65, 8.12, 8.00, 7.78, 7.38–7.62, 7.25–7.31, 5.91, 4.99, 4.55, 4.26, 3.99–4.15, 2.04, 1.25, 1.00–1.18 (29H). **$^{13}$C NMR** (75 MHz, DMSO-$d_6$) δ 168.3, 166.1, 152.7, 152.2, 151.0, 144.6, 134.7, 133.8, 132.9, 131.7, 128.9, 128.6, 127.9, 126.6, 87.5, 81.7, 77.1, 75.2, 61.3, 17.8, 17.7, 17.6, 17.5, 17.4, 17.3, 13.4, 13.0, 12.7, 12.5. **LC-MS:** m/z 614.3 $[M+H]^+$.

**O-((6aS,8R,9R,9aS)-8-(6-Benzamido-9H-purin-9-yl)-2,2,4,4-tetraisopropyltetrahydro-6H-furo[3,2-f][1,3,5,2,4]trioxadisilocin-9-yl) O-(p-tolyl) carbonothioate *(18)***

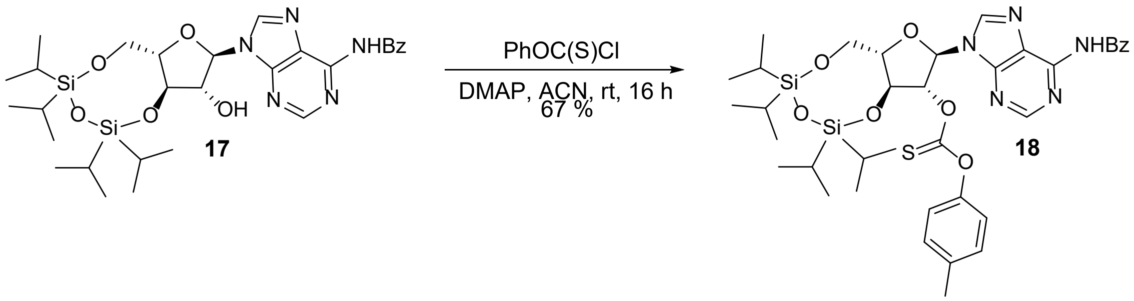


Compound **17** (8.00 g, 13.0 mmol) and 4-dimethylaminopyridine (3.18 g, 26.1 mmol, 2.0 equiv) were dissolved in anhydrous acetonitrile (131 mL), and anhydrous tetrahydrofuran (50 mL) was added to assist dissolution. O-(4-Methylphenyl) chlorothioformate (2.18 mL, 14.3 mmol, 1.2 equiv) was added dropwise, and the reaction mixture was stirred at rt for 16 h (monitored by TLC, dichloromethane/methanol 95:5). The solvent was evaporated under reduced pressure, and the residue was partitioned between ethyl acetate and water. The aqueous layer was extracted with ethyl acetate (2×). The combined organic extracts were washed sequentially with 10% aqueous HCl, water, saturated sodium bicarbonate, water, and brine, then dried over magnesium sulfate, filtered, and concentrated. Purification by Biotage silica chromatography (100 g column, 0–3% methanol/dichloromethane) gave the desired product as a white solid.

**Yield:** 6.67 g (8.73 mmol, 67 %). $^1$**H NMR** (300 MHz, $CDCl_3$) δ 9.05 (s, 1H), 8.86 (s, 1H), 8.22 (s, 1H), 7.96–8.07 (m, 2H), 7.45–7.64 (m, 3H), 7.11–7.28 (m, 3H), 6.88–6.96 (m, 2H), 6.72 (dd, J = 4.16, 6.08 Hz, 1H), 6.29 (d, J = 4.03 Hz, 1H), 4.85 (dd, J = 6.14, 8.06 Hz, 1H), 4.55–4.66 (m, 1H), 4.06 (dd, J = 3.71, 8.83 Hz, 2H), 2.32 (s, 3H), 1.88 (br s, 1H), 1.26 (s, 1H), 1.03–1.19 (m, 31H). **LC-MS:** m/z 763.3 $[M–H]^-$.

**N-(9-((6aS,8R,9aR)-2,2,4,4-Tetraisopropyltetrahydro-6H-furo[3,2-f][1,3,5,2,4]trioxadisilocin-8-yl)-9H-purin-6-yl)benzamide** ***(19)***

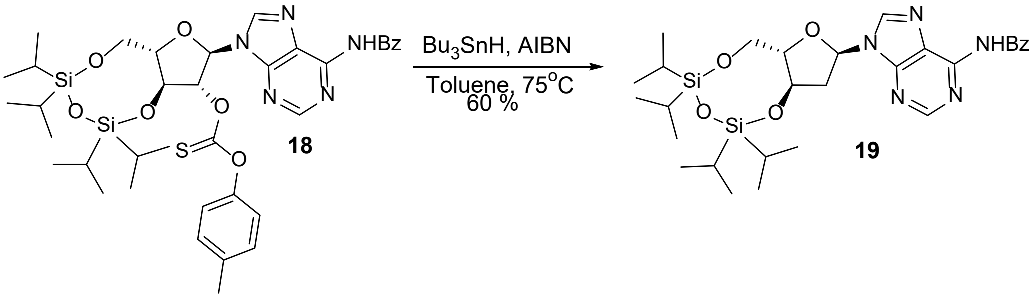


A degassed solution (nitrogen, 30 min) of **18** (6.67 g, 8.73 mmol) in toluene (140 mL) was heated to 80 °C. A solution of azobisisobutyronitrile (0.287 g, 1.75 mmol, 0.20 equiv) and tributyltin hydride (23.50 mL, 87.30 mmol, 10 equiv) in toluene (40 mL) was added dropwise at 80 °C. The reaction was maintained at 80 °C for 1 h, cooled to rt, and concentrated. TLC in ethyl acetate/hexanes (7:3) indicated conversion. Purification by Biotage chromatography (Si, 100 g column, 70 % ethyl acetate/hexanes) afforded a white solid.

**Yield:** 3.0 g (5.15 mmol, 60 %). $^1$**H NMR** (300 MHz, $CDCl_3$) δ 9.02 (s, 1H), 8.84 (s, 1H), 8.54 (s, 1H), 7.97–8.06 (m, 3H), 7.43–7.64 (m, 4H), 7.27 (s, 1H), 6.51 (dd, J = 3.65, 7.23 Hz, 1H), 4.72

(td, J = 4.94, 8.29 Hz, 2H), 3.98–4.21 (m, 3H), 3.82 (dd, J = 7.94, 11.78 Hz, 1H), 2.91–3.03 (m, 1H), 2.66–2.77 (m, 1H), 1.83 (br s, 1H), 1.26 (s, 1H), 0.92–1.18 (m, 39H). **$^{13}$C NMR** (75 MHz, $CDCl_3$) δ 164.6, 152.9, 149.5, 141.8, 133.7, 132.8, 129.8, 128.9, 128.5, 127.9, 86.2, 83.1, 72.9, 63.2, 39.7, 17.4, 17.1, 17.0, 13.4, 13.3, 13.0, 12.5. **LC-MS:** m/z 598.3 $[M+H]^+$.

**N-(9-((2R,4R,5S)-4-Hydroxy-5-(hydroxymethyl)tetrahydrofuran-2-yl)-9H-purin-6-yl)benzamide *(20)***

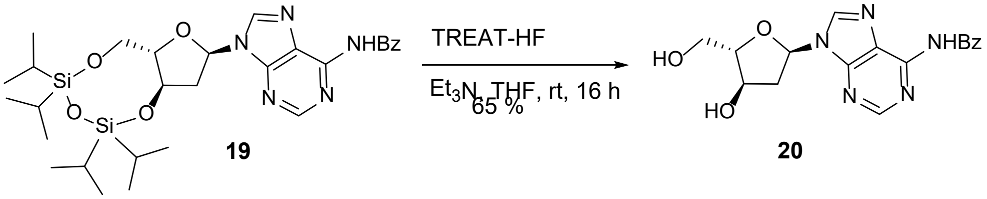


Triethylamine (1.36 mL, 9.80 mmol, 2.5 equiv) was added to a solution of **19** (2.34 g, 3.91 mmol) in tetrahydrofuran (30 mL) at 0 °C under nitrogen. Triethylamine trihydrofluoride (3.19 mL, 20.0 mmol, 5.0 equiv) was added slowly at 0 °C, then the reaction was allowed to warm to rt and stirred for 16 h. The solvent was removed, and the residue was purified by a silica gel plug (50 g, 5–10 % methanol/dichloromethane) to afford a white solid.

**Yield:** 900 mg (2.53 mmol, 65 %). **$^{1}$H NMR** (300 MHz, $CDCl_3$) δ 9.10 (br s, 1H), 8.81 (s, 1H), 8.22 (s, 1H), 7.93–8.06 (m, 2H), 7.48–7.66 (m, 3H), 6.37 (dd, J = 2.11, 8.77 Hz, 1H), 6.21 (br d, J = 9.09 Hz, 1H), 4.56 (br s, 1H), 4.40–4.47 (m, 1H), 4.12 (q, J = 7.08 Hz, 1H), 3.71–3.89 (m, 2H), 3.08 (td, J = 8.42, 15.30 Hz, 1H), 2.63 (br d, J = 15.23 Hz, 1H), 1.68 (br s, 2H). **$^{13}$C NMR** (75 MHz, DMSO-$d_6$) δ 166.0, 151.9, 150.6, 143.7, 133.9, 132.9, 128.9, 126.1, 91.8, 89.4, 84.4, 71.1, 71.0, 65.6, 62.1, 45.8, 8.9. **LC-MS:** m/z 356.2 $[M+H]^+$.

**N-(9-((2R,4R,5S)-5-((Bis(4-methoxyphenyl)(phenyl)methoxy)methyl)-4-hydroxytetrahydro furan-2-yl)-9H-purin-6-yl)benzamide *(21)***

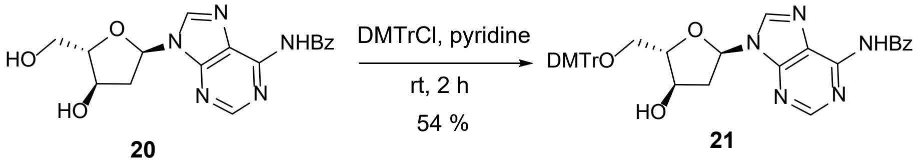


4,4'-Dimethoxytrityl chloride (1.10 g, 3.04 mmol, 1.2 equiv) was added to a solution of **20** (900 mg, 2.53 mmol) in pyridine (20 mL) at rt and stirred for 2 h. The reaction was quenched with methanol (2 mL), diluted with water and ethyl acetate, and the aqueous layer was extracted with ethyl acetate. The combined organics were washed with water, saturated sodium bicarbonate, and brine, dried over sodium sulfate, filtered, and concentrated. The crude material was dissolved in dichloromethane and purified by Biotage chromatography (Si, 50 g column, 0–5 % methanol/dichloromethane) to give a white solid.

**Yield:** 900 mg (1.37 mmol, 54 %). **$^{1}$H NMR** (300 MHz, $CDCl_3$) δ 9.19 (s, 1H), 8.77 (s, 1H), 8.23 (s, 1H), 7.96–8.06 (m, 2H), 7.40–7.62 (m, 5H), 7.20–7.38 (m, 8H), 6.77–6.93 (m, 4H), 6.31–6.45 (m, 2H), 4.44–4.54 (m, 2H), 3.79 (s, 6H), 3.29–3.43 (m, 1H), 3.08–3.27 (m, 2H), 2.56 (br d, J = 14.90 Hz, 1H), 2.02–2.10 (m, 1H). **$^{13}$C NMR** (75 MHz, $CDCl_3$) δ 164.6, 158.6, 151.9, 150.2, 150.0, 144.6, 143.4, 135.8, 135.6, 133.5, 132.9, 130.0, 128.9, 128.1, 128.0, 127.9, 127.0, 124.1, 113.3, 89.4, 86.9, 86.7, 73.3, 64.6, 55.3, 41.2. **LC-HRMS (ESI$^{-}$)** *m/z* 656.2511 $[M{-}H]^{-}$

**(2S,3R,5R)-5-(6-Benzamido-9H-purin-9-yl)-2-((Bis(4-methoxyphenyl)(phenyl)methoxy) methyl)tetrahydrofuran-3-yl (2-cyanoethyl) diisopropylphosphoramidite *(22)***

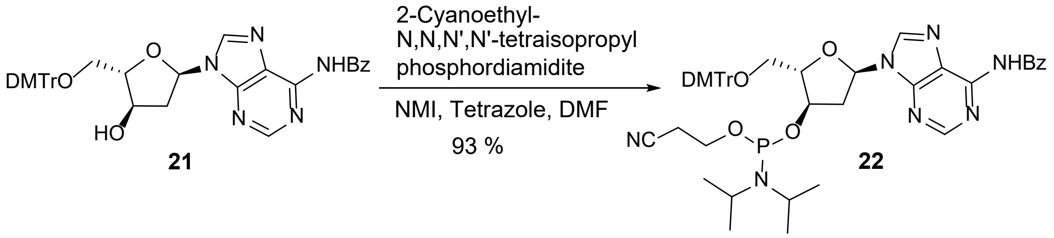


1*H*-Tetrazole (0.075 g, 1.09 mmol, 0.80 equiv) and 1-methylimidazole (0.0271 mL, 0.342 mmol, 0.25 equiv) were added to a solution of **21** (900 mg, 1.37 mmol) in DMF (10 mL) at rt under nitrogen. 2-Cyanoethyl-N,N,N',N'-tetraisopropylphosphorodiamidite (0.652 mL, 2.05 mmol, 1.5 equiv) was added dropwise, and the reaction was stirred at rt for 16 h. Water (1 mL) was added to quench the reaction. A 3:1 mixture of toluene/hexanes (80 mL) was added, and the mixture was washed four times with 3:2 (v/v) DMF/water (50 mL). The upper organic layer was then washed with saturated sodium bicarbonate solution and brine, dried over sodium sulfate, and concentrated under reduced pressure to a white foam. Purification by a silica gel plug (30 g, ethyl acetate/hexanes 9:1) afforded a white amorphous solid.

**Yield:** 1.10 g (1.28 mmol, 93 %). **$^{1}$H NMR** (300 MHz, $CDCl_3$) δ 9.13 (br s, 1H), 8.83 (s, 1H), 8.51 (d, J = 6.26 Hz, 1H), 8.04 (d, J = 7.17 Hz, 2H), 7.42–7.63 (m, 5H), 7.21–7.37 (m, 8H), 6.82–6.90 (m, 4H), 6.70 (t, J = 7.23 Hz, 1H), 4.56–4.70 (m, 2H), 3.80 (d, J = 1.02 Hz, 6H), 3.37–3.69 (m, 5H), 3.32 (dd, J = 4.74, 10.24 Hz, 1H), 3.17–3.25 (m, 1H), 2.88–3.06 (m, 1H), 2.56–2.81 (m, 1H), 2.41–2.53 (m, 2H), 0.98–1.20 (m, 12H). **$^{31}$P NMR** (121 MHz, $CDCl_3$) δ 149.46 (s, 1P), 149.03 (s, 1P). **LC-HRMS (ESI$^-$)** *m/z* 856.3589 $[M{-}H]^-$

**Synthesis of aLd G phosphoramidite.** The synthesis of aLd G has not been reported previously. Here, we describe its preparation starting from 1-O-acetyl-2,3,5-tri-O-benzoyl-L-arabinofuranose (compound 1), constructing the purine ring directly on the L-sugar.

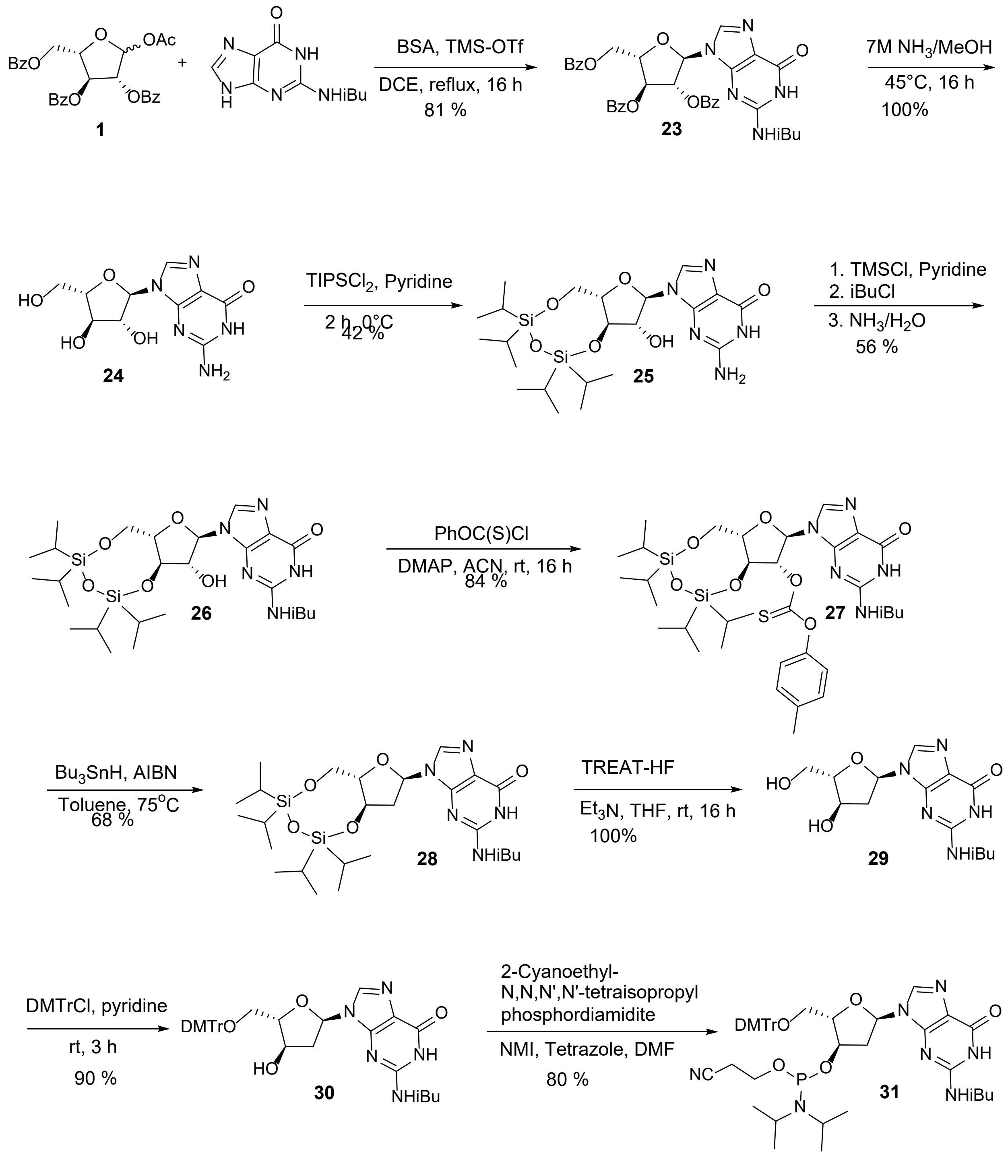

**(2S,3S,4R,5R)-2-((benzoyloxy)methyl)-5-(2-isobutyramido-6-oxo-1,6-dihydro-9H-purin-9-yl)tetrahydrofuran-3,4-diyl dibenzoate** ***(23)***

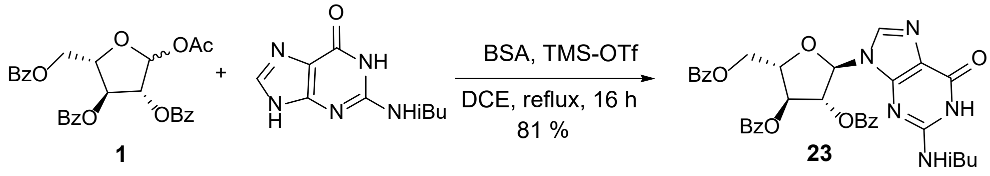


$N^2$-Isobutyrylguanine (23.0 g, 119 mmol) and **1** (40.0 g, 79.3 mmol) were co-evaporated with toluene (4 × 50 mL) at 60 °C. The residue was suspended in anhydrous 1,2-dichloroethane (800 mL), then N,O-bis(trimethylsilyl)acetamide (75.5 mL, 317 mmol) was added. After refluxing at 80 °C for 1 h to obtain a clear solution, the mixture was cooled in an ice bath to 5 °C, trimethylsilyl trifluoromethanesulfonate (23.0 mL, 127 mmol) was added, and the reaction was stirred overnight at 80 °C. The mixture was concentrated, diluted with ethyl acetate, and the organic phase was washed with deionized water, saturated aqueous sodium bicarbonate (to pH 7), and brine. The organics were concentrated to an oil. Purification by silica gel chromatography (packed glass column, ~1000 mL silica; diethyl ether/hexanes 6:4) afforded a white solid.

**Yield:** 43.0 g (64.60 mmol, 81 %). $^{1}$**H NMR** (300 MHz, $CDCl_3$) δ 12.09 (s, 1H), 8.91 (s, 1H), 7.94–8.11 (m, 6H), 7.83 (d, J = 7.72 Hz, 2H), 7.27–7.62 (m, 11H), 6.34–6.37 (m, 1H), 6.30 (s, 1H), 5.75–5.87 (m, 1H), 4.95–5.14 (m, 1H), 4.75 (dd, J = 2.78, 4.94 Hz, 2H), 2.52–2.67 (m, 1H), 1.14–1.30 (m, 6H). $^{13}$**C NMR** (75 MHz, $CDCl_3$) δ 178.3, 166.1, 165.3, 165.1, 155.6, 147.9, 147.6, 136.9, 134.1, 134.0, 133.4, 130.0, 129.9, 129.8, 129.6, 129.4, 128.7, 128.5, 128.3, 121.8, 88.9, 83.4, 80.1, 63.6, 36.3, 19.0, 18.9. **LC-MS:** m/z 666.2 $[M+H]^+$.

**2-Amino-9-((2R,3R,4R,5S)-3,4-dihydroxy-5-(hydroxymethyl)tetrahydrofuran-2-yl)-1,9-dihydro-6H-purin-6-one** ***(24)***

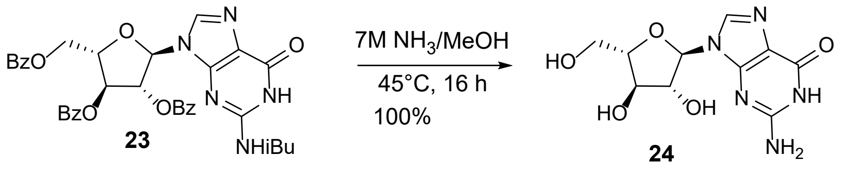


Compound **23** (43.0 g, 64.64 mmol) was suspended in methanol (50.0 mL) and cooled to –20 °C. Ammonia in methanol (7.00 M, 150 mL) was added and the mixture was heated at 45 °C for 16 h. The solution was concentrated to an oil; the residue was suspended in ethyl acetate (100 mL) to precipitate a white solid, which was filtered, rinsed with ethyl acetate, and dried under high vacuum.

**Yield:** 18.3 g (64.64 mmol, 100%). **$^1$H NMR** (300 MHz, DMSO-d$_6$) δ 7.80–8.01 (m, 1H), 7.53 (s, 1H), 6.49 (br s, 2H), 5.67 (d, J = 4.94 Hz, 2H), 4.48 (t, J = 5.16 Hz, 1H), 4.01–4.20 (m, 1H), 3.91–4.00 (m, 1H), 3.86 (s, 1H). **$^{13}$C NMR** (75 MHz, DMSO-d$_6$) δ 157.5, 154.2, 151.7, 136.4, 117.1, 87.9, 85.5, 80.1, 75.7, 61.6. **LC-MS:** m/z 282.2 [M–H]$^-$.

**2-Amino-9-((6aS,8R,9R,9aR)-9-hydroxy-2,2,4,4-tetraisopropyltetrahydro-6H-furo[3,2-f][1,3,5,2,4]trioxadisilocin-8-yl)-1,9-dihydro-6H-purin-6-one** ***(25)***

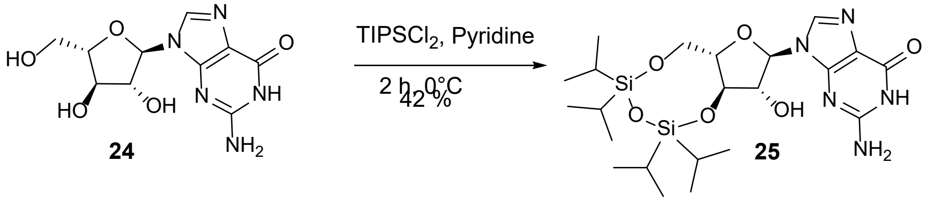


Compound **24** (18.3 g, 64.64 mmol) was dissolved in pyridine (400 mL) under nitrogen and cooled in an ice bath. 1,3-Dichloro-1,1,3,3-tetraisopropyldisiloxane (23.30 mL, 63.60 mmol, 0.98 equiv) was added dropwise. The mixture was allowed to warm to ~10 °C and stirred for 2 h (monitored by TLC, ethyl acetate/hexanes 8:2). The reaction was cooled to 0 °C, quenched by slow addition of water (20 mL), and concentrated to an oil under reduced pressure. The residue was dissolved in ethyl acetate and the organic layer was washed with 10 % aqueous HCl, water, saturated sodium bicarbonate, water, and brine, then concentrated to afford a colorless oil. Suspension in hexanes induced precipitation.

**Yield:** 14.4 g (27.41 mmol, 42 %). **$^1$H NMR** (300 MHz, DMSO-d$_6$) δ 10.65 (s, 1H), 8.03 (s, 1H), 6.45 (br s, 2H), 5.91 (d, J = 6.14 Hz, 1H), 5.60 (d, J = 6.66 Hz, 1H), 4.81 (br d, J = 6.78 Hz, 1H), 4.14–4.35 (m, 2H), 3.87 (br s, 2H), 3.33 (s, 1H), 0.93–1.17 (m, 30H). **$^{13}$C NMR** (75 MHz, DMSO-d$_6$) δ 157.2, 154.1, 151.8, 137.0, 136.8, 129.6, 129.3, 117.6, 117.4, 86.2, 86.1, 85.3, 81.1, 77.4, 77.2, 75.3, 75.2, 61.5, 61.3, 17.8, 17.6, 17.4, 16.9, 13.4, 13.0, 12.8, 12.7, 12.5. **LC-MS:** m/z 526.2 $[M+H]^+$.

**N-(9-((6aS,8R,9R,9aR)-9-hydroxy-2,2,4,4-tetraisopropyltetrahydro-6H-furo[3,2-f][1,3,5,2,4]trioxadisilocin-8-yl)-6-oxo-6,9-dihydro-1H-purin-2-yl)isobutyramide *(26)***

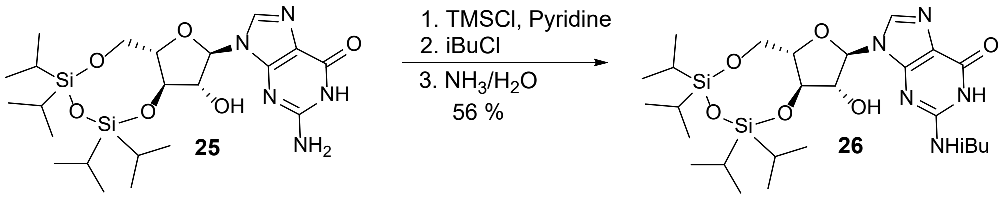


Compound **25** (14.20 g, 27.10 mmol) was dissolved in pyridine (100 mL) under nitrogen, cooled in an ice bath, and trimethylsilyl chloride (13.20 mL, 135 mmol, 5.0 equiv) was added dropwise. The ice bath was removed, and the mixture was stirred for 1 h at rt. The reaction was re-cooled and isobutyryl chloride (13.40 g, 135 mmol, 5.0 equiv) was added dropwise; the mixture was warmed to rt and stirred overnight. The reaction was cooled, water (40 mL) was added dropwise keeping the temperature below 7 °C, and the mixture was stirred at rt for 1 h. After cooling again, ammonium hydroxide (55 mL) was added dropwise and stirred for 30 min. Volatiles were partially removed at rt. The mixture was diluted with ethyl acetate and washed successively with water, saturated sodium bicarbonate, and brine; the organic layer was dried over sodium sulfate, filtered, and concentrated. Purification by Biotage chromatography (Si, 100 g column; dichloromethane/methanol 97:3 containing 1 % triethylamine) afforded a white solid.

**Yield:** 9.0 g (15.10 mmol, 56 %). **$^{1}$H NMR** (300 MHz, $CDCl_3$) δ 12.14 (br s, 1H), 10.69 (br s, 1H), 7.82 (s, 1H), 7.25–7.31 (m, 1H), 6.09 (br s, 1H), 5.72 (d, J = 6.02 Hz, 1H), 5.02 (br s, 2H), 4.41 (t, J = 8.45 Hz, 1H), 4.01–4.17 (m, 2H), 3.90–4.01 (m, 2H), 2.61–2.95 (m, 2H), 2.05 (s, 1H), 0.94–1.30 (m, 44H). **$^{13}$C NMR** (75 MHz, DMSO-$d_6$) δ 180.6, 155.3, 149.9, 149.4, 148.6, 148.5, 138.9, 121.7, 120.9, 86.3, 81.3, 77.4, 75.0, 61.6, 61.3, 46.4, 35.2, 26.4, 19.4, 17.7, 17.6, 17.4, 17.3, 13.5, 13.4, 13.0, 12.9, 12.8, 12.7, 12.5. **LC-MS:** m/z 596.3 $[M+H]^+$.

**O-((6aS,8R,9R,9aS)-8-(2-isobutyramido-6-oxo-1,6-dihydro-9H-purin-9-yl)-2,2,4,4-tetraiso propyltetrahydro-6H-furo[3,2-f][1,3,5,2,4]trioxadisilocin-9-yl) O-(p-tolyl) carbonothioate *(27)***

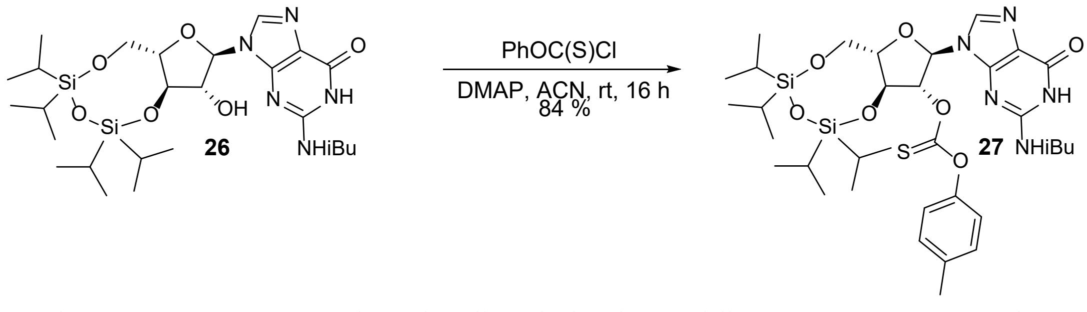


Compound **26** (7.80 g, 13.1 mmol) and 4-dimethylaminopyridine (3.20 g, 26.2 mmol, 2.0 equiv) were dissolved in anhydrous acetonitrile (131 mL); anhydrous tetrahydrofuran (50 mL) was added to aid dissolution. O-4-Methylphenyl chlorothioformate (2.69 mL, 14.4 mmol, 1.2 equiv) was added slowly, and the mixture was stirred at rt for 16 h (monitored by TLC, dichloromethane/methanol 95:5). Solvent was removed, and the residue was partitioned between ethyl acetate and water. The aqueous layer was extracted with ethyl acetate; combined organics were washed with 10 % aqueous HCl, water, saturated sodium bicarbonate, water, and brine; dried over magnesium sulfate; filtered; and concentrated. Purification by Biotage chromatography (Si, 100 g column, 0–3 % methanol/dichloromethane) afforded a white solid.

**Yield:** 8.24 g (11.0 mmol, 84 %). **$^1$H NMR** (300 MHz, $CDCl_3$) δ 12.10 (br s, 1H), 8.94 (br s, 1H), 8.02 (s, 1H), 7.13–7.31 (m, 3H), 6.89–7.00 (m, 2H), 6.44 (dd, J = 3.33, 5.12 Hz, 1H), 6.09 (d, J = 3.46 Hz, 1H), 4.78 (dd, J = 5.12, 7.42 Hz, 1H), 4.22–4.46 (m, 1H), 3.89–4.16 (m, 2H), 2.56–2.74 (m, 1H), 2.36 (s, 3H), 2.05 (s, 1H), 0.97–1.28 (m, 45H). **$^{13}$C NMR** (75 MHz, DMSO-$d_6$) δ 194.3, 180.5, 176.9, 155.2, 151.2, 149.3, 149.0, 148.7, 148.6, 139.0, 136.8, 130.7, 130.4, 122.0, 121.4, 121.0, 87.4, 83.8, 81.1, 72.7, 60.5, 46.5, 35.2, 26.4, 20.8, 19.5, 19.4, 19.3, 19.2, 17.8, 17.7, 17.6, 17.5, 17.4, 17.3, 17.2, 17.1, 13.6, 13.4, 13.0, 12.5, 12.4, 12.3. **LC-MS:** m/z 744.3 $[M–H]^-$.

**N-(6-oxo-9-((6aS,8R,9aR)-2,2,4,4-tetraisopropyltetrahydro-6H-furo[3,2-f][1,3,5,2,4]trioxadisilocin-8-yl)-6,9-dihydro-1H-purin-2-yl)isobutyramide *(28)***

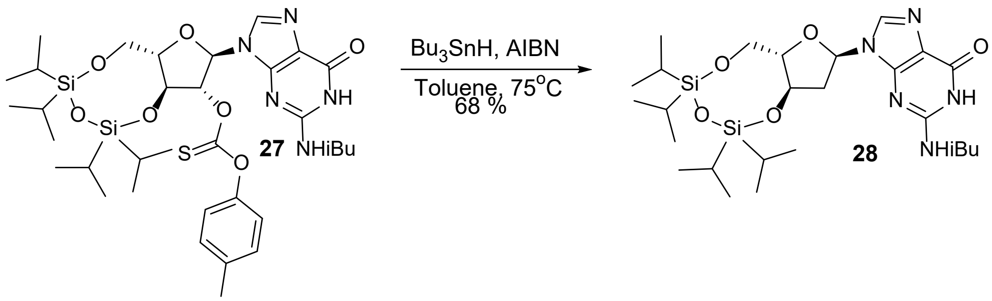


A degassed solution of **27** (6.67 g, 8.94 mmol) in toluene (140 mL) was heated to 80 °C under nitrogen. A separately degassed solution of azobisisobutyronitrile (0.267 g, 1.80 mmol, 0.20 equiv) and tributyltin hydride (24.10 mL, 89.4 mmol, 10 equiv) in toluene (40 mL) was added dropwise at 80 °C. The reaction was held at 80 °C for 1 h, cooled, and concentrated. TLC (ethyl acetate/hexanes 7:3) indicated conversion. Purification by Biotage chromatography (Si, 100 g column, 70 % ethyl acetate/hexanes) afforded a white solid.

**Yield:** 3.54 g (6.11 mmol, 68 %). $^{1}$**H NMR** (300 MHz, DMSO-$d_6$) δ 12.11 (br s, 1H), 11.65 (br s, 1H), 8.31 (s, 1H), 6.13–6.22 (m, 1H), 4.60 (br d, J = 6.01 Hz, 1H), 3.77–4.02 (m, 3H), 2.63–2.94 (m, 3H), 0.98–1.16 (m, 34H). $^{13}$**C NMR** (75 MHz, DMSO-$d_6$) δ 180.6, 155.3, 149.1, 148.5, 138.2, 121.0, 84.8, 82.2, 72.4, 62.7, 38.6, 35.2, 19.4, 19.3, 17.7, 17.4, 17.3, 17.2, 13.2, 13.1, 12.7, 12.4. **LC-MS:** m/z 578.3 $[M–H]^-$.

**N-(9-((2R,4R,5S)-4-hydroxy-5-(hydroxymethyl)tetrahydrofuran-2-yl)-6-oxo-6,9-dihydro-1H-purin-2-yl)isobutyramide *(29)***

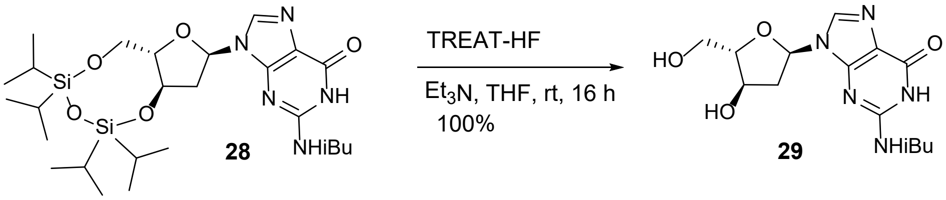


Triethylamine (2.13 mL, 15.40 mmol, 2.5 equiv) was added to a solution of **28** (3.54 g, 6.11 mmol) in tetrahydrofuran (30 mL) at 0 °C under nitrogen. Triethylamine trihydrofluoride (4.98 mL, 30.5 mmol, 5.0 equiv) was added slowly at 0 °C; the mixture was warmed to rt and stirred for 16 h. Solvent was removed, and the residue was purified by a silica gel plug (50 g, 5–10 % methanol/dichloromethane) to give a white solid as the TEA·HF salt, with corresponding TEA signals in the NMR

**Yield:** 2.05 g (6.01 mmol, 100%). $^{1}$**H NMR** (300 MHz, DMSO-$d_6$) δ 11.78 (br s, 1H), 8.24 (s, 1H), 6.21 (dd, J = 2.51, 7.72 Hz, 1H), 5.99 (s, 2H), 4.29–4.38 (m, 1H), 4.10–4.17 (m, 1H), 3.45 (d, J = 4.49 Hz, 2H), 3.18 (s, 1H), 3.07 (q, J = 7.30 Hz, 9H, TEA–HF), 2.67–2.86 (m, 2H), 2.50–2.53 (m, 1H), 2.32 (br d, J = 14.18 Hz, 1H), 1.11–1.23 (m, 20H, TEA–HF). $^{13}$**C NMR** (75 MHz,

DMSO-$d_6$) δ 180.6, 166.0, 155.4, 148.8, 148.4, 138.6, 120.5, 89.2, 84.0, 71.1, 62.1, 45.8, 35.2, 19.3, 8.9. **LC-MS:** m/z 338.2 $[M+H]^+$.

**N-(9-((2R,4R,5S)-5-((bis(4-methoxyphenyl)(phenyl)methoxy)methyl)-4-hydroxytetrahydro furan-2-yl)-6-oxo-6,9-dihydro-1H-purin-2-yl)isobutyramide** ***(30)***

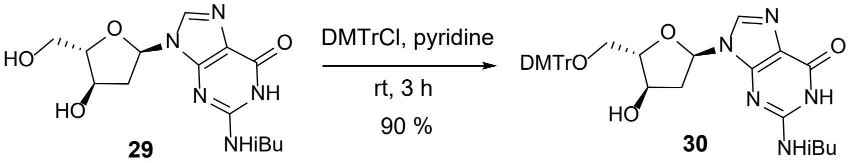


4,4'-Dimethoxytrityl chloride (2.41 g, 7.12 mmol, 1.2 equiv) was added to a solution of **29** (2.0 g, 5.93 mmol) in pyridine (30 mL) at rt and stirred for 2 h. The reaction was quenched with methanol (2 mL), diluted with water and ethyl acetate, and the aqueous phase was extracted with ethyl acetate. Combined organics were washed with water, saturated sodium bicarbonate, and brine; dried over sodium sulfate; filtered; and concentrated. The crude residue was dissolved in dichloromethane and purified by Biotage chromatography (Si, 100 g column, 0–5 % methanol/dichloromethane) to give a white solid.

**Yield:** 3.40 g (5.32 mmol, 90 %). **$^1$H NMR** (300 MHz, $CDCl_3$) δ 11.83–12.18 (m, 1H), 9.10–9.44 (m, 1H), 8.05 (s, 1H), 7.40 (d, J = 7.27 Hz, 2H), 7.15–7.34 (m, 8H), 6.81 (d, J = 8.80 Hz, 4H), 6.17–6.25 (m, 1H), 5.29 (br s, 1H), 4.44–4.54 (m, 2H), 3.77 (s, 6H), 3.27 (dd, J = 4.08, 10.19 Hz, 1H), 3.14 (dd, J = 3.55, 10.19 Hz, 1H), 2.63–2.90 (m, 3H), 1.18–1.31 (m, 6H). **$^{13}$C NMR** (75 MHz, $CDCl_3$) δ 178.9, 158.5, 155.3, 147.4, 147.3, 144.7, 139.1, 135.9, 135.7, 130.1, 128.1, 127.9, 126.9, 120.7, 113.2, 88.5, 86.5, 86.0, 72.8, 64.3, 55.3, 41.0, 36.4, 19.0, 18.9. **LC-HRMS (ESI$^-$):** m/z 638.2617 $[M-H]^-$

**(2S,3R,5R)-2-((bis(4-methoxyphenyl)(phenyl)methoxy)methyl)-5-(2-isobutyramido-6-oxo-1,6-dihydro-9H-purin-9-yl)tetrahydrofuran-3-yl(2-cyanoethyl) diisopropylphosphoramidite *(31)***

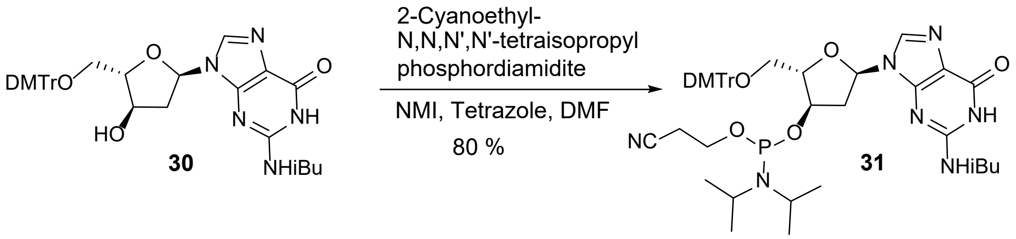


1*H*-Tetrazole (0.294 g, 4.25 mmol, 0.80 equiv) and 1-methylimidazole (0.105 mL, 1.33 mmol, 0.25 equiv) were added to a solution of **30** (3.40 g, 5.32 mmol) in DMF (40 mL) at rt under nitrogen. 2-Cyanoethyl-N,N,N',N'-tetraisopropylphosphorodiamidite (2.53 mL, 7.97 mmol, 1.5 equiv) was added dropwise, and the mixture was stirred at rt for 16 h. Water (1 mL) was added to quench the reaction. A 3:1 mixture of toluene/hexanes (80 mL) was added, and the mixture was washed four times with 3:2 (v/v) DMF/water (50 mL). The organic phase was washed with saturated sodium bicarbonate and brine, dried over sodium sulfate, and concentrated to a white foam. Purification by a silica gel plug (50 g, 100 % ethyl acetate) afforded a white amorphous solid.

**Yield:** 3.54 g (4.21 mmol, 80 %). **$^{1}$H NMR** (300 MHz, $CDCl_3$) δ 8.14 (d, J = 9.78 Hz, 1H), 7.40–7.47 (m, 2H), 7.19–7.38 (m, 8H), 6.84 (dd, J = 1.97, 8.89 Hz, 4H), 6.20–6.33 (m, 1H), 4.53–4.67 (m, 2H), 4.44 (br s, 1H), 3.72–3.82 (m, 7H), 3.38–3.67 (m, 5H), 3.09–3.34 (m, 2H), 2.72–2.96 (m, 2H), 2.50–2.72 (m, 3H), 2.32–2.47 (m, 1H), 1.04–1.31 (m, 23H). **$^{31}$P NMR** (121 MHz, $CDCl_3$) δ 149.92 (s, 1P), 148.82 (s, 1P). **LC-HRMS (ESI$^{-}$):** m/z 838.3691 $[M-H]^{-}$

**Synthesis of dx U phosphoramidite.** The synthesis of dx U phosphoramidite has not been previously reported. The synthesis of dx T amidite has been described[5] starting from the known dx T nucleoside[6-8]. This methodology relies on activating the 3′-OH group, followed by intramolecular cyclization to form the 2,3′-anhydronucleoside, which is then opened with alkali to achieve the C3′ inversion. Here, we follow the same route with modifications.

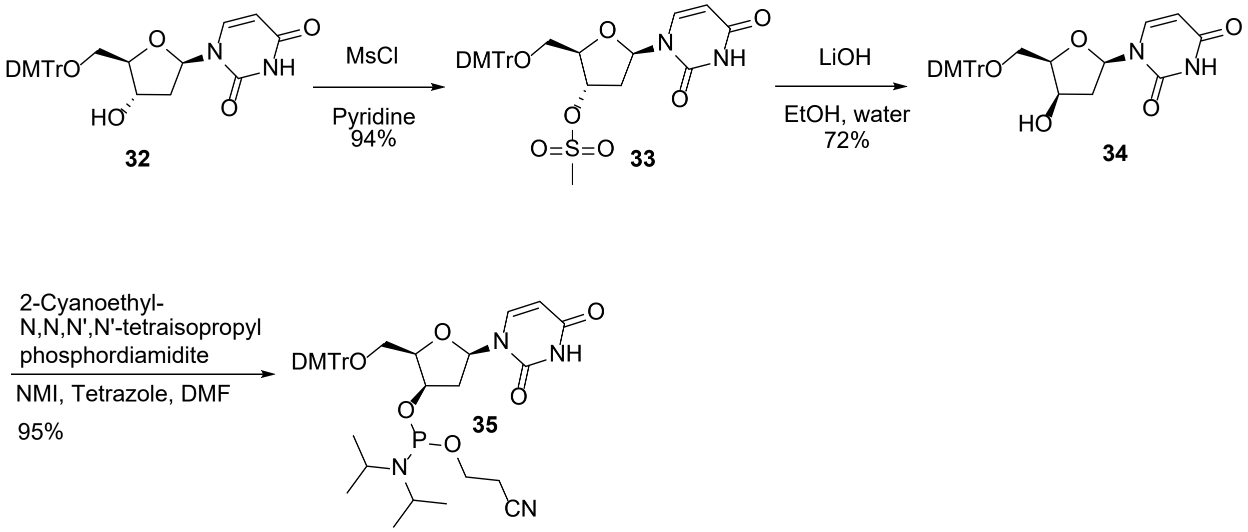


**(2R,3S,5R)-2-((Bis(4-methoxyphenyl)(phenyl)methoxy)methyl)-5-(2,4-dioxo-3,4-dihydropyrimidin-1(2H)-yl)tetrahydrofuran-3-yl methanesulfonate (*33*)**

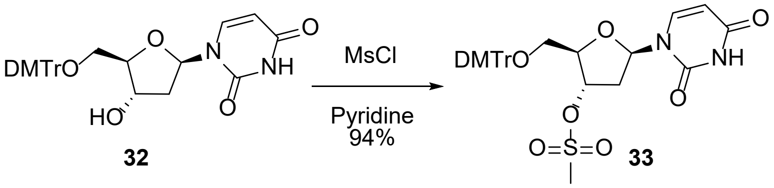


A solution of commercially available 5'-O-(4,4'-dimethoxytrityl)-2'-deoxyuridine **32** (18.0 g, 33.93 mmol) in dry pyridine (135.5 mL) was cooled to 0 °C (ice bath), and methanesulfonyl chloride (3.17 mL, 40.71 mmol, 1.20 equiv) was added dropwise over 20 min. The bath was removed, and the mixture was stirred at rt for 2 h. The reaction was diluted with saturated aqueous sodium carbonate and extracted with ethyl acetate. Combined organic layers were washed with brine and concentrated to a crude solid. Purification by Biotage (Si, 220 g column, 0–50 % ethyl acetate/hexanes) afforded the product as a white solid.

**Yield:** 19.50 g (32.04 mmol, 94 %). **$^1$H NMR** (300 MHz, $CDCl_3$) δ 8.80 (s, 1H), 8.63 (br d, J = 4.40 Hz, 1H), 7.64–7.72 (m, 1H), 7.16–7.37 (m, 11H), 6.85 (d, J = 8.80 Hz, 4H), 6.35 (dd, J = 6.01, 7.72 Hz, 1H), 5.31–5.44 (m, 2H), 4.31 (d, J = 2.33 Hz, 1H), 3.80 (s, 6H), 3.51 (d, J = 2.69 Hz, 2H), 3.01 (s, 3H), 2.70 (ddd, J = 1.84, 5.70, 14.36 Hz, 1H), 2.34–2.48 (m, 1H). **$^{13}$C NMR** (75 MHz, $CDCl_3$) δ 162.7, 158.9, 150.1, 149.8, 144.0, 139.6, 136.0, 134.9, 130.1, 128.1, 127.3, 123.7, 113.4, 102.8, 87.5, 84.7, 83.9, 79.4, 62.6, 55.3, 38.8, 38.6. **LC-MS:** m/z 607.1 $[M–H]^-$.

**1-((2R,4R,5R)-5-((Bis(4-methoxyphenyl)(phenyl)methoxy)methyl)-4-hydroxytetrahydrofuran-2-yl)pyrimidine-2,4(1H,3H)-dione (*34*)**

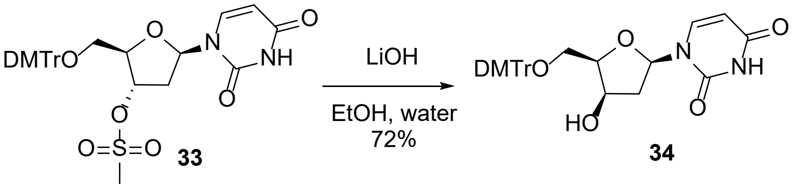


To a solution of **33** (27.0 g, 44.36 mmol) in ethanol/water (600 mL, 2:1 v/v) was added LiOH (3.19 g, 133.08 mmol, 3.00 equiv). The mixture was heated at 80 °C for 3 h, ethanol was removed, and the residue was extracted with dichloromethane. The organic layer was concentrated and purified by Biotage (Si, 100 g column, 0–100 % ethyl acetate/hexanes) to give the product as a white solid.

**Yield:** 16.91 g (31.87 mmol, 72 %). **$^1$H NMR** (300 MHz, $CDCl_3$) δ 7.79 (d, J = 8.17 Hz, 1H), 7.44 (d, J = 7.18 Hz, 2H), 7.15–7.37 (m, 8H), 6.83 (d, J = 8.89 Hz, 4H), 6.14 (br d, J = 6.64 Hz, 1H), 5.53 (d, J = 8.08 Hz, 1H), 4.36–4.44 (m, 1H), 4.00–4.05 (m, 1H), 3.77 (s, 6H), 3.58–3.73 (m, 1H), 3.37–3.56 (m, 1H), 2.52 (ddd, J = 5.03, 7.88, 14.92 Hz, 1H), 2.20 (br d, J = 15.08 Hz, 1H). **$^{13}$C NMR** (75 MHz, $CDCl_3$) δ 164.1, 158.7, 158.7, 150.9, 144.4, 141.4, 135.5, 135.4, 130.0, 130.0,

129.2, 128.1, 128.0, 127.8, 127.8, 127.0, 113.3, 113.2, 101.4, 86.9, 85.8, 83.6, 70.7, 61.9, 55.2, 41.1. **LC-HRMS (ESI$^-$):** m/z 529.1975 [M−H]$^-$

**(2R,3R,5R)-2-((Bis(4-methoxyphenyl)(phenyl)methoxy)methyl)-5-(2,4-dioxo-3,4-dihydropyrimidin-1(2H)-yl)tetrahydrofuran-3-yl (2-cyanoethyl) diisopropylphosphoramidite (*35*)**

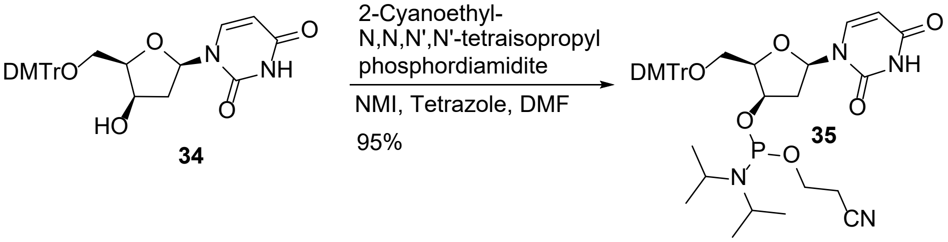


1*H*-Tetrazole (832.9 mg, 12.1 mmol, 0.80 equiv) and 1-methylimidazole (0.301 mL, 3.77 mmol, 0.25 equiv) were added to a solution of **34** (8 g, 15.1 mmol, 1 equiv) in DMF (200 mL) at rt under nitrogen. 2-Cyanoethyl-N,N,N',N'-tetraisopropylphosphorodiamidite (6.23 mL, 19.6 mmol, 1.50 equiv) was added dropwise, and the reaction was stirred at rt for 4 h. Water (1 mL) was then added to quench the reaction. The mixture was diluted with 3:1 toluene/hexanes (80 mL) and washed four times with 3:2 (v/v) DMF/water (50 mL), then with saturated sodium bicarbonate and brine. The upper organic phase was dried over sodium sulfate and concentrated under reduced pressure to a white foam. The residue was dissolved in dichloromethane, loaded onto a silica plug, and eluted with 7:3 ethyl acetate/hexanes to afford the product as a white amorphous solid.

**Yield:** 10.50 g (14.37 mmol, 95 %). **$^1$H NMR** (300 MHz, $CDCl_3$) δ 8.43–8.63 (m, 1H), 7.63 (d, J = 8.12 Hz, 1H), 7.42–7.49 (m, 2H), 7.16–7.38 (m, 9H), 6.77–6.86 (m, 4H), 6.19 (t, J = 6.97 Hz, 1H), 5.56 (dd, J = 2.15, 8.17 Hz, 1H), 4.40–4.50 (m, 1H), 4.25–4.31 (m, 1H), 3.79 (d, J = 3.05 Hz, 7H), 3.29–3.64 (m, 6H), 2.49–2.77 (m, 2H), 2.20–2.45 (m, 2H), 1.09 (t, J = 6.91 Hz, 6H), 0.95 (br d, J = 11.04 Hz, 3H), 0.92 (br d, J = 11.04 Hz, 3H). **$^{31}$P NMR** (121 MHz, $CDCl_3$) δ 151.23 (s, 1P), 147.68 (s, 1P). **LC-HRMS (ESI$^-$):** m/z 729.3052 [M−H]$^-$

**Synthesis of dx C phosphoramidite.** The synthesis of dx C phosphoramidite has been reported, though yields and selectivity vary depending on the route.[9-11] Reported methods start from $N^4$-benzoylcytidine and differ mainly in how the 3′-carbon inversion and subsequent phosphitylation are achieved. Here, we describe an alternative synthesis from dx U via a U→C conversion through a 4-triazolyl intermediate, providing efficient access to the target amidite.

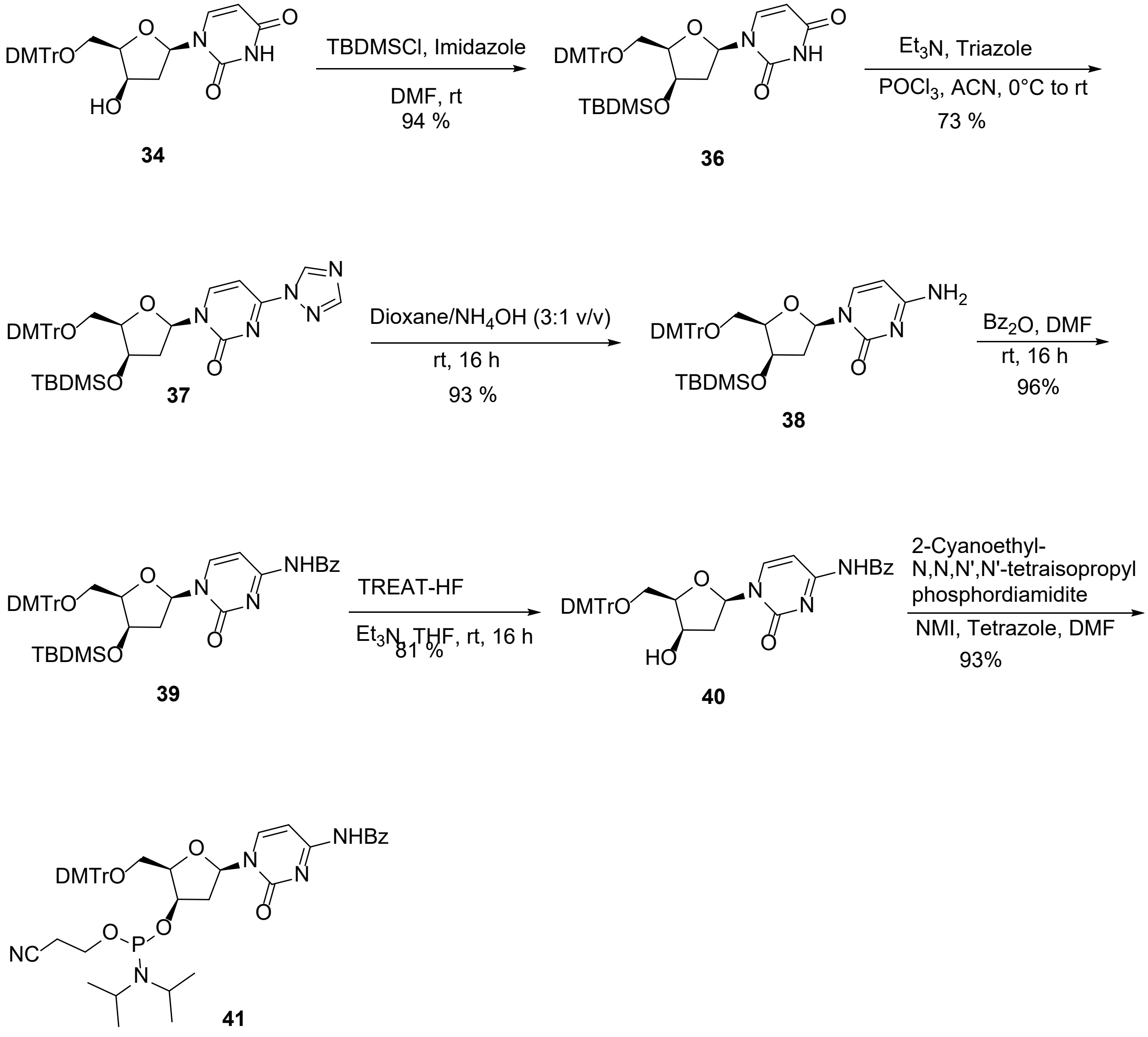

**1-((2R,4R,5R)-5-((Bis(4-methoxyphenyl)(phenyl)methoxy)methyl)-4-((tert-butyldimethylsilyl)oxy)tetrahydrofuran-2-yl)pyrimidine-2,4(1H,3H)-dione (*36*)**

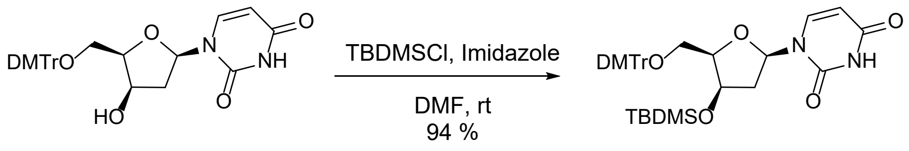


A solution of **34** (8.10 g, 15.27 mmol) and 1*H*-imidazole (2.08 g, 30.53 mmol, 2.0 equiv) in dry DMF (61.0 mL) was cooled to 0 °C (ice bath). tert-Butyldimethylsilyl chloride (2.53 g, 16.71 mmol, 1.1 equiv) was added dropwise over 10 min. The ice bath was removed, and the mixture was stirred at rt for 16 h. The reaction was diluted with saturated aqueous sodium carbonate and extracted with ethyl acetate. The combined organic layers were washed with brine and concentrated to a crude solid. Purification by Biotage chromatography (Si, 100 g column, 0–60 % ethyl acetate/hexanes) afforded the desired product as a white solid.

**Yield:** 9.24 g (14.3 mmol, 94 %). **$^1$H NMR** (300 MHz, $CDCl_3$) δ 8.59 (br s, 1H), 7.71 (d, J = 8.02 Hz, 1H), 7.54–7.61 (m, 2H), 7.30–7.50 (m, 9H), 6.87–6.99 (m, 4H), 6.27 (d, J = 6.46 Hz, 1H), 5.65 (dd, J = 2.15, 8.17 Hz, 1H), 4.19–4.38 (m, 3H), 3.89 (s, 6H), 3.63 (dd, J = 7.58, 10.55 Hz, 1H), 3.35 (dd, J = 3.10, 10.55 Hz, 1H), 2.63 (ddd, J = 4.58, 7.61, 14.74 Hz, 1H), 2.10–2.19 (m, 2H), 1.70 (s, 1H), 1.37 (t, J = 7.14 Hz, 1H), 1.00 (d, J = 11.67 Hz, 1H), 0.79 (s, 9H). **$^{13}$C NMR** (75 MHz, $CDCl_3$) δ 163.1, 158.6, 158.6, 150.2, 144.6, 141.1, 136.0, 135.8, 130.1, 130.0, 128.2, 127.9, 126.9, 113.2, 113.2, 101.0, 86.5, 86.1, 85.2, 71.2, 63.2, 55.2, 42.3, 25.5. **LC-MS:** m/z 643.3 $[M-H]^-$.

**1-((2R,4R,5R)-5-((Bis(4-methoxyphenyl)(phenyl)methoxy)methyl)-4-((tert-butyldimethylsilyl)oxy)tetrahydrofuran-2-yl)-4-(1H-1,2,4-triazol-1-yl)pyrimidine-2(1H)-one (*37*)**

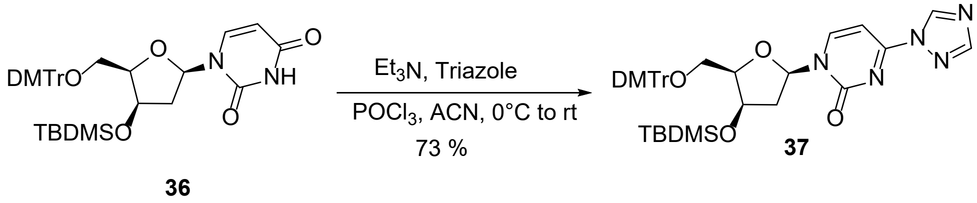


1*H*-1,2,4-Triazole (38.75 g, 561 mmol, 34 equiv) was suspended in anhydrous acetonitrile (200 mL), and phosphorus oxychloride (8.65 mL, 94.54 mmol, 5.7 equiv) was added dropwise over 40 min under nitrogen. The mixture was cooled (ice bath) and triethylamine (92.0 mL, 660 mmol, 7.0 equiv) was added dropwise over 15 min. The solution was stirred 40 min while warming to rt, cooled again to 0 °C, and nucleoside **36** (10.64 g, 16.50 mmol) in acetonitrile (20 mL) was added dropwise over 15 min. The reaction was stirred 16 h at rt, cooled, and quenched with water (50 mL). The mixture was concentrated, diluted with ethyl acetate, and washed with saturated sodium bicarbonate (2×), water, and brine. Concentration gave a yellow oil, which was purified by Biotage (Si, 100 g column, 0–100 % ethyl acetate/hexanes) to yield a white solid.

**Yield:** 8.40 g (12.07 mmol, 73 %). **$^1$H NMR** (300 MHz, $CDCl_3$) δ 9.45 (s, 1H), 8.37 (s, 1H), 8.30 (s, 1H), 8.29 (d, J = 7.77 Hz, 2H), 7.40–7.71 (m, 9H), 6.97–7.10 (m, 5H), 6.37 (d, J = 6.82 Hz, 1H), 4.59 (td, J = 2.95, 7.92 Hz, 1H), 4.47 (t, J = 3.46 Hz, 1H), 4.31 (q, J = 7.18 Hz, 1H), 3.99 (s, 5H), 3.76 (dd, J = 8.03, 10.64 Hz, 1H), 3.51 (dd, J = 2.96, 10.68 Hz, 1H), 2.80 (ddd, J = 4.26, 7.25, 14.88 Hz, 1H), 2.47 (d, J = 14.90 Hz, 1H), 2.23 (s, 1H), 1.37–1.48 (m, 2H), 0.75–0.81 (m, 8H).

**$^{13}$C NMR** (75 MHz, $CDCl_3$) δ 159.1, 158.6, 154.6, 153.9, 147.6, 144.6, 143.2, 135.9, 130.2, 128.2, 127.9, 127.0, 113.2, 93.4, 88.8, 86.6, 86.2, 71.5, 63.3, 60.4, 55.2, 42.1, 25.5, 17.7, 14.2.

**4-Amino-1-((2R,4R,5R)-5-((Bis(4-methoxyphenyl)(phenyl)methoxy)methyl)-4-((tert-butyldimethylsilyl)oxy)tetrahydrofuran-2-yl)pyrimidine-2(1H)-one (*38*)**

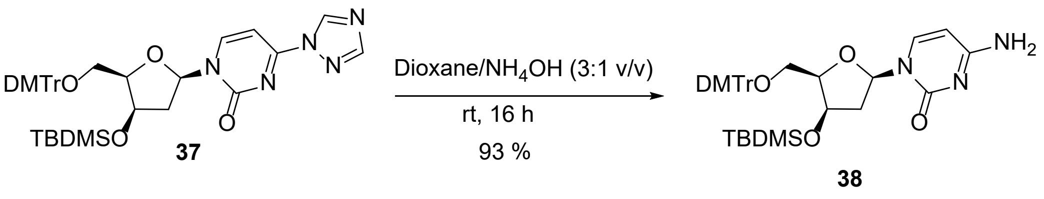


A solution of **37** (8.36 g, 12.02 mmol, 1 equiv) in dioxane/ammonium hydroxide (3:1, v/v; 80 mL) was stirred at rt for 16 h. The solvent was removed to give a white foam, which was purified by Biotage (Si, 100 g column, 0–20 % methanol/dichloromethane) to afford a white solid.

**Yield:** 7.20 g (11.18 mmol, 93 %). **$^{1}$H NMR** (300 MHz, DMSO-$d_6$) δ 7.52 (d, J = 7.36 Hz, 1H), 7.39–7.46 (m, 2H), 7.06–7.34 (m, 8H), 6.88 (d, J = 8.71 Hz, 6H), 6.04 (d, J = 6.91 Hz, 1H), 5.55 (d, J = 7.45 Hz, 1H), 4.19–4.36 (m, 2H), 4.03 (d, J = 7.09 Hz, 1H), 3.74 (s, 6H), 3.39 (s, 1H), 3.30 (br s, 1H), 2.94–3.19 (m, 1H), 2.51 (br s, 4H), 1.99 (s, 1H), 1.82 (br d, J = 14.54 Hz, 1H), 1.17 (t, J = 7.14 Hz, 1H), 0.61 (s, 9H), –0.18 (s, 3H), –0.22 (s, 3H). **$^{13}$C NMR** (75 MHz, DMSO-$d_6$) δ 165.5, 158.1, 155.0, 144.7, 140.8, 135.6, 135.5, 129.6, 127.7, 126.6, 113.1, 92.5, 85.8, 85.6, 83.8, 71.2, 63.4, 55.0, 40.3, 40.1, 39.8, 39.2, 39.0, 38.7, 25.2, 17.3, 14.0, –5.2, –5.9. **LC-MS:** m/z 642.3 $[M–H]^-$.

**N-(1-((2R,4R,5R)-5-((bis(4-methoxyphenyl)(phenyl)methoxy)methyl)-4-((tert-butyldimethylsilyl)oxy)tetrahydrofuran-2-yl)-2-oxo-1,2-dihydropyrimidin-4-yl)benzamide (*39*)**

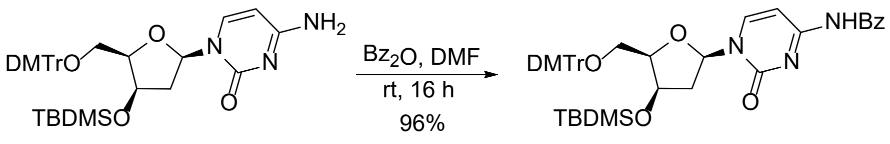

38 39

Benzoic anhydride (4.21 g, 18.59 mmol, 1.5 equiv) in anhydrous DMF (10 mL) was added dropwise to a solution of **38** (7.98 g, 12.39 mmol) in anhydrous DMF (50 mL) at rt. The mixture was stirred overnight, quenched with methanol (0.5 mL), and diluted with water/ethyl acetate (1:1, 100 mL). The aqueous phase was separated, and the organic layer was washed with saturated sodium bicarbonate (1 × 50 mL) and brine (1 × 50 mL), dried over sodium sulfate, filtered, and concentrated under reduced pressure. Purification by Biotage (Si, 100 g column, 0–3 % methanol/dichloromethane) afforded a white solid.

**Yield:** 8.92 g (11.93 mmol, 96 %). **$^1$H NMR** (300 MHz, $CDCl_3$) δ 8.05–8.21 (m, 3H), 7.76–7.82 (m, 1H), 7.66–7.73 (m, 4H), 7.40–7.62 (m, 10H), 7.03 (dd, J = 1.39, 8.84 Hz, 4H), 6.36 (d, J = 6.82 Hz, 1H), 4.43–4.57 (m, 2H), 4.30 (q, J = 7.12 Hz, 1H), 3.99 (d, J = 1.35 Hz, 6H), 3.77 (dd, J = 7.90, 10.50 Hz, 1H), 3.48 (dd, J = 3.01, 10.55 Hz, 1H), 3.13 (s, 1H), 3.06 (s, 1H), 2.70–2.83 (m, 1H), 2.43 (d, J = 14.81 Hz, 1H), 2.22 (s, 1H), 1.44 (t, J = 7.14 Hz, 1H), 0.80 (s, 9H). **$^{13}$C NMR** (75 MHz, $CDCl_3$) δ 162.5, 162.0, 158.6, 158.6, 145.7, 144.6, 136.0, 135.8, 133.1, 130.1, 130.0, 129.0, 128.2, 127.9, 127.6, 126.9, 113.2, 113.2, 88.1, 86.5, 85.9, 71.5, 63.4, 60.4, 55.3, 42.3, 36.4, 31.4, 25.4, 17.7, 14.2. **LC-MS:** m/z 746.3 $[M–H]^-$.

**N-(1-((2R,4R,5R)-5-((Bis(4-methoxyphenyl)(phenyl)methoxy)methyl)-4-hydroxytetrahydrofuran-2-yl)-2-oxo-1,2-dihydropyrimidin-4-yl)benzamide (*40*)**

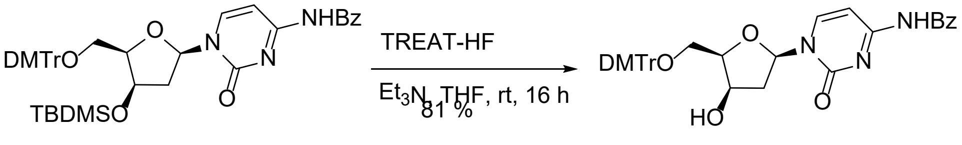


TEA (4.12 mL, 59.90 mmol, 5.0 equiv) was added to a solution of **39** (8.90 g, 11.90 mmol) in THF (50.0 mL) at 0 °C (ice bath) under nitrogen. Triethylamine trihydrofluoride (9.76 mL, 59.49 mmol, 5.0 equiv) was added slowly at 0 °C, then the reaction was warmed to rt and stirred for 16 h. The reaction was quenched by addition of TEA (6 mL), and the solvents were removed under reduced pressure. The residue was diluted with water/ethyl acetate (1:1, 100 mL). The aqueous layer was removed, and the organic layer was washed with saturated sodium bicarbonate (1 × 50.0 mL) and brine (1 × 50.0 mL), then dried over $Na_2SO_4$, filtered, and concentrated under reduced pressure. Purification by Biotage chromatography (Si, 100 g column, 0–3 % methanol/dichloromethane) afforded the desired product as a white solid.

**Yield:** 6.10 g (9.63 mmol, 81 %). **$^1$H NMR** (300 MHz, $CDCl_3$) δ 8.51–8.75 (m, 1H), 8.10–8.24 (m, 1H), 7.88 (br d, J = 7.36 Hz, 2H), 7.40–7.61 (m, 5H), 7.27–7.39 (m, 7H), 7.06–7.24 (m, 1H), 6.86 (d, J = 7.90 Hz, 4H), 6.14 (br t, J = 5.92 Hz, 1H), 4.45 (br s, 1H), 4.19–4.24 (m, 1H), 3.81 (s, 6H), 3.56–3.75 (m, 2H), 2.52–2.67 (m, 1H), 2.45 (br d, J = 13.37 Hz, 1H). **$^{13}$C NMR** (75 MHz, $CDCl_3$) δ 162.1, 158.8, 158.7, 144.2, 135.4, 135.4, 133.0, 130.0, 130.0, 129.0, 128.1, 127.6, 127.1, 113.4, 87.9, 87.1, 84.4, 71.1, 62.0, 55.3, 41.5. **LC-HRMS (ESI$^-$):** m/z 632.2398 $[M-H]^-$

**(2S,3R,5R)-5-(4-Benzamido-2-oxopyrimidin-1(2H)-yl)-2-((Bis(4-methoxyphenyl)(phenyl) methoxy)methyl)tetrahydrofuran-3-yl (2-cyanoethyl) diisopropylphosphoramidite (*41*)**

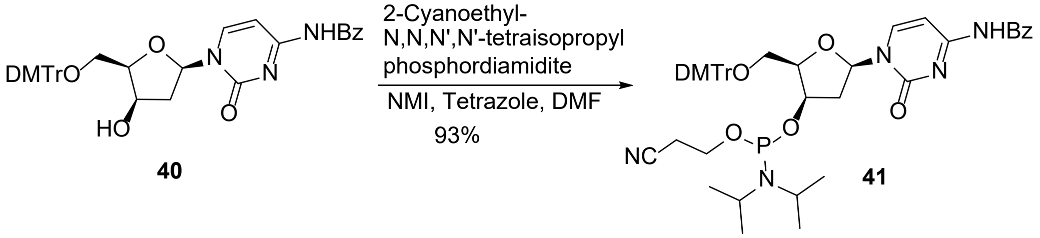


1*H*-Tetrazole (537.8 mg, 7.79 mmol, 0.8 equiv) and 1-methylimidazole (0.194 mL, 2.43 mmol, 0.25 equiv) were added to a solution of **40** (6.17 g, 9.74 mmol) in DMF (48 mL) under nitrogen at rt. 2-Cyanoethyl-N,N,N',N'-tetraisopropylphosphorodiamidite (4.64 mL, 14.60 mmol, 1.5 equiv) was added dropwise, and the reaction was stirred for 16 h at rt. Water (1 mL) was then added to quench the reaction. A 3:1 (v/v) toluene/hexanes mixture (80 mL) was added, and the mixture was washed four times with 3:2 (v/v) DMF/water (50 mL), then with saturated sodium bicarbonate and brine, dried over sodium sulfate, and concentrated. Purification by silica gel plug (Si, 50 g column, 80–100 % ethyl acetate/hexanes) afforded a white amorphous solid.

**Yield:** 7.58 g (9.09 mmol, 93 %). **$^1$H NMR** (300 MHz, $CDCl_3$) δ 8.60 (br s, 1H), 7.83–8.03 (m, 4H), 7.46–7.63 (m, 5H), 7.21–7.43 (m, 10H), 6.84 (dd, J = 1.80, 8.89 Hz, 4H), 6.14 (d, J = 6.55 Hz, 1H), 5.89 (s, 1H), 4.40–4.49 (m, 2H), 4.05–4.27 (m, 2H), 3.81 (d, J = 2.06 Hz, 6H), 3.41–3.70 (m, 5H), 3.20–3.38 (m, 3H), 2.67–2.79 (m, 3H), 2.55 (dt, J = 3.59, 6.15 Hz, 3H), 1.01 (d, J = 6.73 Hz, 6H), 0.89 (d, J = 6.73 Hz, 6H). **$^{31}$P NMR** (121 MHz, $CDCl_3$) δ 151.23 (s, 1P), 147.69 (s, 1P). **LC-HRMS (ESI$^-$):** m/z 832.3477 $[M-H]^-$

**Synthesis of dx A phosphoramidite.** The synthesis of dx A phosphoramidite has been reported previously.[9, 12] We followed the established Mitsunobu-based route for C3′ inversion of $N^6$-benzoyladenosine.[13, 14] In this approach, a Mitsunobu reaction with a carboxylic acid introduces the 3′-O-acyl group with stereochemical inversion, generating the xylo-intermediate. Subsequent phosphitylation afforded the target amidite.

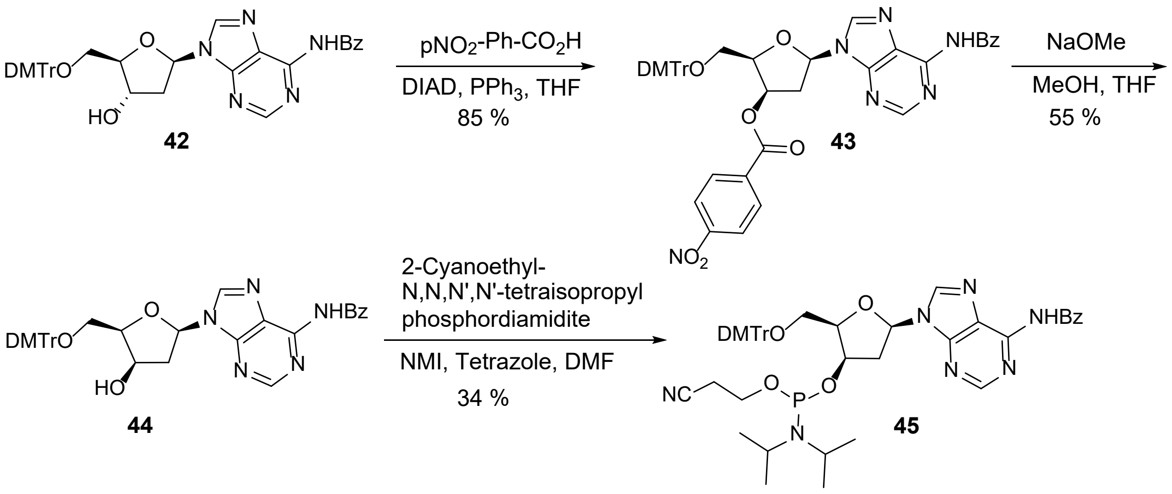


**(2R,3R,5R)-5-(6-Benzamido-9H-purin-9-yl)-2-((bis(4-methoxyphenyl)(phenyl)methoxy) methyl)tetrahydrofuran-3-yl 4-nitrobenzoate (*43*)**

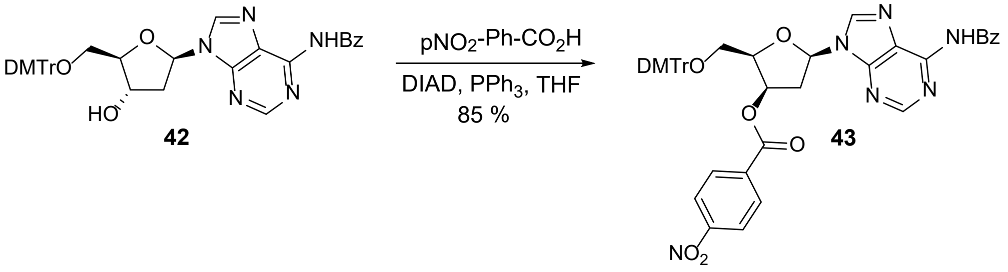


A solution of commercially available $N^6$-benzoyl-5'-O-(4,4'-dimethoxytrityl)-2'-deoxyadenosine **42** (8.00 g, 12.2 mmol) in THF (70 mL) was treated with 4-nitrobenzoic acid (4.07 g, 24.3 mmol) and triphenylphosphine (6.38 g, 24.3 mmol) at rt under nitrogen. The mixture was cooled to 0 °C (ice bath), and diisopropyl azodicarboxylate in THF (4.71 mL, 24.3 mmol, added in 10 mL THF) was added dropwise. The reaction was stirred for 30 min at 0 °C and then for 1 h at rt. The mixture was diluted with water, ethyl acetate, and saturated sodium bicarbonate. The aqueous layer was extracted with ethyl acetate, and the combined organic layers were washed with brine and concentrated. Purification by Biotage chromatography (Si, 50 g column, 0–100 % ethyl acetate/hexanes) afforded the product as an off-white foam.

**Yield:** 8.35 g (10.3 mmol, 85.1 %). **$^{1}$H NMR** (300 MHz, $CDCl_3$) δ 8.89 (s, 1H), 8.63 (s, 1H), 8.12–8.25 (m, 3H), 8.00 (dd, J = 1.54, 7.04 Hz, 2H), 7.54 (dd, J = 1.22, 7.10 Hz, 5H), 7.33–7.37 (m, 2H), 7.19 (br dd, J = 3.33, 5.50 Hz, 7H), 6.65–6.73 (m, 4H), 6.54 (dd, J = 2.62, 6.46 Hz, 1H), 5.95 (t, J = 3.33 Hz, 1H), 4.57–4.70 (m, 1H), 3.74 (s, 3H), 3.73 (s, 3H), 3.68 (dd, J = 5.63, 9.22 Hz, 1H), 3.46 (dd, J = 7.30, 9.22 Hz, 1H), 3.06 (s, 2H). **$^{13}$C NMR** (75 MHz, $CDCl_3$) δ 164.6, 163.3, 158.6, 158.5, 152.5, 151.4, 150.6, 149.5, 144.2, 140.7, 135.3, 135.0, 133.4, 133.2, 132.9, 131.9, 131.9, 131.9, 130.4, 130.0, 130.0, 129.9, 128.9, 127.9, 127.0, 123.7, 113.1, 113.1, 86.6, 84.6, 82.5, 73.4, 60.8, 55.2, 55.2, 55.1, 39.0. **LC–MS:** m/z 807.3 $[M+H]^+$.

**N-(9-((2R,4R,5R)-5-((bis(4-methoxyphenyl)(phenyl)methoxy)methyl)-4-hydroxytetrahydrofuran-2-yl)-9H-purin-6-yl)benzamide (*44*)**

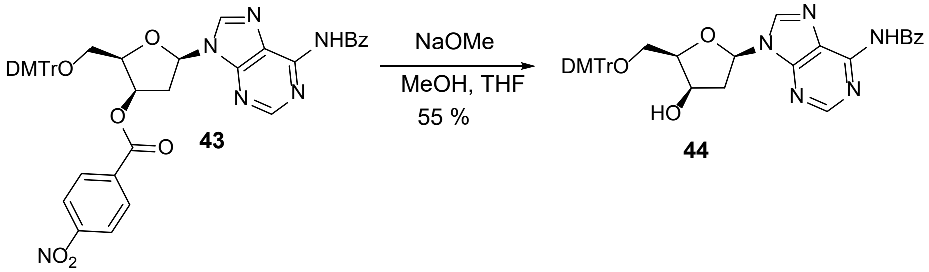


A solution of the 4-nitrobenzoate (8.35 g, 10.3 mmol) in THF (69.1 mL) was cooled to 0 °C (ice bath), and sodium methoxide in methanol (0.50 M, 20.7 mL, 10.3 mmol) was added. The reaction was stirred for 45 min at 0 °C, diluted with water and ethyl acetate, and the aqueous layer was extracted with ethyl acetate. Combined organic layers were washed with brine and concentrated. Purification by Biotage chromatography (Si, 220 g column, 0–100 % ethyl acetate/hexanes) afforded a white foam.

**Yield:** 3.71 g (5.64 mmol, 54.5 %). $^1$**H NMR** (300 MHz, DMSO-d$_6$) δ 11.20 (br s, 1H), 8.77 (s, 1H), 8.49 (br s, 1H), 8.05 (br d, J = 6.91 Hz, 2H), 7.48–7.70 (m, 3H), 7.32–7.46 (m, 2H), 7.14–7.32 (m, 7H), 6.83 (br d, J = 8.80 Hz, 2H), 6.79 (br d, J = 8.80 Hz, 2H), 6.52 (br d, J = 7.09 Hz, 1H), 5.50 (br s, 1H), 4.37 (br s, 1H), 4.25 (br s, 1H), 3.63–3.76 (m, 6H), 3.11–3.27 (m, 1H), 2.65–2.93 (m, 1H), 2.30–2.38 (m, 1H). $^{13}$**C NMR** (75 MHz, DMSO-d$_6$) δ 166.0, 158.5, 152.3, 152.0, 150.7, 145.4, 143.5, 136.2, 136.0, 133.8, 132.9, 130.2, 128.9, 128.2, 127.1, 126.1, 113.5, 85.9, 84.5, 83.7, 69.8, 63.7, 55.4, 41.1. **LC-HRMS (ESI$^-$):** m/z 656.2511 [M−H]$^-$

**(2R,3R,5R)-5-(6-Benzamido-9H-purin-9-yl)-2-((bis(4-methoxyphenyl)(phenyl)methoxy) methyl)tetrahydrofuran-3-yl (2-cyanoethyl) diisopropylphosphoramidite (*45*)**

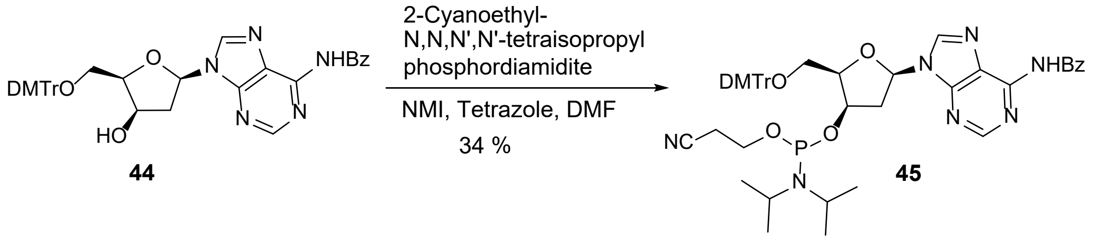


A solution of **44** (3.71 g, 5.64 mmol) in dry DMF (57.2 mL) was treated at rt under nitrogen with 1*H*-tetrazole (0.316 g, 4.51 mmol) and 1-methylimidazole (0.112 mL, 1.41 mmol), followed by dropwise addition of 2-cyanoethyl-N,N,N',N'-tetraisopropylphosphorodiamidite (2.69 mL, 8.46 mmol). The mixture was stirred for 90 min at rt. Water (1 mL) was added, then 3:1 toluene/hexanes (80 mL). The mixture was washed four times with 3:2 (v/v) DMF/water (50 mL), then with saturated sodium bicarbonate and brine, dried over sodium sulfate, and concentrated. Purification by Biotage chromatography (Si, 220 g column, 0–100 % ethyl acetate) afforded a white solid.

**Yield:** 1.65 g (1.93 mmol, 34.2 %). $^1$**H NMR** (300 MHz, CDCl$_3$) δ 8.93 (s, 1H), 8.75–8.85 (m, 1H), 8.32 (s, 1H), 8.01 (br d, J = 7.04 Hz, 2H), 7.43–7.69 (m, 5H), 7.27–7.41 (m, 5H), 7.23 (br d, J = 6.78 Hz, 2H), 6.76–6.89 (m, 4H), 6.49–6.64 (m, 1H), 4.57 (br s, 1H), 4.36–4.50 (m, 1H), 3.72–

3.89 (m, 6H), 3.51–3.68 (m, 2H), 3.18–3.50 (m, 4H), 2.67 (s, 2H), 2.25–2.50 (m, 2H), 0.98–1.18 (m, 6H), 0.81–0.97 (m, 6H). **$^{31}$P NMR** (121 MHz, $CDCl_3$) δ 150.94 (s), 147.91 (s). **LC-HRMS (ESI$^-$):** m/z 856.3591 [M−H]$^-$.

**Synthesis of dx G phosphoramidite.** The synthesis of dx G phosphoramidite has been reported previously.[9, 15] We followed the established triflate-based route for C3′ inversion of $N^2$-isobutyrylguanosine.[16, 17] In this approach, the 3′-hydroxyl group is activated with trifluoromethanesulfonic anhydride ($Tf_2O$), promoting intramolecular attack and resulting in stereochemical inversion to form the xylo intermediate. Subsequent phosphitylation provided the target amidite.

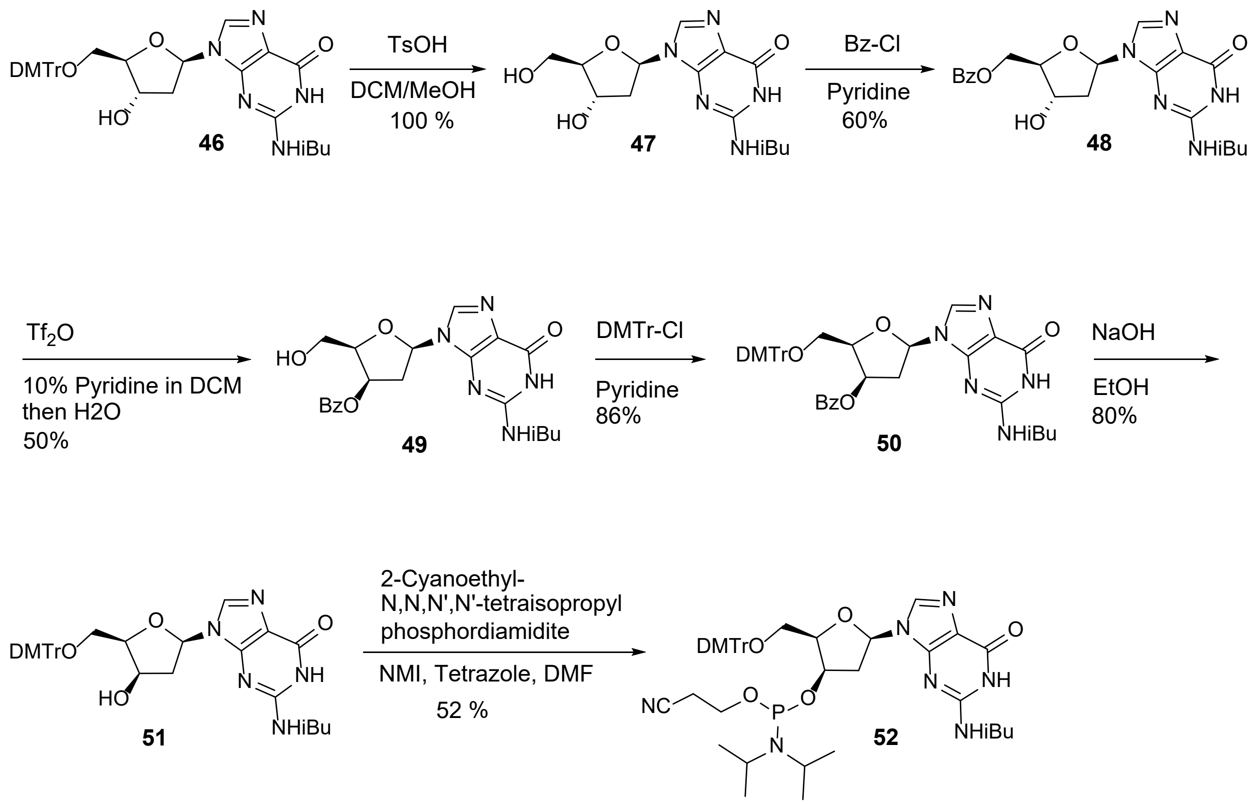

**N-(9-((2R,4S,5R)-4-hydroxy-5-(hydroxymethyl)tetrahydrofuran-2-yl)-6-oxo-6,9-dihydro-1H-purin-2-yl)isobutyramide (*47*)**

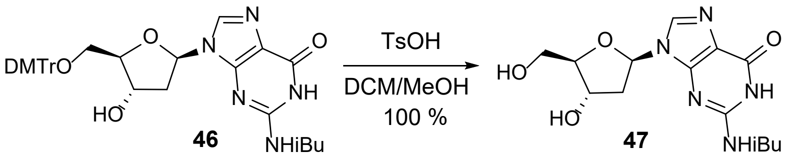


A solution of commercially available N$^2$-isobutyryl-5'-O-(4,4'-dimethoxytrityl)-2'-deoxyguanosine **46** (50.0 g, 78.2 mmol) in dichloromethane/methanol (3:1, 1560 mL) was cooled to 0 °C. p-Toluenesulfonic acid (17.8 g, 93.8 mmol) was added, and the orange reaction mixture was stirred at 0 °C for 60 min. Sodium carbonate (9.94 g, 93.8 mmol) was then added at 0 °C and stirring was continued until the orange color disappeared. The solvents were removed under reduced pressure. Dichloromethane was added to the residue, and the resulting white precipitate was collected and dried under high vacuum to give the crude product as a white solid.

**Yield:** 26.4 g (78.3 mmol, 100 %). **$^1$H NMR** (300 MHz, DMSO-d$_6$) δ 12.08 (br s, 1H), 11.68 (br s, 1H), 8.24 (s, 1H), 6.21 (dd, J = 6.02, 7.42 Hz, 1H), 5.31 (d, J = 3.84 Hz, 1H), 4.95 (t, J = 5.44 Hz, 1H), 4.37 (dd, J = 2.88, 5.70 Hz, 1H), 3.79–3.91 (m, 1H), 3.45–3.63 (m, 2H), 2.77 (quin, J = 6.82 Hz, 1H), 2.53–2.62 (m, 1H), 2.19–2.28 (m, 1H), 1.12 (d, J = 6.78 Hz, 6H). **$^{13}$C NMR** (75 MHz, DMSO-d$_6$) δ 180.1, 154.8, 148.3, 148.0, 137.8, 137.4, 120.1, 87.7, 82.9, 70.4, 61.4, 34.7, 18.8. **LC–MS:** m/z 338.1 [M+H]$^+$.

**((2R,3S,5R)-3-hydroxy-5-(2-isobutyramido-6-oxo-1,6-dihydro-9H-purin-9-yl)tetrahydrofuran-2-yl)methyl benzoate (*48*)**

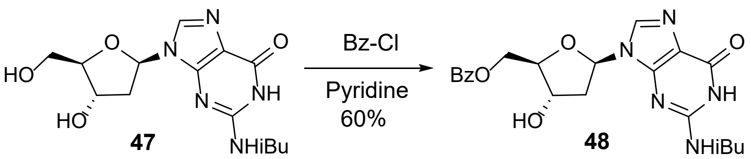

Crude **47** (26.4 g, 78.3 mmol) was suspended in pyridine (780 mL) under nitrogen. Benzoyl chloride (9.08 mL, 78.3 mmol) was added dropwise, and the reaction mixture was stirred at rt for 1 h. The solvents were removed under reduced pressure, and the residue was partitioned between dichloromethane and water. The organic layer was collected, washed with water (3×) and brine, dried over sodium sulfate, and concentrated. Purification by Biotage chromatography (Si, 330 g column, 0–10 % methanol/dichloromethane) afforded the desired product as a white solid.

**Yield:** 20.7 g (46.9 mmol, 59.9 %). **$^1$H NMR** (300 MHz, DMSO-d$_6$) δ 12.08 (br s, 1H), 11.64 (br s, 1H), 8.19 (s, 1H), 7.85–8.01 (m, 2H), 7.62–7.73 (m, 1H), 7.47–7.57 (m, 2H), 6.26 (t, J = 6.66 Hz, 1H), 5.54 (d, J = 4.10 Hz, 1H), 4.54–4.62 (m, 1H), 4.37–4.54 (m, 2H), 4.05–4.19 (m, 1H), 2.76 (quin, J = 6.56 Hz, 1H), 2.39 (ddd, J = 4.42, 6.43, 13.35 Hz, 1H), 1.12 (d, J = 6.78 Hz, 6H). **$^{13}$C NMR** (75 MHz, DMSO-d$_6$) δ 180.6, 166.0, 155.3, 148.9, 148.6, 137.9, 133.9, 129.8, 129.6, 129.2, 120.9, 84.6, 83.4, 70.8, 64.9, 35.2, 19.3. **LC–MS:** m/z 442.2 [M+H]$^+$.

**(2R,3R,5R)-2-(hydroxymethyl)-5-(2-isobutyramido-6-oxo-1,6-dihydro-9H-purin-9-yl)tetrahydrofuran-3-yl benzoate (*49*)**

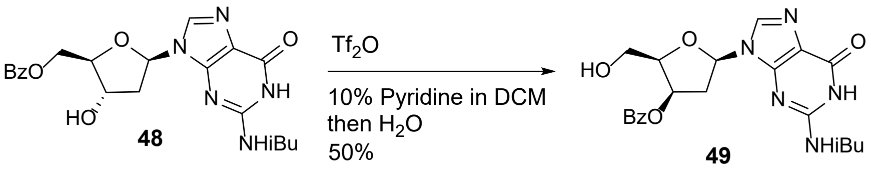


A solution of **48** (10.0 g, 22.7 mmol) in 10 % pyridine in dichloromethane (164 mL) was cooled to −35 °C (acetone/dry ice bath) under nitrogen. Trifluoromethanesulfonic anhydride (5.72 mL, 34.0 mmol) was added dropwise. After completion of addition, the reaction was warmed to 0 °C and stirred for 45 min, then water (4.92 mL, 273 mmol) was added. The mixture was allowed to warm to rt and stirred overnight. The solvents were removed under reduced pressure, and the residue was partitioned between water (150 mL) and ethyl acetate (150 mL). The white precipitate that formed was collected and dried under high vacuum to give the desired product as a white solid.

**Yield:** 5.24 g (11.9 mmol, 52.4 %). **$^{1}$H NMR** (300 MHz, DMSO-$d_6$) δ 12.05 (s, 1H), 11.71 (s, 1H), 8.19 (s, 1H), 7.78–7.89 (m, 2H), 7.60–7.75 (m, 1H), 6.25 (dd, J = 2.24, 7.62 Hz, 1H), 5.69 (t, J = 4.16 Hz, 1H), 4.28–4.41 (m, 1H), 3.67–3.86 (m, 2H), 3.01 (dq, J = 5.57, 7.57 Hz, 1H), 2.67–2.86 (m, 2H), 1.11 (d, J = 6.91 Hz, 6H). **$^{13}$C NMR** (75 MHz, DMSO-$d_6$) δ 180.6, 180.4, 165.3, 155.2, 148.6, 148.5, 137.4, 134.0, 129.9, 129.7, 129.6, 129.2, 129.1, 129.0, 120.9, 84.0, 83.7, 73.4, 59.6, 38.7, 35.2, 19.4, 19.3, 19.3. **LC–MS:** m/z 442.2 $[M+H]^+$.

**(2R,3R,5R)-2-((bis(4-methoxyphenyl)(phenyl)methoxy)methyl)-5-(2-isobutyramido-6-oxo-1,6-dihydro-9H-purin-9-yl)tetrahydrofuran-3-yl benzoate (*50*)**

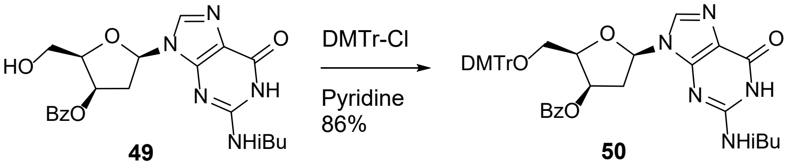


4,4'-Dimethoxytrityl chloride (3.68 g, 10.9 mmol) was added to a solution of [(2R,3R,5R)-2-(hydroxymethyl)-5-[2-(2-methylpropanoylamino)-6-oxo-1H-purin-9-yl]tetrahydrofuran-3-yl] benzoate (4.00 g, 9.06 mmol) in pyridine (30.2 mL) at rt, and the reaction was stirred for 2 h. The mixture was concentrated to an oil, and purification by Biotage chromatography (Si, 10 g column, 0–100 % ethyl acetate/hexanes) afforded the desired product as a white solid.

**Yield:** 5.79 g (7.78 mmol, 85.9 %). **$^{1}$H NMR** (300 MHz, DMSO-$d_6$) δ 12.05 (br s, 1H), 11.73 (br s, 1H), 7.95 (s, 1H), 7.57–7.74 (m, 3H), 7.40–7.51 (m, 2H), 7.26–7.37 (m, 2H), 7.10–7.25 (m, 7H), 6.73 (t, J = 8.58 Hz, 4H), 6.28 (dd, J = 2.37, 7.62 Hz, 1H), 5.82 (t, J = 4.22 Hz, 1H), 4.50–4.63 (m,

1H), 3.68 (d, J = 4.99 Hz, 6H), 3.27 (d, J = 6.02 Hz, 2H), 2.96–3.11 (m, 1H), 2.83–2.93 (m, 1H), 2.77 (quin, J = 6.78 Hz, 1H), 1.14 (m, 1H), 1.11 (dd, J = 0.90, 6.78 Hz, 5H). **$^{13}$C NMR** (75 MHz, DMSO-$d_6$) δ 180.6, 165.1, 158.5, 158.5, 155.2, 148.8, 148.5, 145.1, 137.3, 135.7, 135.7, 134.0, 130.1, 130.0, 129.6, 129.4, 129.1, 128.2, 128.0, 129.5, 127.1, 120.9, 113.5, 113.2, 86.2, 83.7, 81.8, 73.4, 61.9, 55.4, 37.9, 35.2, 19.3, 19.3. **LC–MS:** m/z 744.3 $[M+H]^+$.

**N-(9-((2R,4R,5R)-5-((bis(4-methoxyphenyl)(phenyl)methoxy)methyl)-4-hydroxytetrahydrofuran-2-yl)-6-oxo-6,9-dihydro-1H-purin-2-yl)isobutyramide (*51*)**

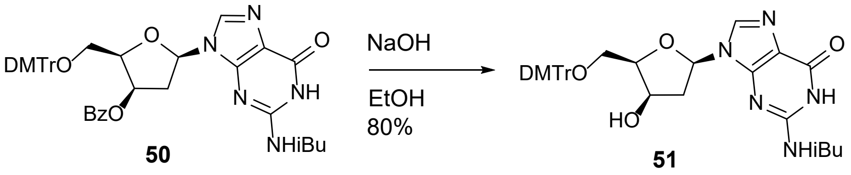


A solution of **50** (5.45 g, 7.33 mmol) in a 1:1:1 mixture of THF (54.5 mL), 1,4-dioxane (54.5 mL), and methanol (54.5 mL) was cooled to 0 °C. A solution of 1 N NaOH (54.5 mL) was added, and the reaction was stirred at 0 °C for 2 h. The mixture was diluted with ethyl acetate and water, and the aqueous fraction was extracted with ethyl acetate. Combined organic layers were washed with brine, dried over sodium sulfate, and concentrated. Purification by Biotage chromatography (Si, 10 g column, 0–5 % methanol/dichloromethane) afforded the desired product as a white solid.

**Yield:** 3.73 g (5.83 mmol, 79.6 %). **$^{1}$H NMR** (300 MHz, DMSO-$d_6$) δ 12.11 (br s, 1H), 11.76 (br s, 1H), 8.01 (s, 1H), 7.36–7.47 (m, 2H), 7.17–7.33 (m, 7H), 6.82 (dd, J = 8.96, 10.37 Hz, 4H), 6.22 (d, J = 6.53 Hz, 1H), 5.30 (d, J = 3.58 Hz, 1H), 4.34 (br d, J = 3.58 Hz, 1H), 4.15–4.27 (m, 1H), 3.72 (d, J = 2.56 Hz, 6H), 3.35–3.40 (m, 1H), 3.18 (dd, J = 2.69, 9.98 Hz, 1H), 2.77 (br d, J = 6.91 Hz, 2H), 2.28 (br d, J = 14.59 Hz, 1H), 1.12 (dd, J = 1.79, 6.78 Hz, 6H). **$^{13}$C NMR** (75 MHz, DMSO-$d_6$) δ 180.7, 158.5, 158.5, 155.4, 148.7, 148.5, 145.5, 138.4, 136.2, 136.0, 130.2, 128.2,

128.2, 127.1, 120.5, 113.5, 85.9, 84.5, 83.4, 69.7, 66.8, 63.7, 55.5, 55.4, 41.2, 35.2, 19.4, 19.3. **LC-HRMS (ESI⁻):** m/z 638.2617 [M−H]⁻

**(2R,3R,5R)-2-((bis(4-methoxyphenyl)(phenyl)methoxy)methyl)-5-(2-isobutyramido-6-oxo-1,6-dihydro-9H-purin-9-yl)tetrahydrofuran-3-yl (2-cyanoethyl) diisopropylphosphoramidite (*52*)**

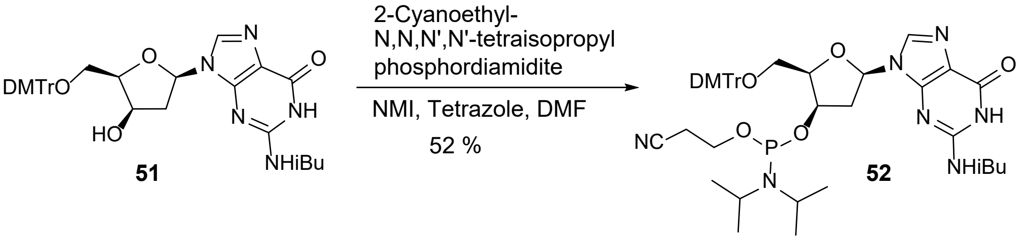


Compound **51** (3.00 g, 4.69 mmol) was dissolved in dry DMF (46.8 mL) under nitrogen. 1*H*-Tetrazole (0.263 g, 3.75 mmol) and 1-methylimidazole (0.0930 mL, 1.17 mmol) were added, followed by dropwise addition of 2-cyanoethyl-N,N,N',N'-tetraisopropylphosphorodiamidite (2.23 mL, 7.03 mmol). The reaction mixture was stirred at rt overnight. Water (1 mL) was added, followed by a 3:1 mixture of toluene/hexanes (80 mL). The mixture was washed four times with 3:2 (v/v) DMF/water (50 mL), then with saturated sodium bicarbonate and brine, dried over sodium sulfate, and concentrated to an oil. Purification by Biotage chromatography (Si, 50 g column, 0–100 % ethyl acetate) afforded the desired product as a white solid.

**Yield:** 2.03 g (2.42 mmol, 51.5 %). **$^{1}$H NMR** (300 MHz, $CDCl_3$) δ 12.03 (br s, 1H), 8.58–8.87 (m, 1H), 8.02 (d, J = 9.47 Hz, 1H), 7.41–7.53 (m, 2H), 7.30–7.38 (m, 4H), 7.27–7.30 (m, 1H), 7.24 (br d, J = 7.42 Hz, 2H), 6.69–6.88 (m, 4H), 6.11–6.26 (m, 1H), 4.52 (br d, J = 4.10 Hz, 1H), 4.30–4.42 (m, 1H), 3.20–3.71 (m, 7H), 2.23–2.88 (m, 6H), 1.15–1.24 (m, 10H), 1.06 (dd, J = 6.72, 15.30 Hz, 6H), 0.86–0.96 (m, 5H). **$^{31}$P** NMR (121 MHz, $CDCl_3$) δ 151.47 (s, 1P), 146.99 (s, 1P). **LC-HRMS (ESI⁻):** m/z 838.3693 [M−H]⁻

**Oligonucleotide Synthesis.** Oligonucleotide guide strands were synthesized on an Applied Biosystems 394 DNA/RNA Synthesizer at a 2 μmol scale using VIMAD UnyLinker solid support (200 μmol/g). Standard solid-phase protocols were employed, including deblocking with 3% dichloroacetic acid in dichloromethane, activation with 4,5-dicyanoimidazole (1 M) and N-methylimidazole (0.1 M) in acetonitrile, capping with 10% acetic anhydride in tetrahydrofuran (THF) and 10% N-methylimidazole in THF/pyridine, and thiolation using xanthane hydride (0.1 M) in pyridine:acetonitrile (3:2 v/v). Phosphoramidites were prepared at 0.1 M in acetonitrile with a 6-min coupling time, except for the 5′-POM-(*E*)-Vinylphosphonate (VP)-2′-MOE-T phosphoramidite, which was used at 0.15 M. It has been reported that the coupling kinetics of dx phosphoramidites are slower than those of the corresponding deoxyribo analogs[11]; however, under our experimental conditions these monomers coupled efficiently.

Following synthesis, oligonucleotides were cleaved and deprotected in 10% diethylamine/90% (9:1 $NH_4OH$:EtOH) for 36 h at rt. The crude products were filtered and purified by ion-pair reversed-phase HPLC (XBridge Prep C18, 5 μm, 19 × 250 mm) using a 1-90% acetonitrile gradient over 60 min in 5 mM tetrabutylammonium acetate (TBAA). Pure fractions were pooled and further purified by strong anion-exchange chromatography (SOURCE 30Q resin) using 100 mM ammonium acetate (Buffer A) and 1.5 M NaBr/100 mM ammonium acetate (Buffer B) in 3:7 acetonitrile:water. Final desalting was achieved on a C18 reverse-phase column, followed by drying in a vacuum concentrator. Passenger strands were synthesized in a similar manner on pre-loaded GalNAc support (154 μmol/g). After 5′-dimethoxytrityl removal, oligonucleotides were cleaved with 9:1 $NH_4OH$:EtOH for 36 h at rt, with subsequent purification steps as described above.

Large-scale (40 μmol) syntheses of guide and passenger strands were performed on an AKTA OligoPilot 10, following a reported procedure[18, 19], using NittoPhaseHL UnyLinker solid support (317 μmol/g) for guide strands and pre-loaded GalNAc support for passenger strands. Cleavage, deprotection, and purification followed the procedures outlined above.

**Dual-Luciferase Reporter Assay.** The effects of RNAi duplexes on on-target and off-target activity were assessed using the psiCHECK2™ reporter vector and Dual-Glo® Luciferase Assay System (Promega). On-target reporter vectors for *Ttr* and *ACTN1* contained a single fully complementary site to the respective siRNA guide strand, inserted into the 3′ UTR of the Renilla luciferase cassette. For *Ttr*, the sequence was 5′-AAAACAGTGTTCTTGCTCTATAA-3′, and for *ACTN1*, 5′-ATGTGTGTTTGCTAGCTCACTTA-3′. Off-target reporter plasmids contained four tandem seed-complementary sites separated by a 19-nt spacer sequence, also inserted into the 3′ UTR. The seed-complementary sequence for *Ttr* was 5′-GCTCTATAA-3′ with spacer 5′-TAATATTACATAAATAAAA-3′, and for *ACTN1*, the seed sequence was 5′-GCTCACTTA-3′ with spacer 5′-TAATATTACAAAAATAAAT-3′.

COS-7 cells (ATCC, Manassas, VA) were grown to near confluence, trypsinized, and seeded into 96-well plates in 100 μL of complete medium overnight prior to transfection. siRNA duplexes were co-transfected with psiCHECK2 plasmids (10 ng/well) by mixing the duplexes with 5 μL Opti-MEM containing Lipofectamine 2000 (2 μg/mL) and 5 μL psiCHECK2 plasmid, incubating the mixture for 15 min at rt, and then adding it to the cells. 48 h post-transfection, Firefly luciferase (transfection control) and Renilla luciferase (reporter, containing either on-target or off-target sequence) activities were measured according to the manufacturer's instructions. siRNA activity was determined by normalizing Renilla to Firefly signals within each well, and results were expressed relative to plasmid-only controls (% control). $IC_{50}$ values were calculated in Prism software using a four-parameter non-linear dose-response function. This assay was used to compare the relative effects of modifications within each experiment, rather than for direct quantitative comparison of $IC_{50}$ values across independent runs.

**Cell Culture Assays in A431 Cells.** Off-target effects of modified oligonucleotides targeting *ACTN1* mRNA were evaluated in human A431 cells (ATCC, CRL-1555). Cells were seeded at 10,000 cells per well in 96-well plates and transfected with *ACTN1* siRNAs across a 10-point dose-response range (5000, 1000, 200, 40, 8, 1.6, 0.32, 0.064, 0.0128, and 0.002 nM). Transfections were performed using Lipofectamine RNAiMAX (Thermo Fisher, #13778150) diluted in Opti-MEM (Gibco, #31985070), according to the manufacturer's reverse transfection protocol. After 96 h, cells were lysed and total RNA was isolated using a glass fiber filter plate (Pall, #5072). *ACTN1* mRNA levels were quantified by RT-qPCR on a QuantStudio 7 Flex Real-Time PCR System (Thermo Fisher). Reactions (5 µL) containing 1 µL RNA were set up with AgPath-ID One-Step RT-PCR reagents (Thermo Fisher) and custom primer-probe sets (Integrated DNA Technologies). Percent untreated control (% UTC) values were calculated using the formula:

% UTC = ((Sample QuantityTarget / Sample QuantityNormalization) / (Average UTC QuantityTarget / UTC QuantityNormalization)) × 100

Half-maximal inhibitory concentration ($IC_{50}$) values were determined in GraphPad Prism (v10; GraphPad Software, San Diego, CA) using the "log(inhibitor) vs. normalized response (variable slope)" function.

**Animal Studies.** All animal procedures were conducted in accordance with the guidelines of the American Association for the Accreditation of Laboratory Animal Care (AAALAC) and approved by the Animal Welfare Committee (Cold Spring Harbor Laboratory's IACUC). Mice were housed in micro-isolator cages under a 12 h light-dark cycle with controlled temperature and humidity, with food and water provided ad libitum.

Five-week-old C57BL/6J mice (Jackson Laboratories) received a single subcutaneous injection of *Ttr* siRNAs; four male C57BL/6 mice injected with PBS served as controls. For RNA analysis, 50-100 mg of liver tissue was homogenized in guanidinium thiocyanate with 8% β-mercaptoethanol using an Omni Tissue Homogenizer (Omni International). Total RNA was isolated with the PureLink Pro 96 Total RNA Purification Kit (Life Technologies, Carlsbad, CA). qRT-PCR was performed on a StepOne Real-Time PCR System (Life Technologies) using Express One-Step SuperMix qRT-PCR reagents (Life Technologies) and TaqMan primer/probe sets (Integrated DNA Technologies). The sequences for the mouse *Ttr* assay were: Forward: 5′-CGTACTGGAAGACACTTGGCATT-3′, Reverse: 5′-GAGTCGTTGGCTGTGAAAACC-3′, Probe: 5′-CCCGTTCCATGAATTCGCGGATG-3′. Target RNA levels were normalized to total RNA quantified with RIBOGREEN® RNA Quantitation Reagent (Molecular Probes). Results were expressed as percent *Ttr* RNA relative to PBS-treated controls (% control). Half-maximal effective dose ($ED_{50}$) values were calculated in GraphPad Prism 10 (GraphPad Software, San Diego, CA). For tolerability evaluation, single subcutaneous injections of Marc1 siRNAs were administered to 7-8 week-old BALB/c mice (Charles River Laboratories). Mice were euthanized 7 days post-injection for histopathology assessment, and cardiac blood was collected for clinical chemistry. The *Marc1* primer/probe sequences were: Forward: 5′-GAAACGGGTGATGGCTTGTA-3′, Reverse: 5′-GCGGTAGCTCTTCAGTGTTT-3′, Probe: 5′-CTTCCTGTCCGAGATGCCAGTGTC-3′.

**RNA Sample Processing and DGE Analysis.** RNA samples from *in vitro* and *in vivo* studies were subjected to differential gene expression (DGE) profiling using the QuantSeq 3′ mRNA-Seq

Library Prep Kit FWD for Illumina on the Illumina sequencing platform. Sequencing yielded more than 2.5 million unique mapped reads per sample, providing expression data for over 10,000 unigenes. Differentially expressed genes were defined as those showing >2-fold change (up- or down-regulation) with $p < 0.01$ and $q < 0.1$. Data were visualized using volcano plots, where the x-axis represents $\log_2$ fold change and the y-axis represents –log10 p-value.

To assess siRNA seed-match-mediated off-target effects, cumulative distribution function (CDF) curves of $\log_2$ fold change were generated for genes with or without seed matches. A leftward shift of seed-match genes relative to the no-seed baseline indicated a greater fraction of seed-match genes being downregulated. Estimates of $\Delta$ $\log_2$ fold change for seed-match categories were obtained from beta coefficients of a linear model regressing $\log_2$ fold change against seed-match type, with statistical significance evaluated using two-tailed t-statistics. Seed-match genes were identified using TargetRank (http://hollywood.mit.edu/targetrank)

**Histology.** Liver samples were fixed in 10% neutral buffered formalin and processed using a Sakura Tissue Tek tissue processor. Four-micron sections were cut, air-dried overnight, and baked at 60 °C for 1 h. Sections were stained on a Leica Spectra stainer with Gills II Hematoxylin (Leica/Surgipath, cat# 3801520; 4 min), differentiated in 0.5% acid alcohol (1 s), and counterstained with Eosin (Leica/Surgipath, cat# 3801600; 2 min). Stained slides were scanned at 20× resolution using a Hamamatsu S360 scanner and evaluated by a board-certified pathologist.

**Molecular Dynamics Simulations.** The modified residues of the *Ttr* sequence studied in this work (vinyl phosphonate T, 2′-fluoro nucleosides, 2′-O-methyl nucleosides, and the aLd and dx analogs) were built using the modXNA methodology as described by Love and collaborators.[20] A starting RNA•RNA duplex was created with BIOVIA Discovery Studio v24.2.500 and used as a template.

Modified nucleotides were inserted to reproduce the *Ttr* sequence (Table 1). The resulting duplex was structurally relaxed and then inserted into the hAgo2 system based on PDB entry 6N4O.[21] Missing regions of the protein were modeled using Discovery Studio. AMBER-type topology and coordinate files (prmtop/inpcrd) were generated with LEaP (AmberTools 23 and 24). Canonical nucleic acids were described using the OL21 force field[22, 23], and protein was parameterized with FF14SB.[24] The system was solvated in a truncated octahedral periodic box with OPC water molecules (12 Å solute-edge buffer). NaCl ions were added to neutralize the system, using the Joung-Cheatham parameters.[25] Initial minimization and heating were performed at 310 K using the AmberMDPrep protocol[26] with default parameters. Each system was simulated in triplicate, with independent random seeds, for approximately 5 μs per replica using the PMEMD CUDA engine in AMBER 20. Trajectories were analyzed with CPPTRAJ v6.29.8 (AmberTools).[27]

**Table S1.** Mass and UV purity characterization of synthesized siRNAs

| Target | As/ Sense | Modification | Formula | Average Mass | Observed Mass | UV purity (%) |
|---|---|---|---|---|---|---|
| *Ttr* | Sense | N/A | $C_{280}H_{390}N_{81}O_{171}P_{21}S_2F_4$ | 8417.0 | 8416.0 | 96.0 |
| *Ttr* | As | None | $C_{242}H_{310}N_{85}O_{154}P_{23}S_4F_4$ | 7790.1 | 7789.2 | 95.0 |
| *Ttr* | As | dx G6 | $C_{242}H_{311}N_{85}O_{154}P_{23}S_4F_3$ | 7772.2 | 7771.2 | 92.1 |
| *Ttr* | As | dx A7 | $C_{241}H_{308}N_{85}O_{153}P_{23}S_4F_4$ | 7760.1 | 7759.0 | 90.6 |
| *Ttr* | As | aLd G6 | $C_{242}H_{311}N_{85}O_{154}P_{23}S_4F_3$ | 7772.2 | 7771.6 | 95.7 |
| *Ttr* | As | aLd A7 | $C_{241}H_{308}N_{85}O_{153}P_{23}S_4F_4$ | 7760.1 | 7759.5 | 91.5 |
| *ACTN1* | Sense | N/A | $C_{279}H_{389}N_{78}O_{173}P_{21}S_4F_4$ | 8458.1 | 8457.1 | 96.5 |
| *ACTN1* | As | None | $C_{244}H_{314}N_{93}O_{148}P_{23}S_4F_4$ | 7834.2 | 7833.6 | 95.4 |
| *ACTN1* | As | dx G6 | $C_{244}H_{315}N_{93}O_{148}P_{23}S_4F_3$ | 7816.3 | 7815.3 | 95.9 |
| *ACTN1* | As | dx A7 | $C_{243}H_{312}N_{93}O_{147}P_{23}S_4F_4$ | 7804.3 | 7803.2 | 90.7 |
| *ACTN1* | As | aLd G6 | $C_{244}H_{315}N_{93}O_{148}P_{23}S_4F_3$ | 7816.3 | 7815.6 | 93.3 |
| *ACTN1* | As | aLd A7 | $C_{243}H_{312}N_{93}O_{147}P_{23}S_4F_4$ | 7804.3 | 7803.6 | 94.6 |
| *Marc1* | Sense | N/A | $C_{281}H_{392}N_{85}O_{170}P_{21}S_2F_4$ | 8471.1 | 8470.1 | 96.1 |
| *Marc1* | As | None | $C_{243}H_{313}N_{90}O_{151}P_{23}S_4F_4$ | 7827.2 | 7826.2 | 94.6 |
| *Marc1* | As | dx G6 | $C_{243}H_{314}N_{90}O_{151}P_{23}S_4F_3$ | 7809.3 | 7808.5 | 95.9 |
| *Marc1* | As | dx G7 | $C_{242}H_{311}N_{90}O_{150}P_{23}S_4F_4$ | 7797.2 | 7796.4 | 92.8 |
| *Marc1* | As | aLd G6 | $C_{243}H_{314}N_{90}O_{151}P_{23}S_4F_3$ | 7809.3 | 7808.4 | 95.7 |
| *Marc1* | As | aLd G7 | $C_{242}H_{311}N_{90}O_{150}P_{23}S_4F_4$ | 7797.2 | 7796.5 | 95.1 |

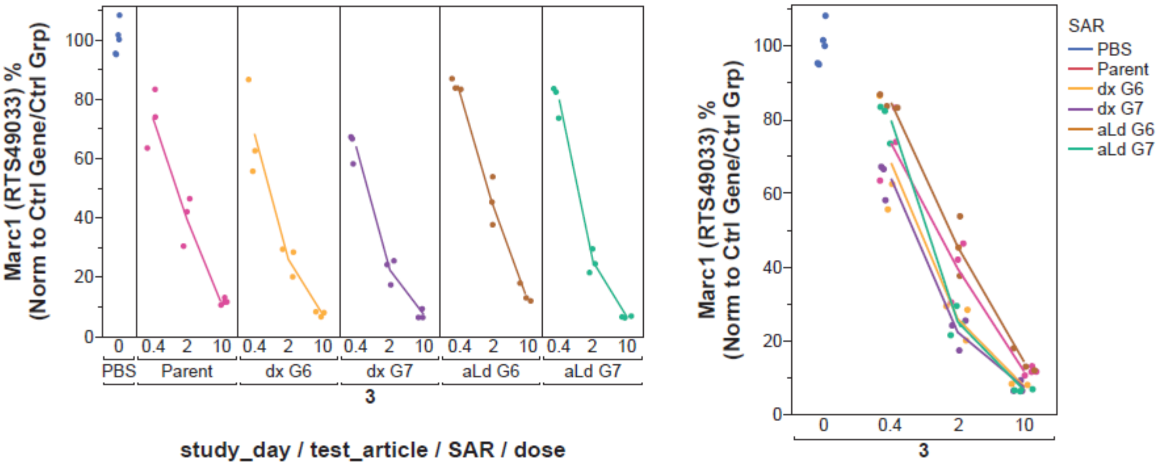


| siRNA | Parent | dx G6 | dx G7 | aLd G6 | aLd G7 |
|---|---|---|---|---|---|
| On-target | 0.985 | 0.590 | 0.480 | 1.50 | 0.86 |

**Figure S1.** Hepatic *Marc1* mRNA knockdown after three doses (0.4, 2, and 10 mg/kg). qPCR analysis showed that all modified siRNAs maintained on-target suppression comparable to the parent siRNA, but the dx at position 7 modification enhanced activity, lowering the $ED_{50}$ from 0.985 mg/kg (parent) to 0.480 mg/kg. All $ED_{50}$ values are reported in mg/Kg

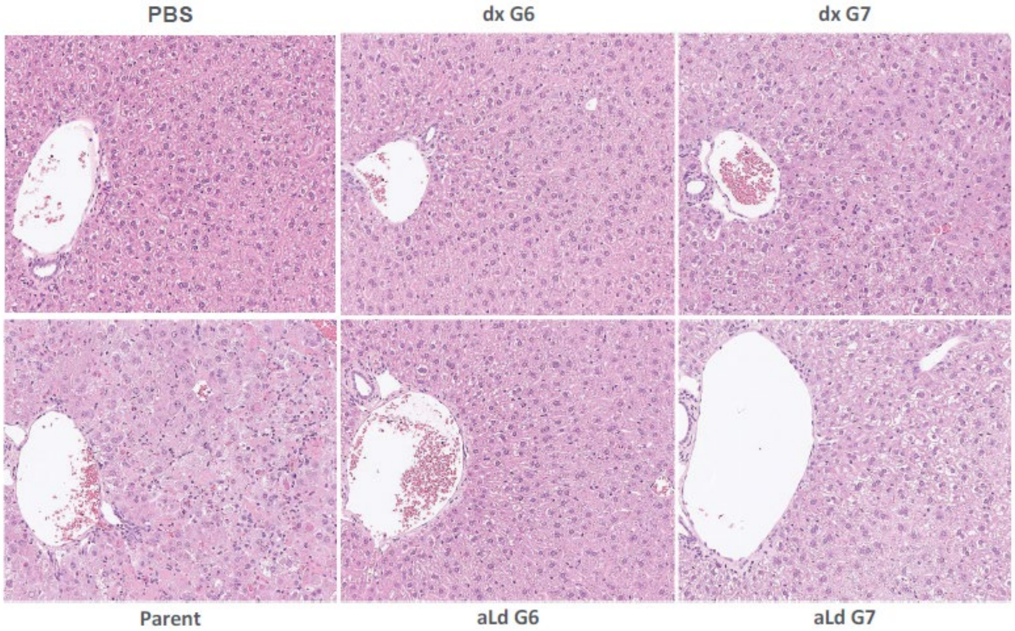


**Figure S2.** Liver histopathology one week after a 10 mg/kg dose of parent and modified *Marc1* siRNAs. H&E-stained liver sections collected one week after dosing show marked injury with the parent siRNA, whereas dx G6 and aLd G6 appear normal, and dx G7 and aLd G7 exhibit minimal to mild alterations.

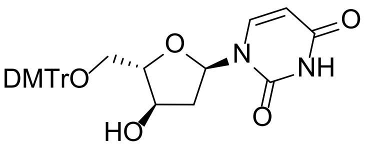

**8**

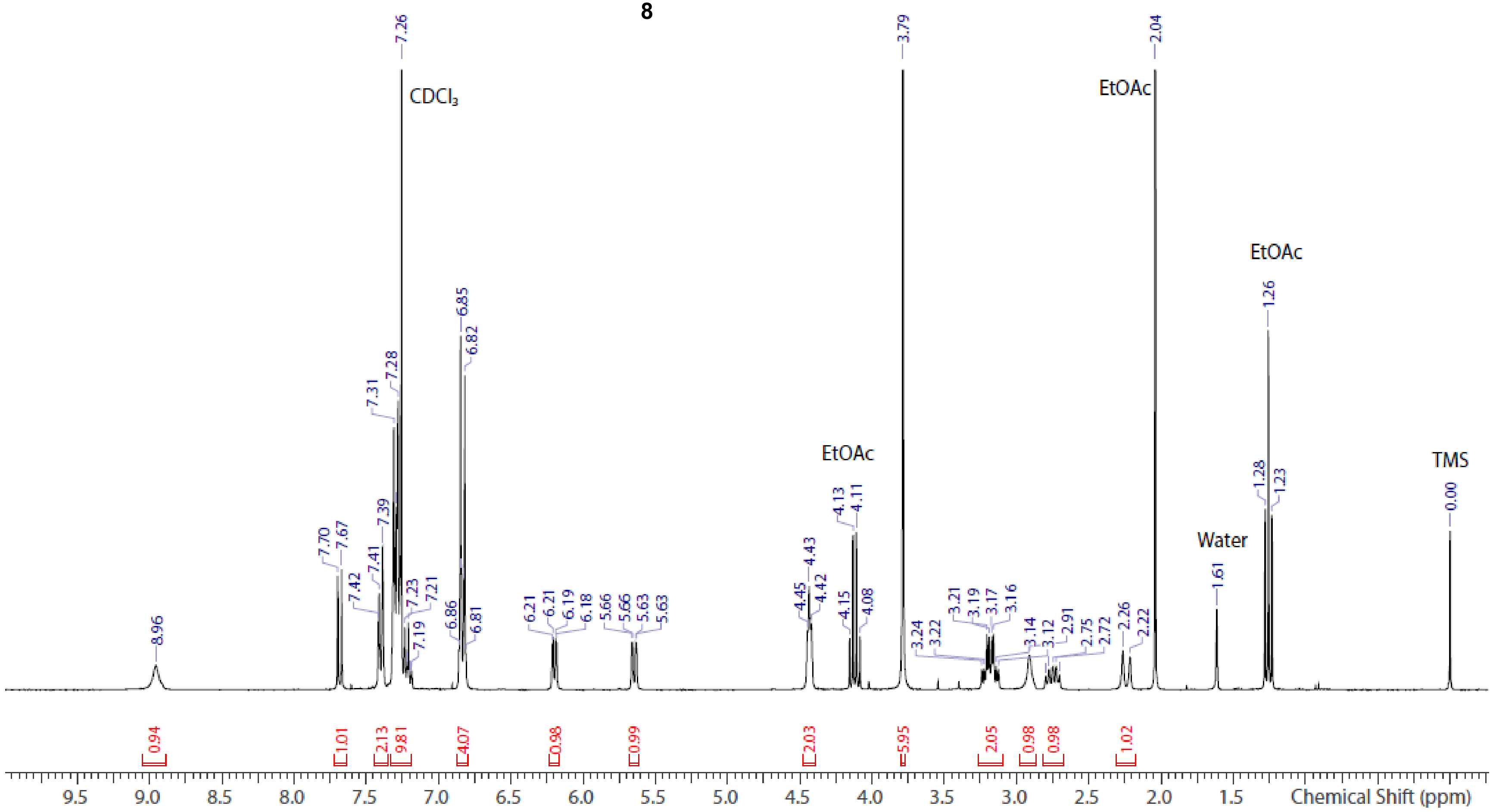


**Spectrum S1.** [1]H NMR spectrum of aLd U **8** ($CDCl_3$)

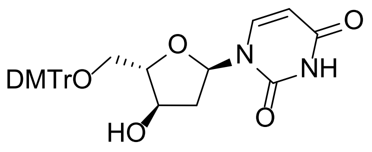


**8**

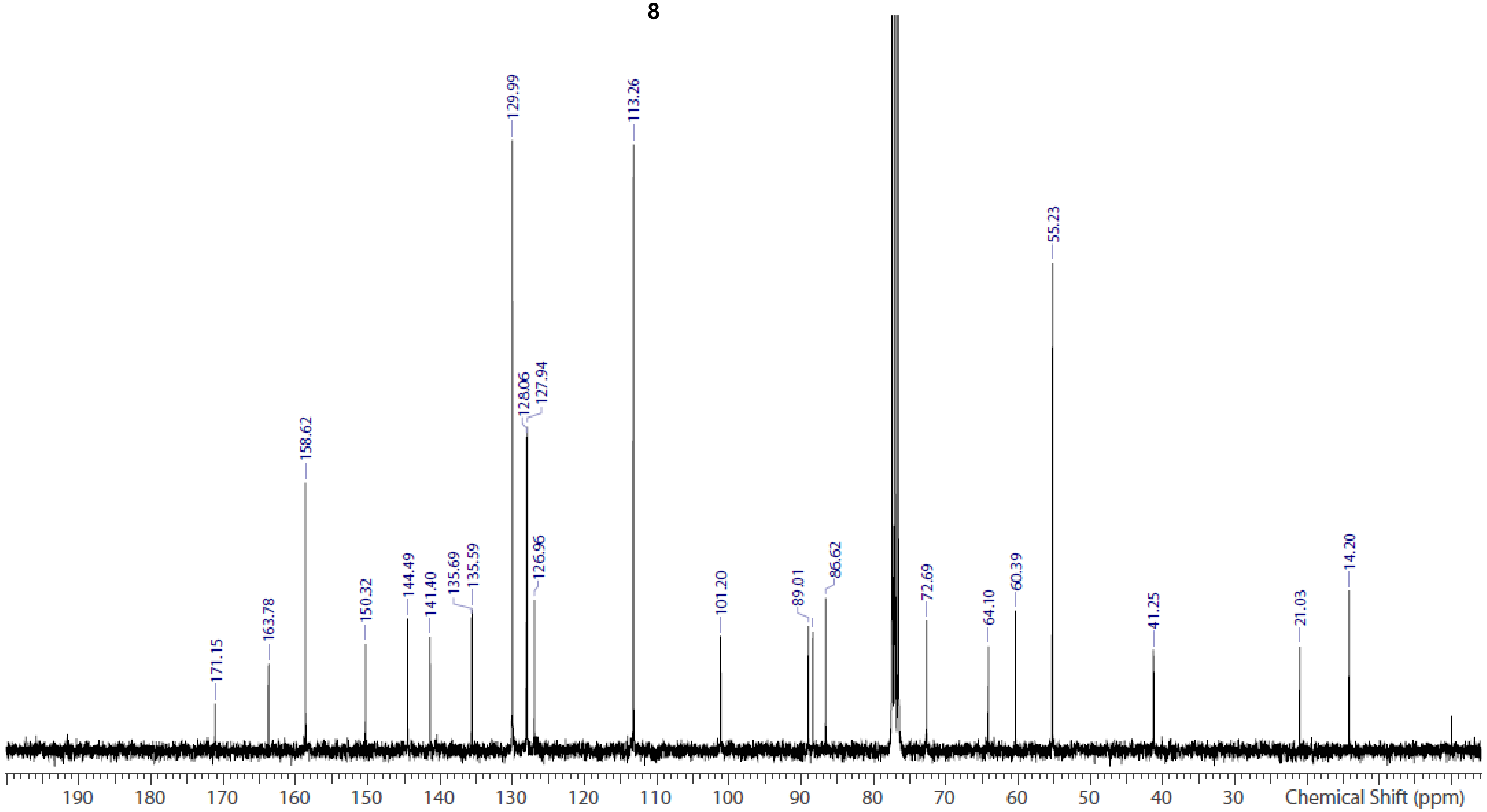


**Spectrum S2.** $^{13}C$ NMR spectrum of aLd U **8** ($CDCl_3$)

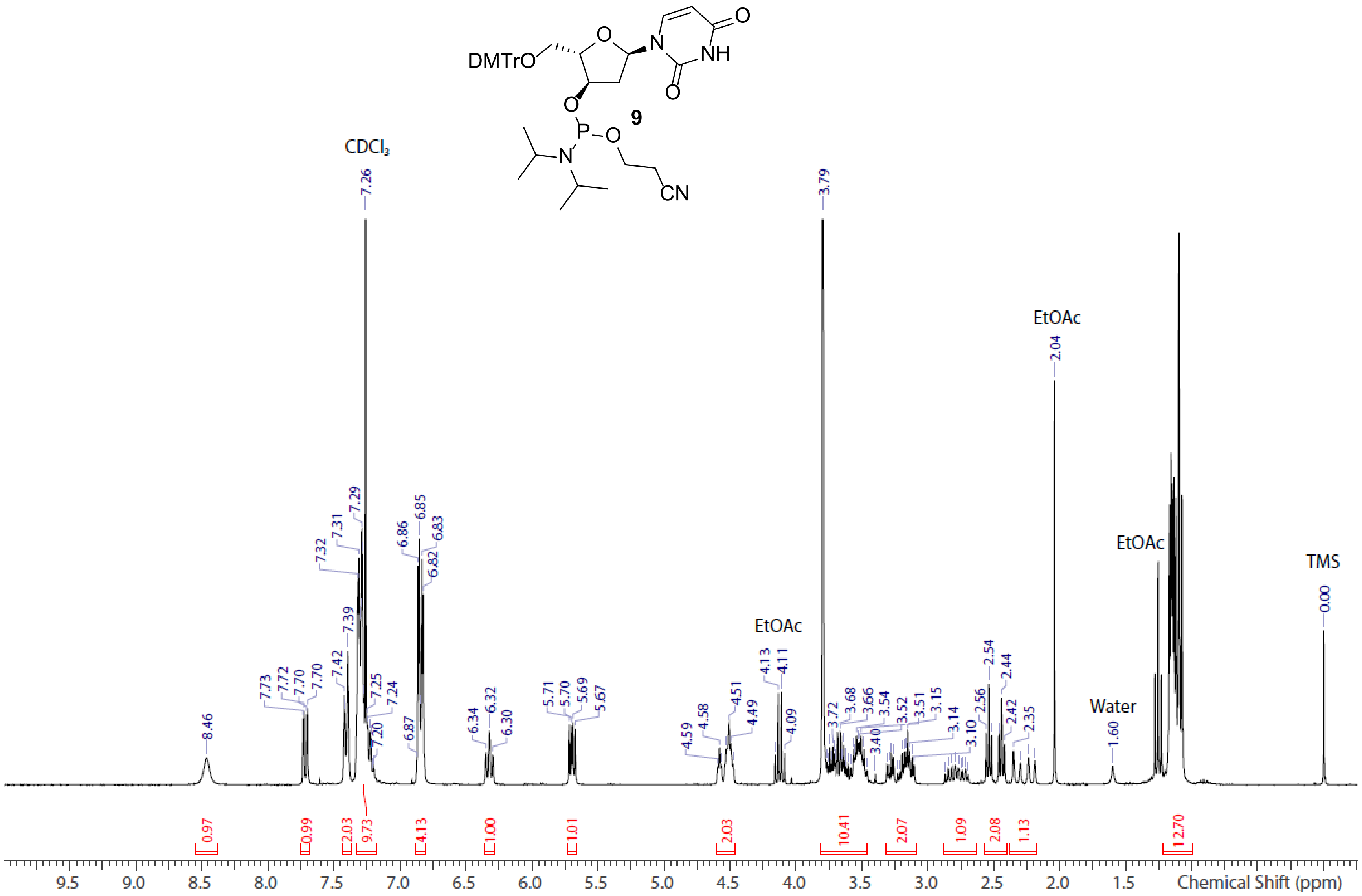


**Spectrum S3.** [1]H NMR spectrum of aLd U phosphoramidite **9** ($CDCl_3$)

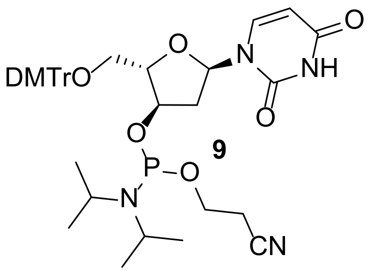


**9**

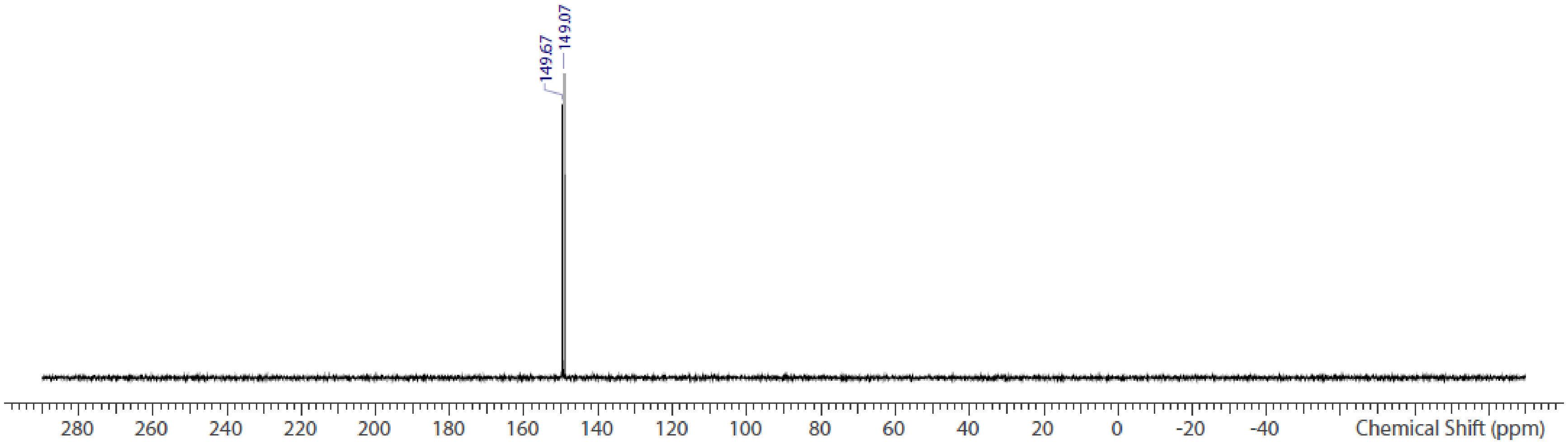


**Spectrum S4.** $^{31}P$ NMR spectrum of aLd U phosphoramidite **9** ($CDCl_3$)

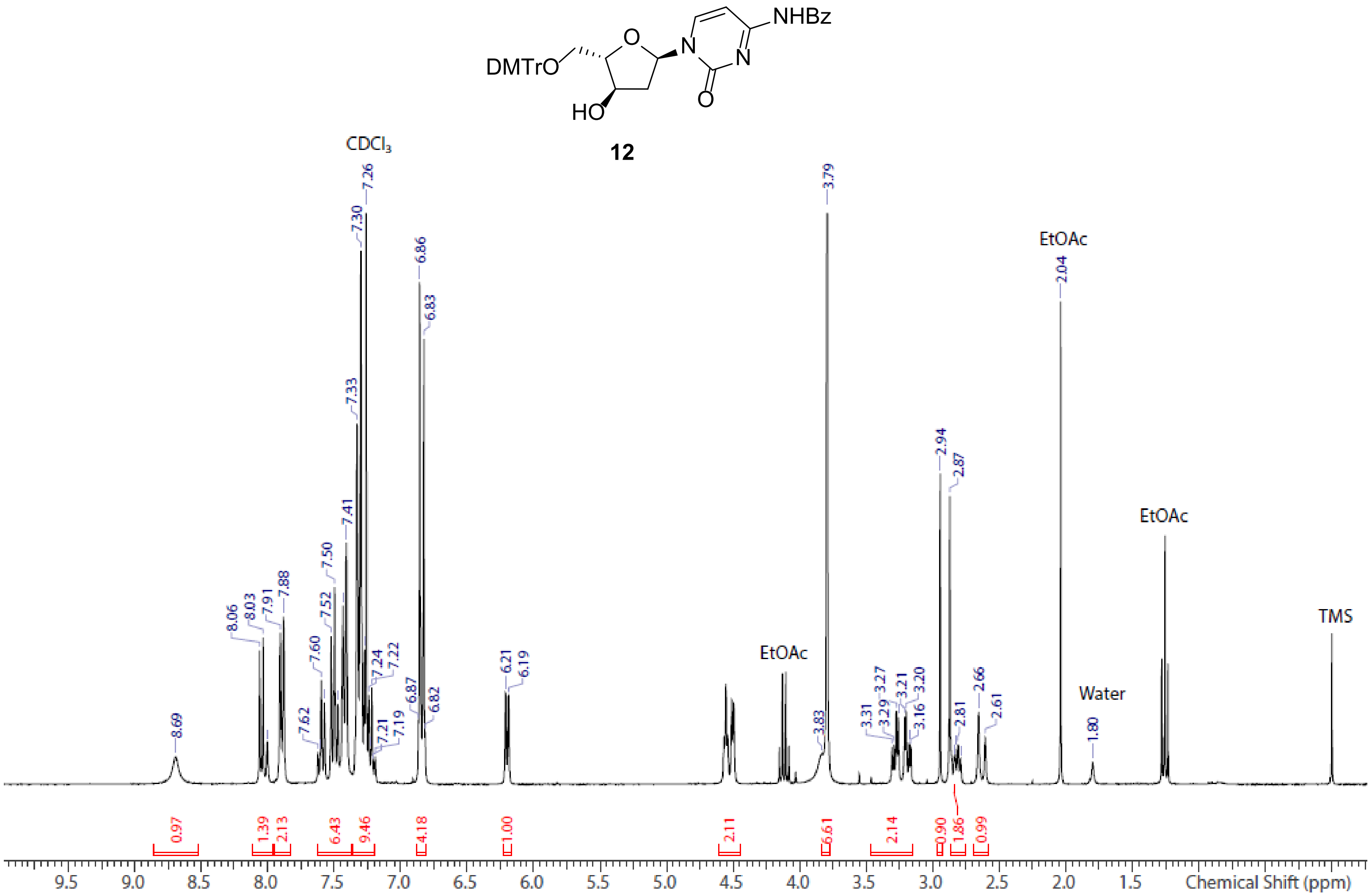


**Spectrum S5.** $^1$H NMR spectrum of aLd C **12** ($CDCl_3$)

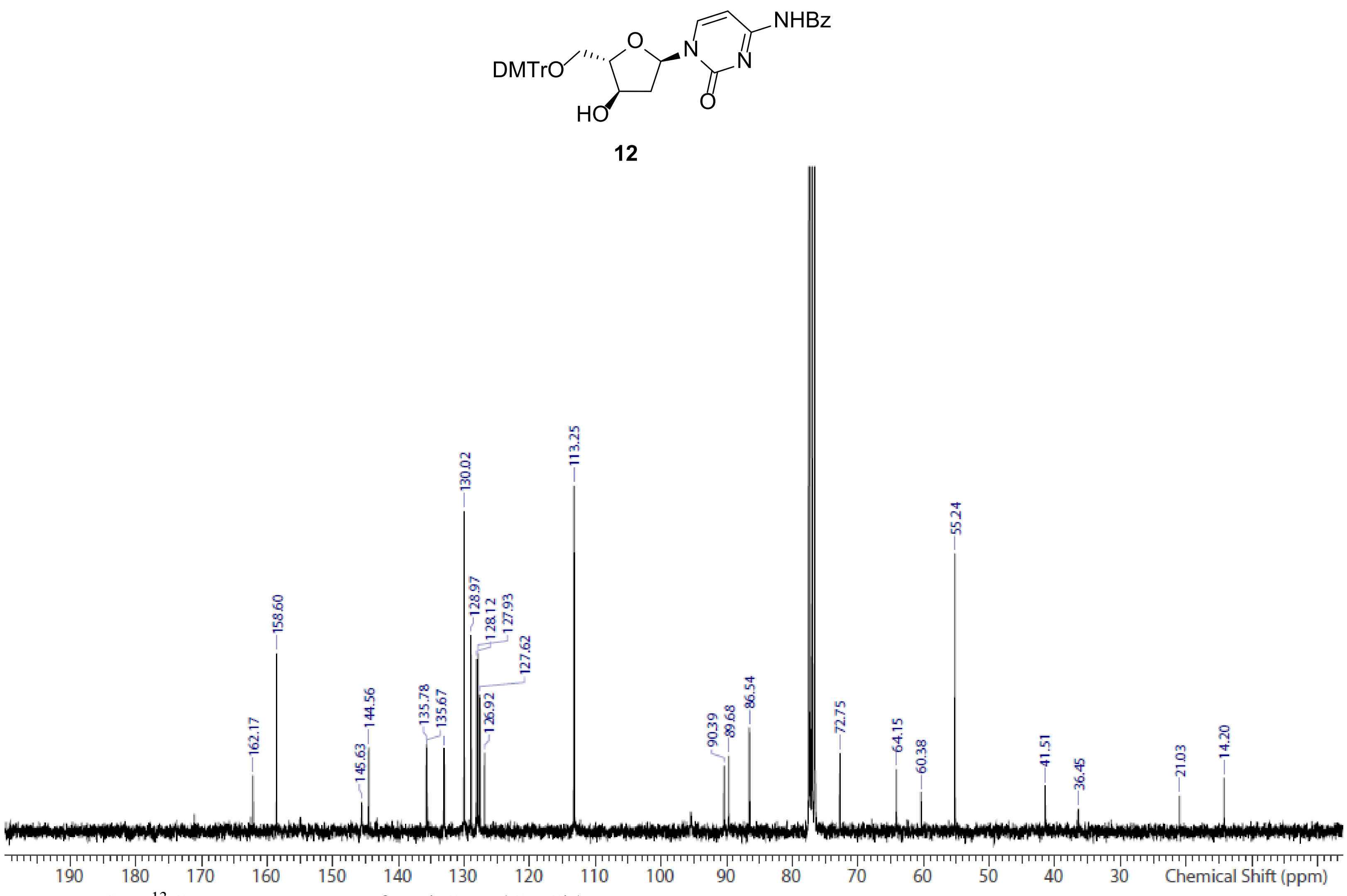


**Spectrum S6.** $^{13}C$ NMR spectrum of aLd C **12** ($CDCl_3$)

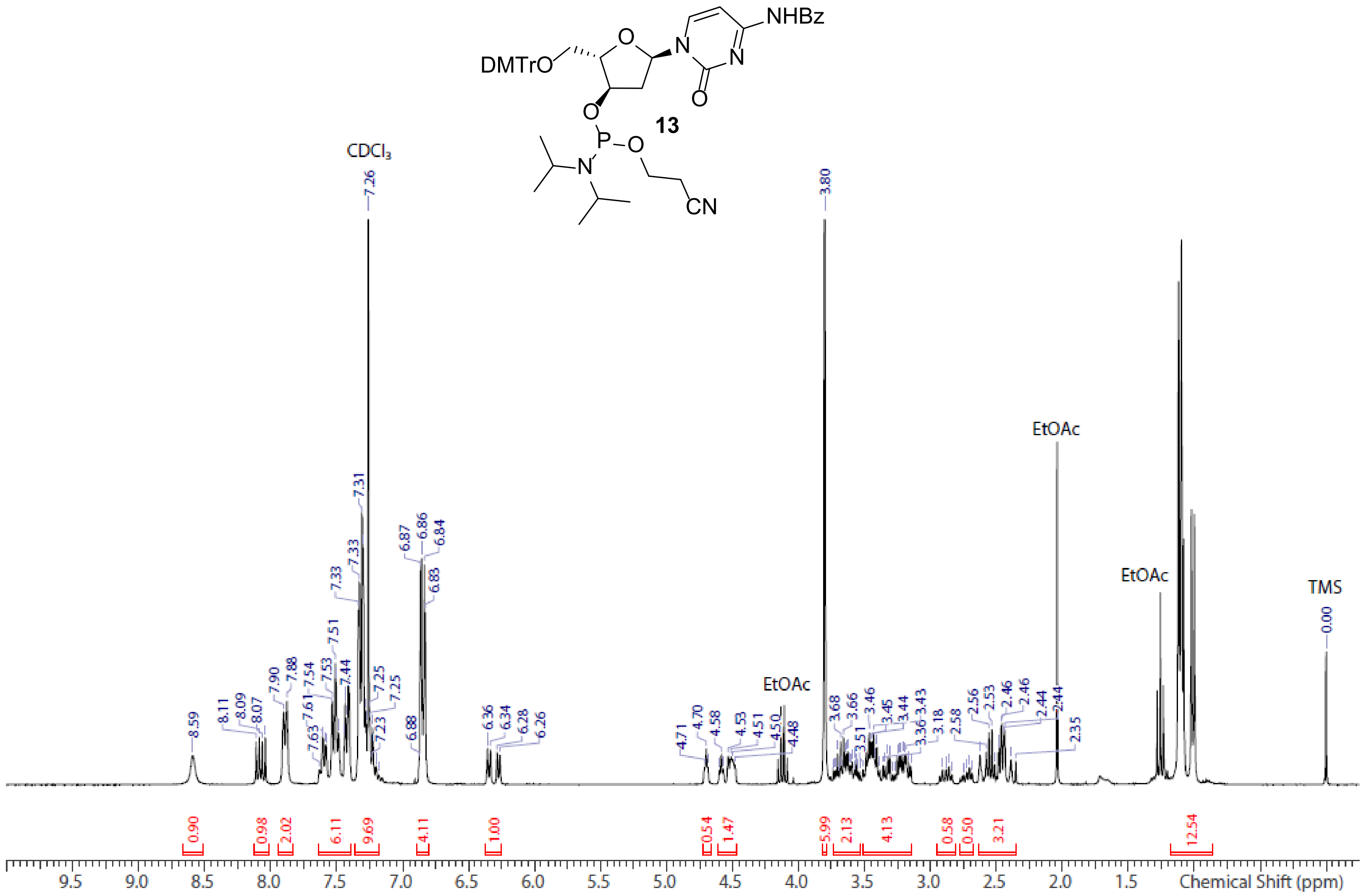


**Spectrum S7.** $^{1}$H NMR spectrum of aLd C phosphoramidite **13** ($CDCl_3$)

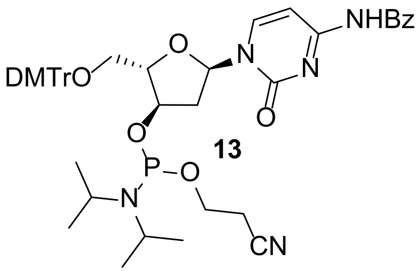


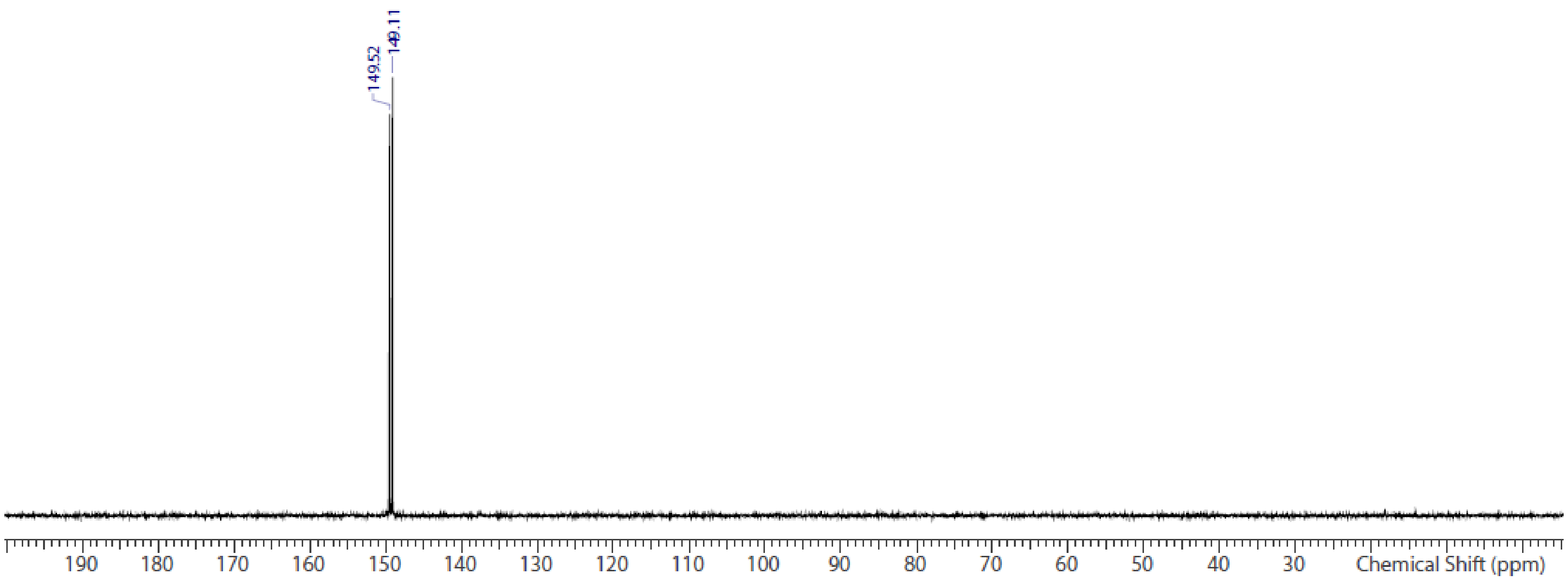


**Spectrum S8.** $^{31}P$ NMR spectrum of aLd C phosphoramidite **13** ($CDCl_3$)

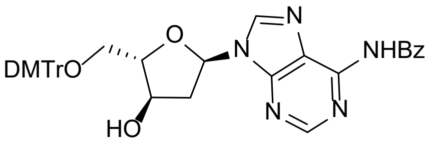


**21**

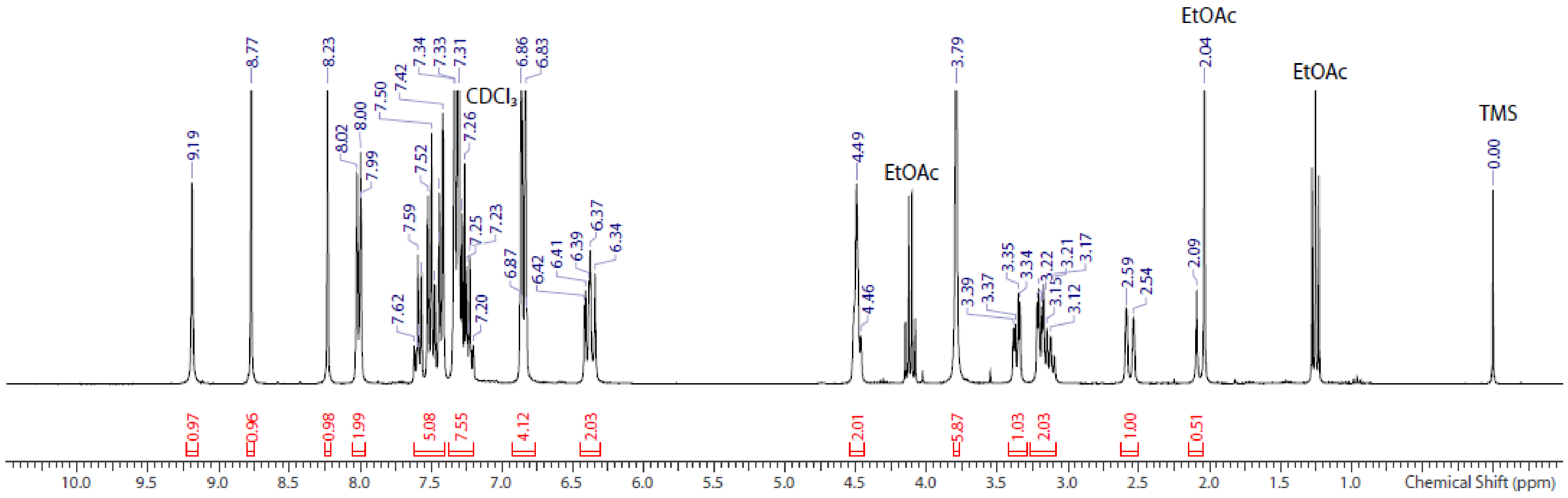


**Spectrum S9.** [1]H NMR spectrum of aLd A **21** ($CDCl_3$)

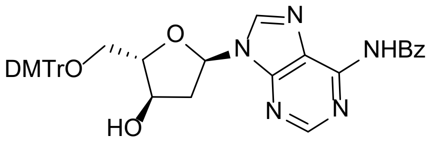


**21**

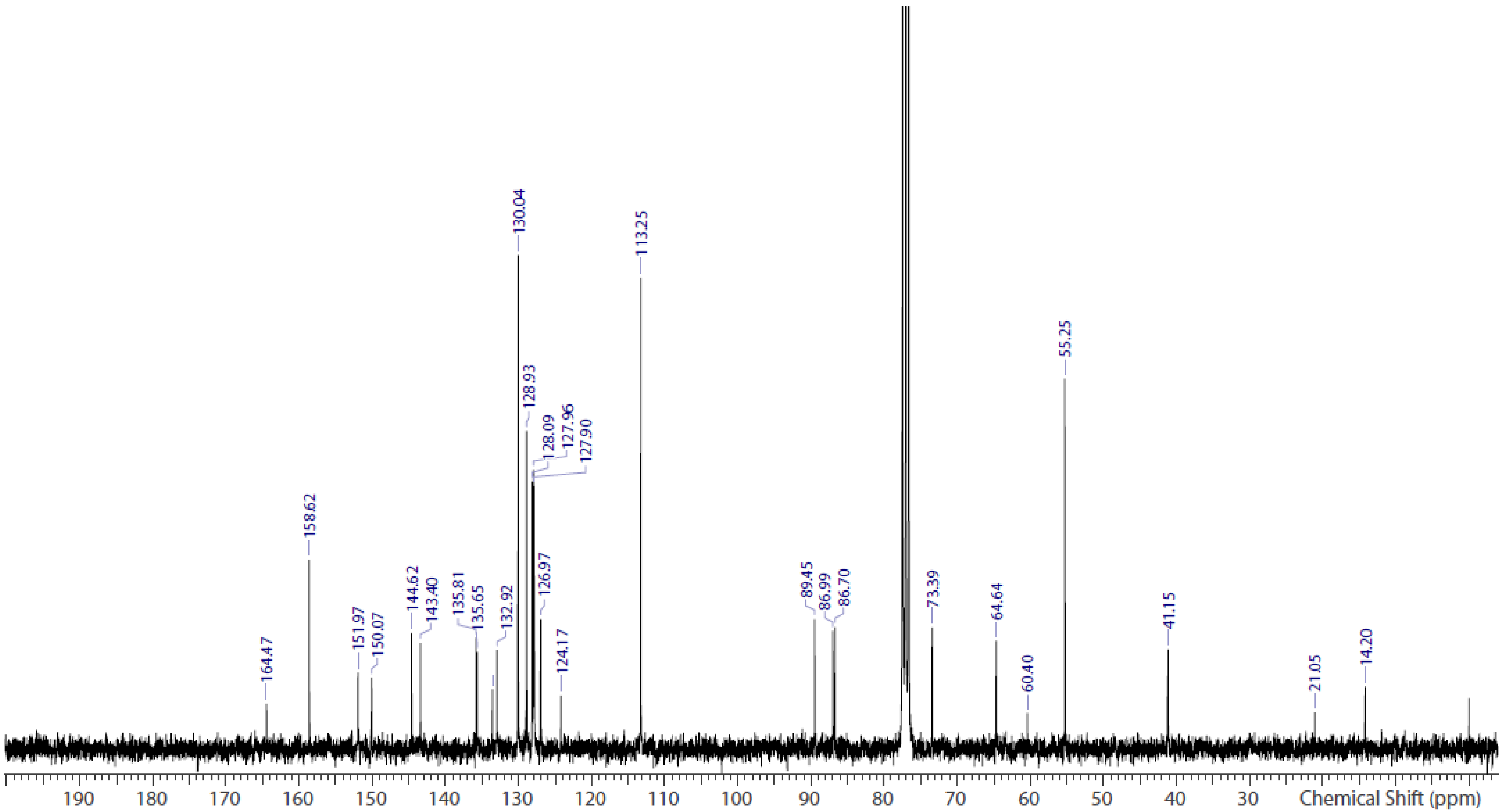


**Spectrum S10.** $^{13}C$ NMR spectrum of aLd A **21** ($CDCl_3$)

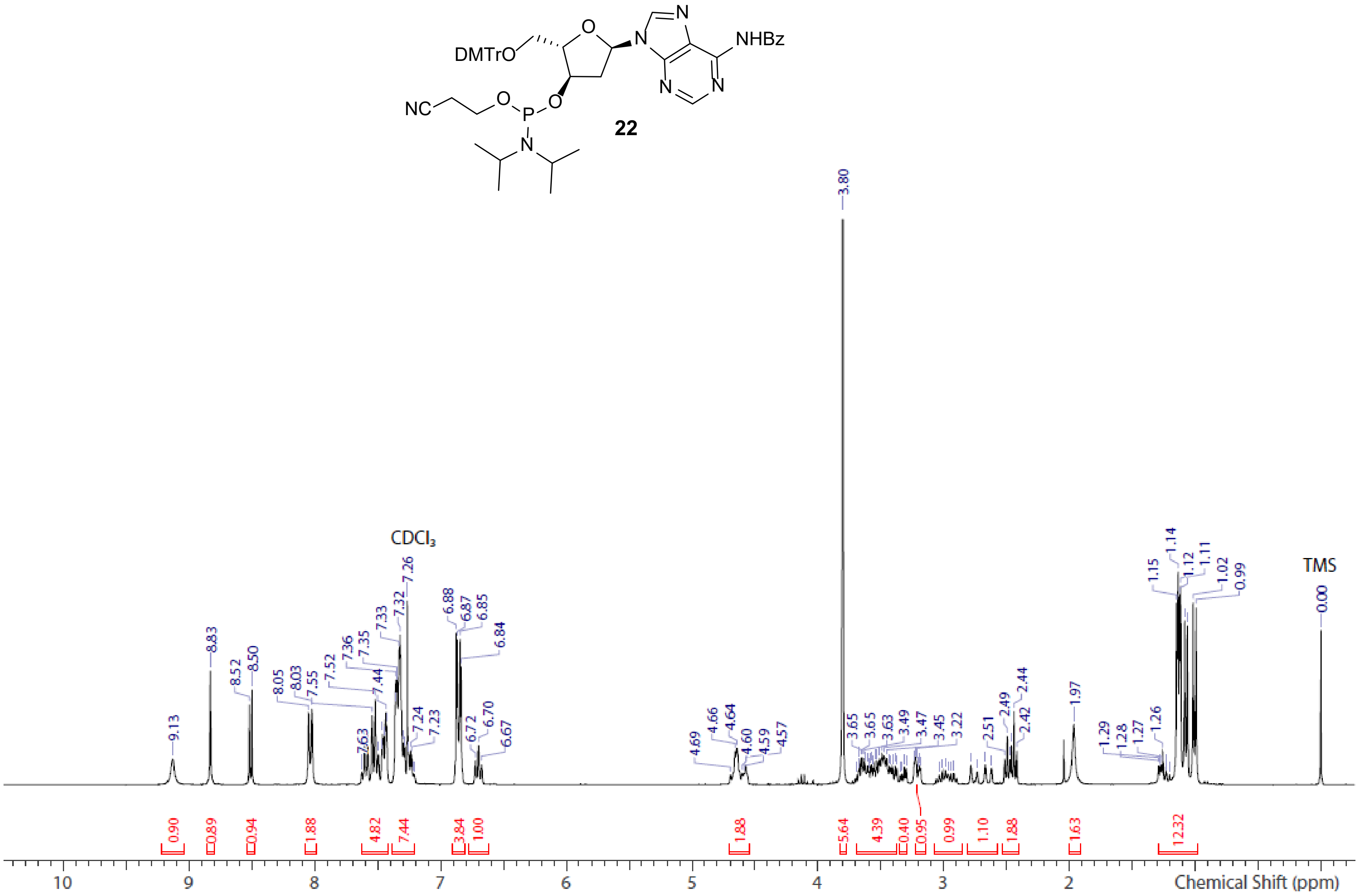


**Spectrum S11.** $^1$H NMR spectrum of aLd A phosphoramidite **22** ($CDCl_3$)

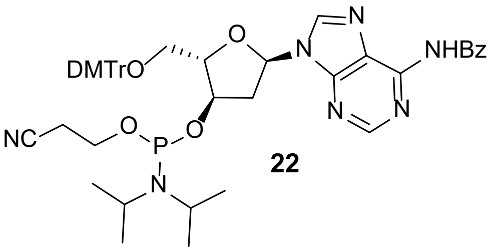


**22**

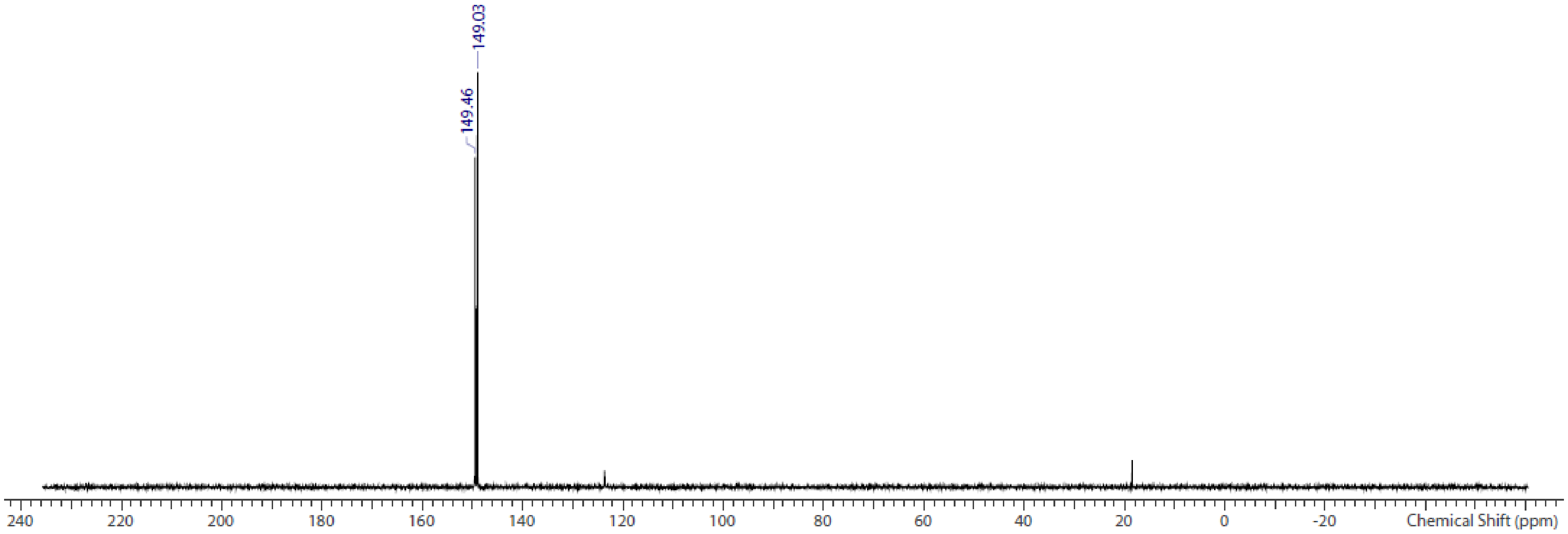


**Spectrum S12.** $^{31}$P NMR spectrum of aLd A phosphoramidite **22** ($CDCl_3$)

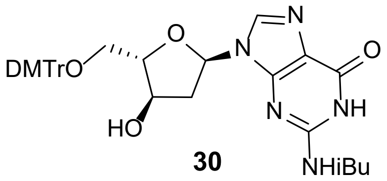


**30**

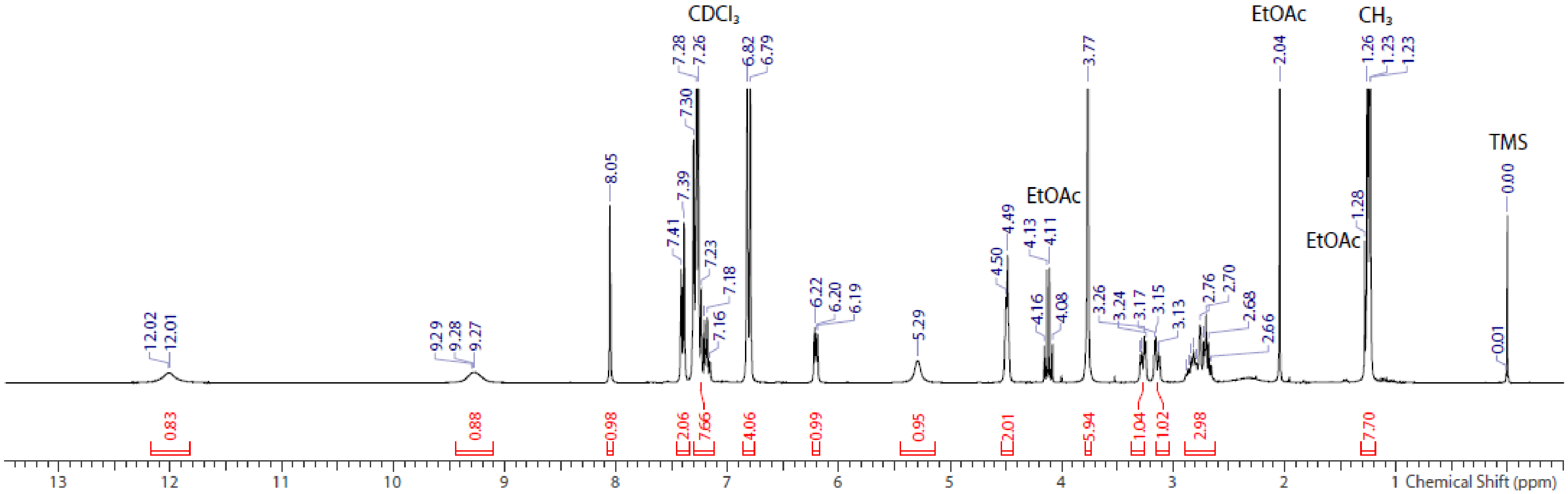


**Spectrum S13.** [1]H NMR spectrum of aLd G **30** ($CDCl_3$)

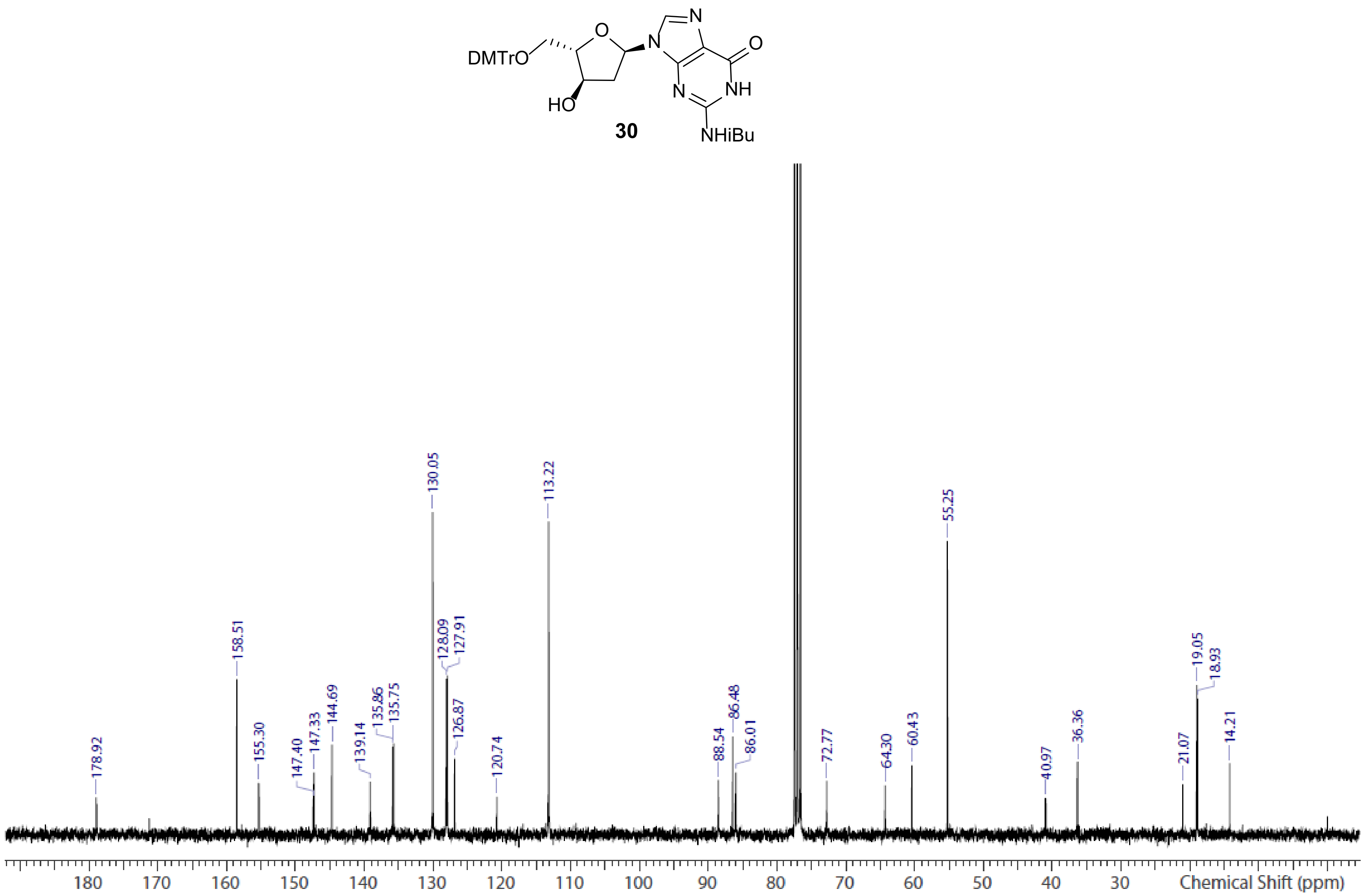


**Spectrum S14.** $^{13}$C NMR spectrum of aLd G **30** ($CDCl_3$)

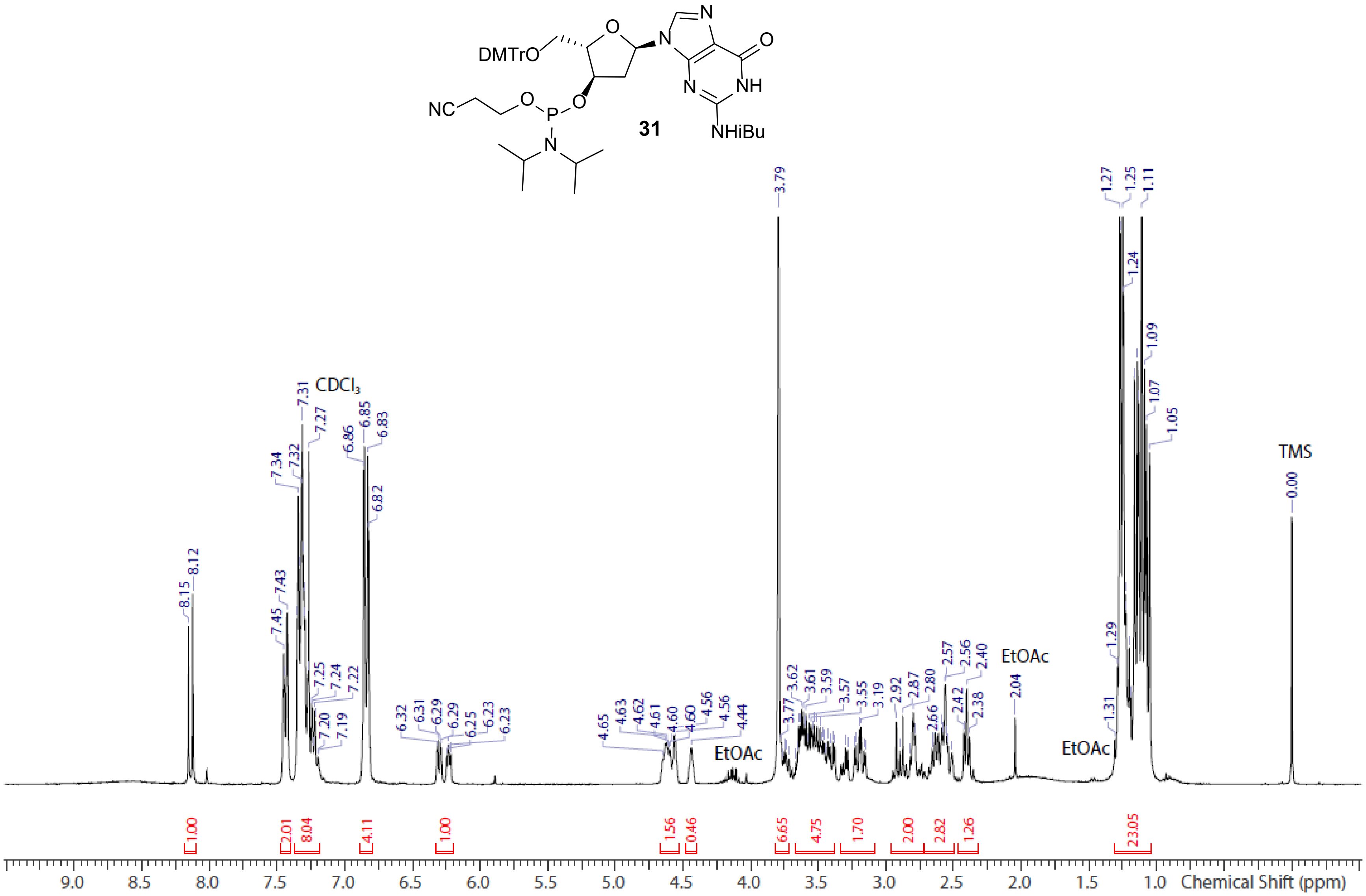


**Spectrum S15.** [1]H NMR spectrum of aLd G phosphoramidite **31** ($CDCl_3$)

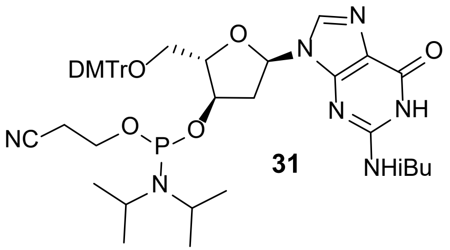


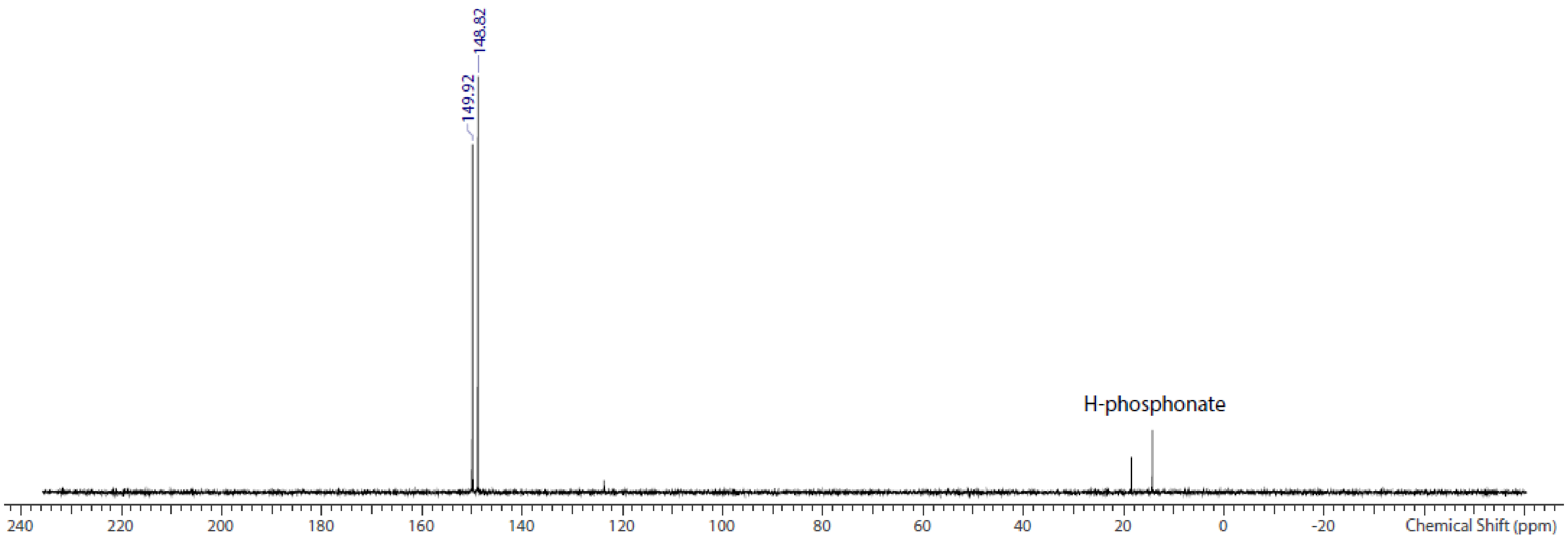


**Spectrum S16.** $^{31}$P NMR spectrum of aLd G phosphoramidite **31** ($CDCl_3$)

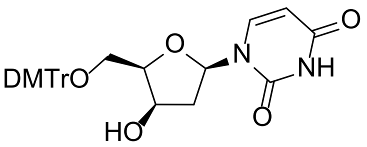


**34**

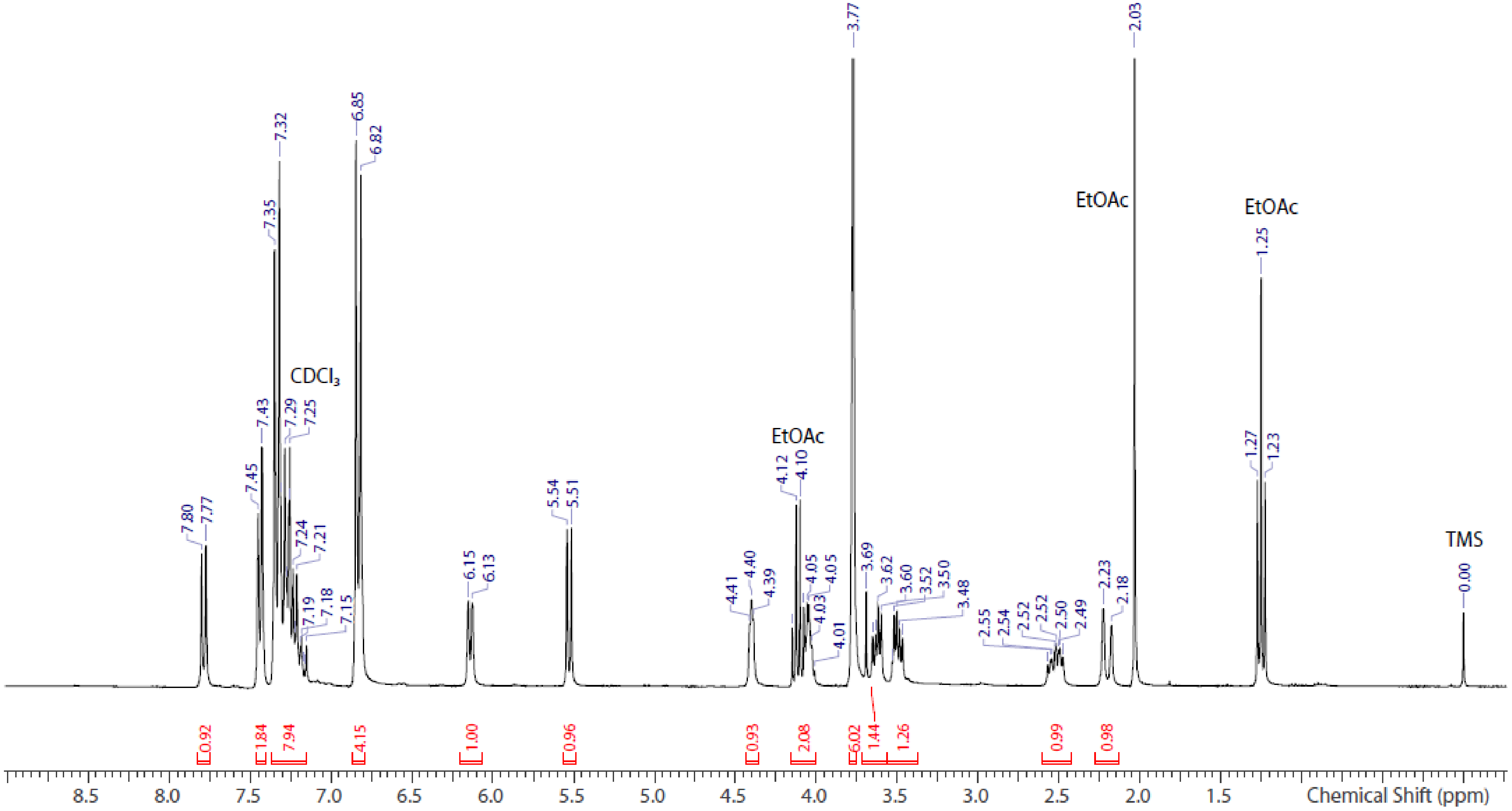


**Spectrum S17.** $^{1}$H NMR spectrum of dx U **34** ($CDCl_3$)

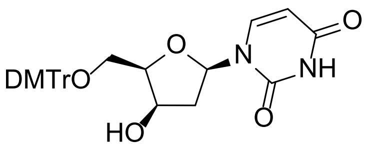


**34**

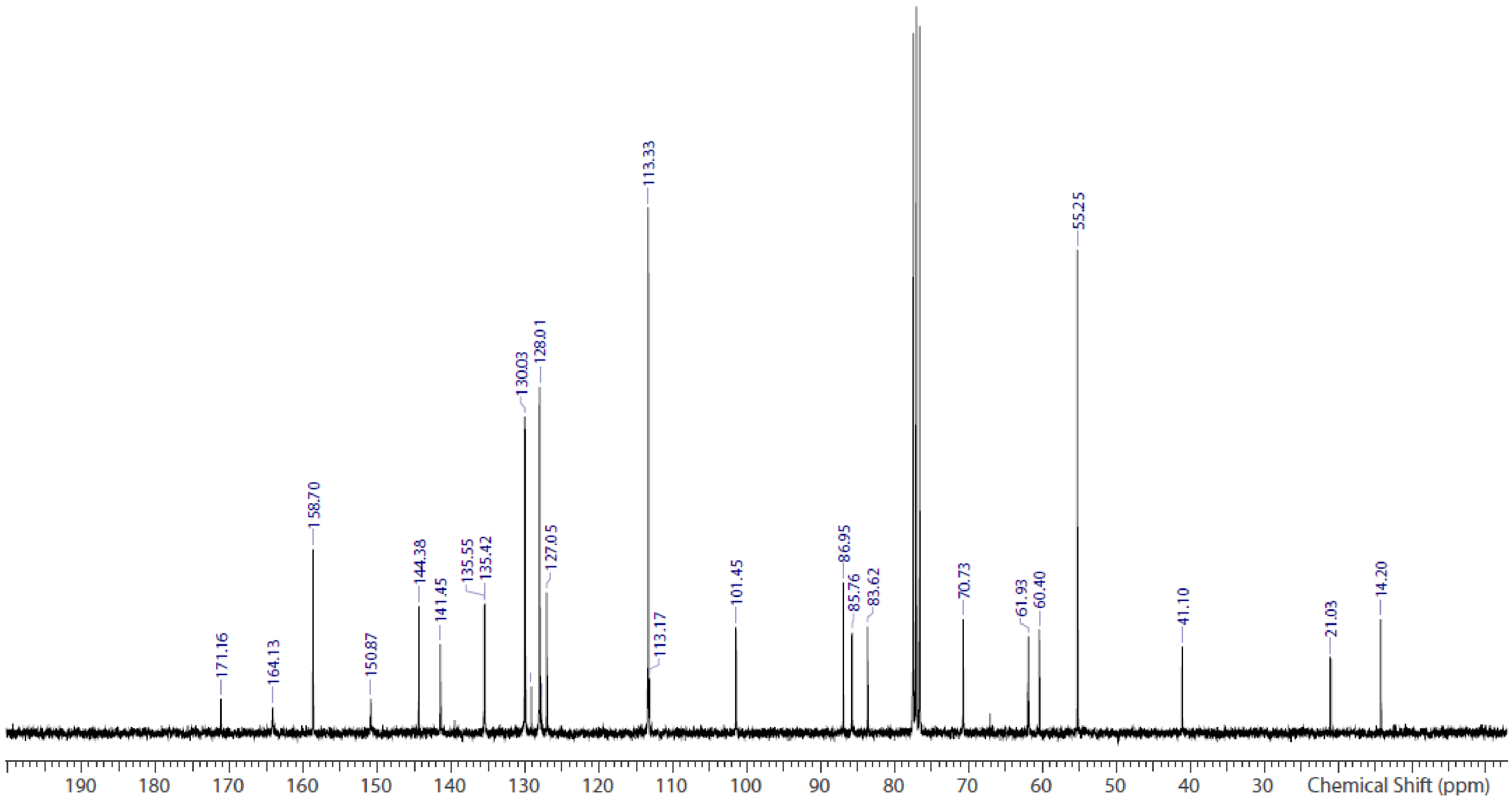


**Spectrum S18.** $^{13}$C NMR spectrum of dx U **34** ($CDCl_3$)

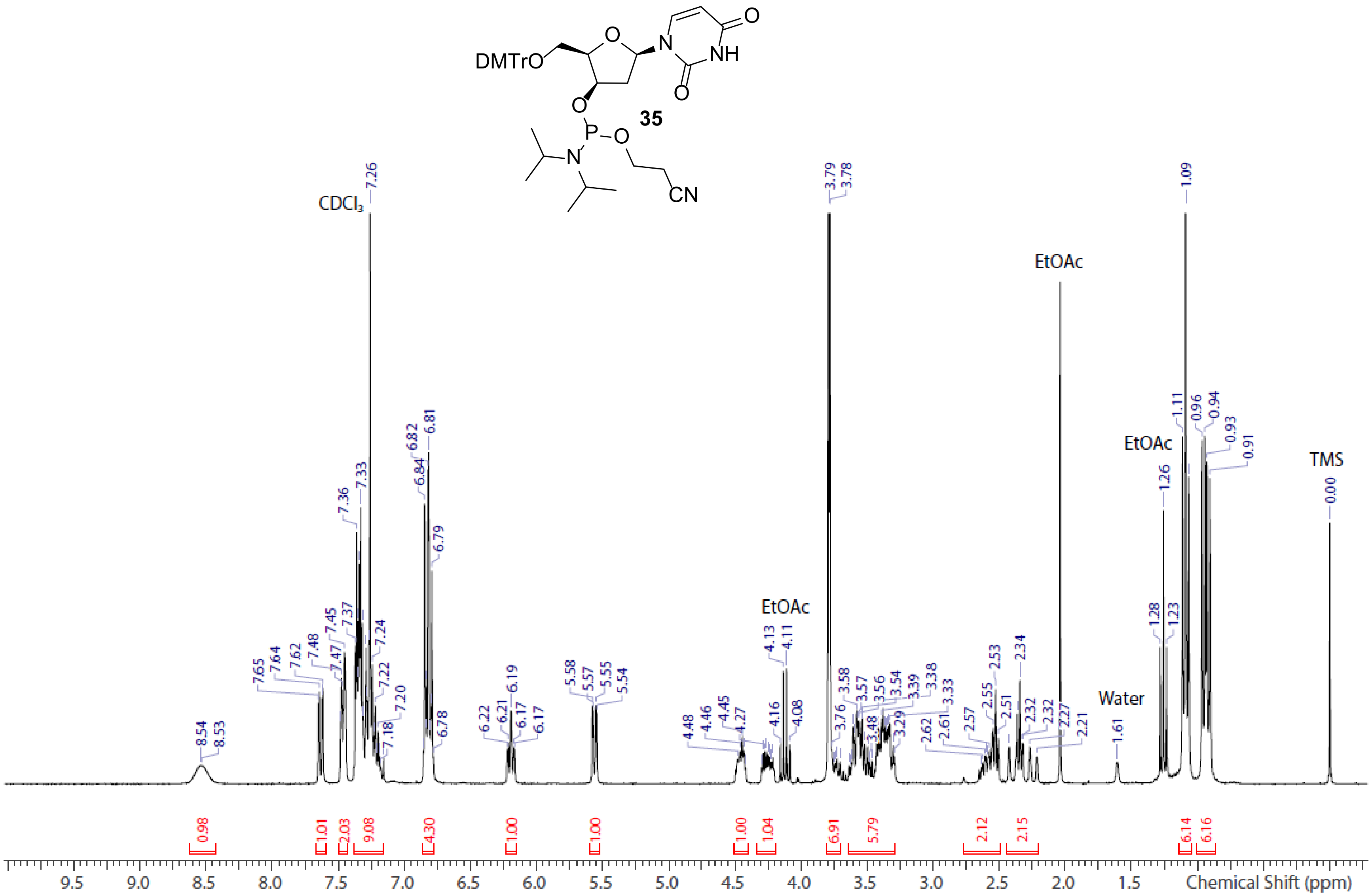


**Spectrum S19.** $^{1}$H NMR spectrum of dx U phosphoramidite **35** ($CDCl_3$)

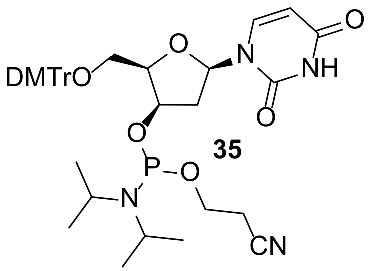


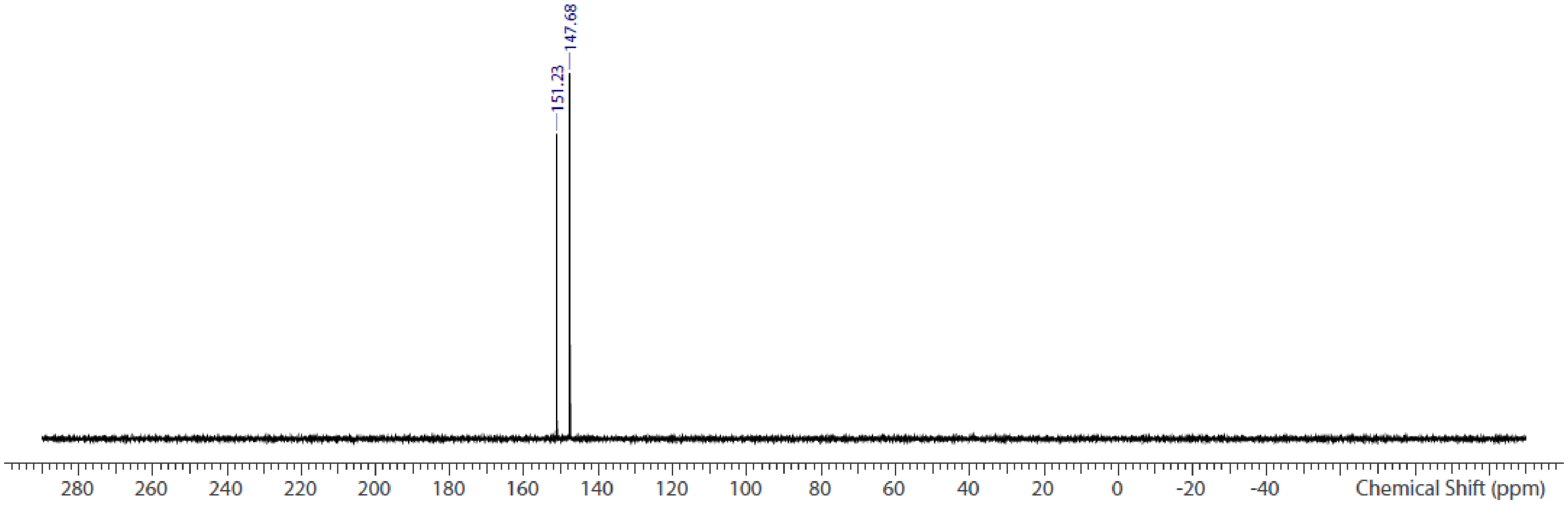


**Spectrum S20.** $^{31}$P NMR spectrum of dx U phosphoramidite **35** ($CDCl_3$)

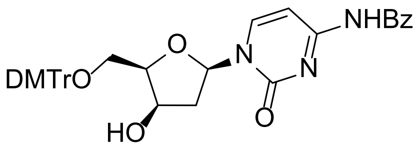


**40**

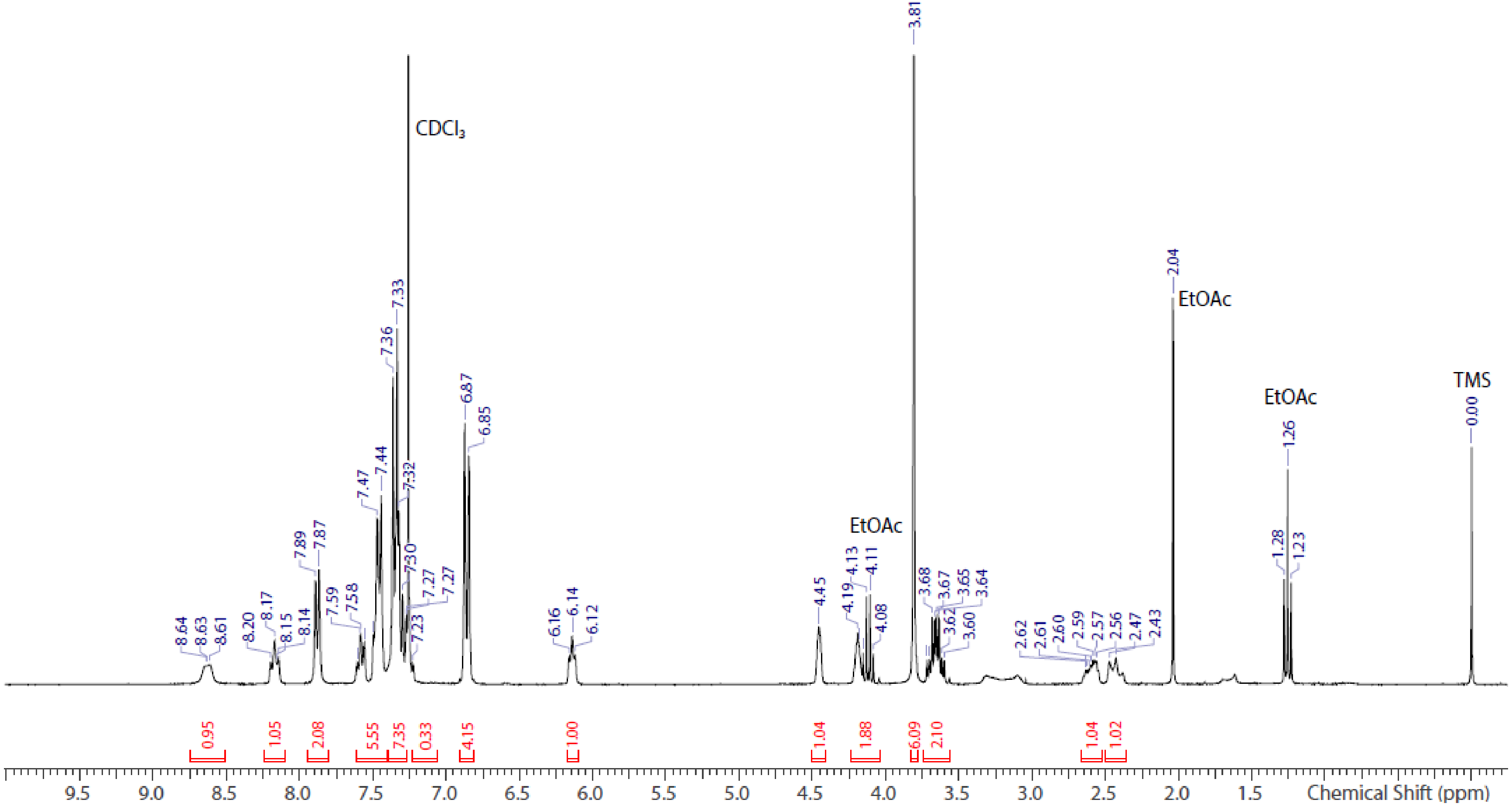


**Spectrum S21.** [1]H NMR spectrum of dx C **40** ($CDCl_3$)

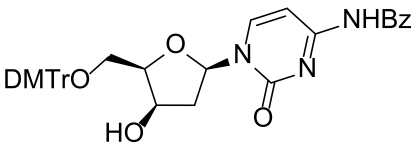


**40**

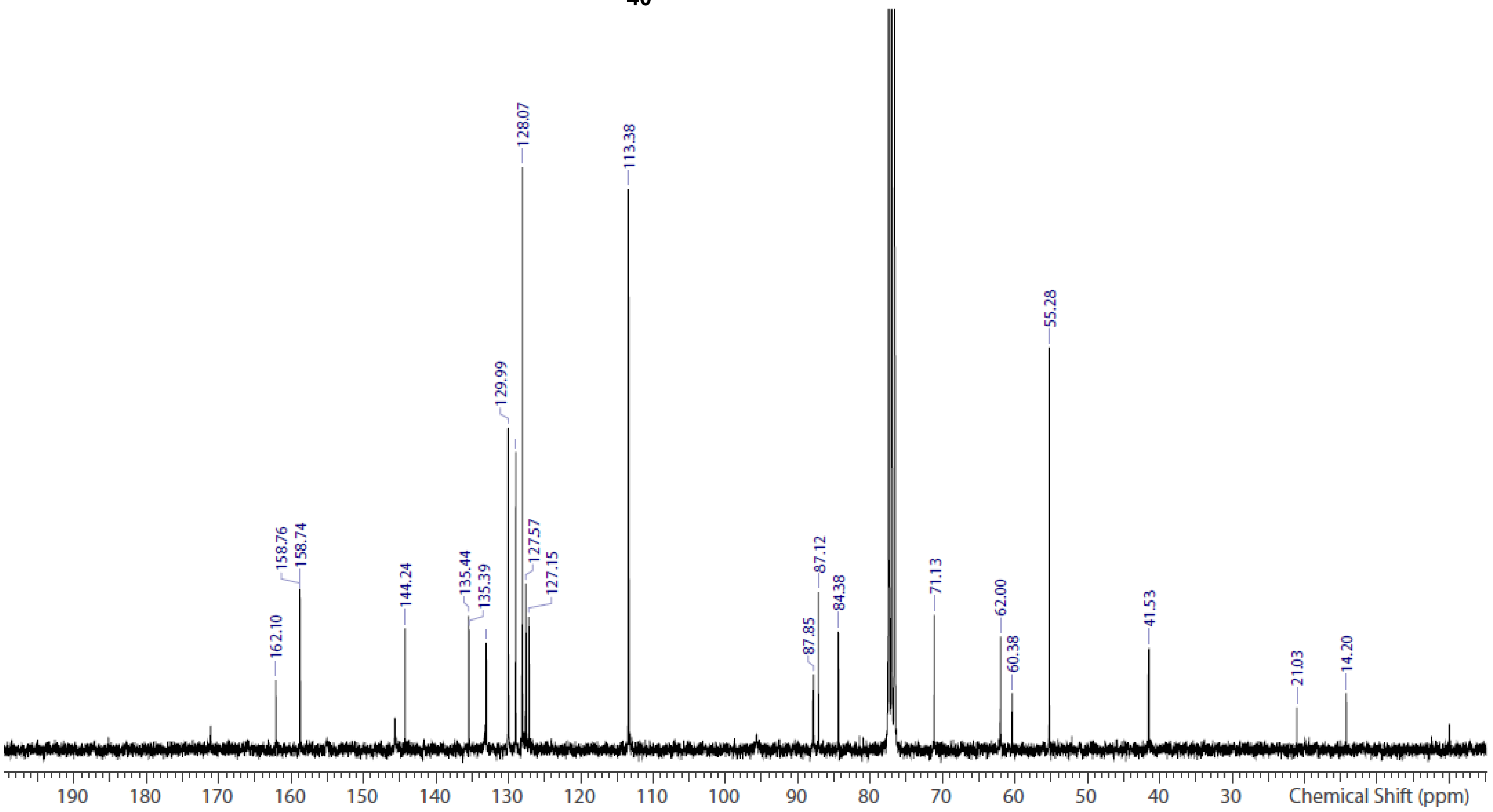


**Spectrum S22.** $^{13}$C NMR spectrum of dx C **40** ($CDCl_3$)

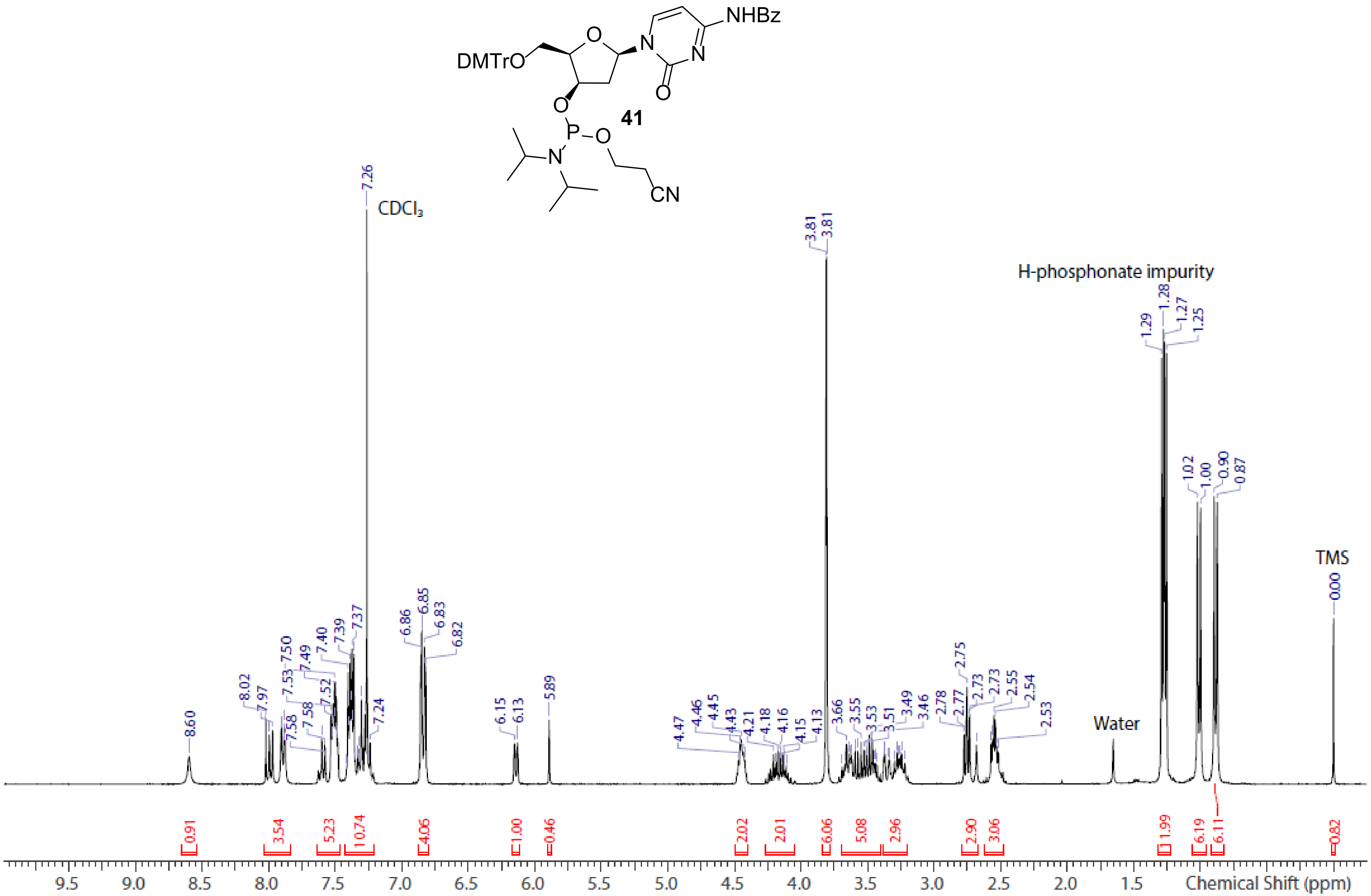


**Spectrum S23.** [1]H NMR spectrum of dx C phosphoramidite **41** ($CDCl_3$)

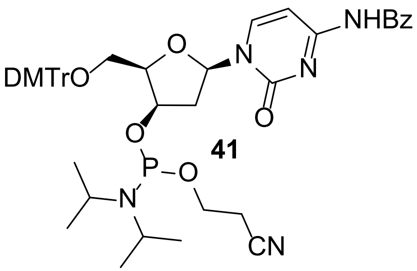


41

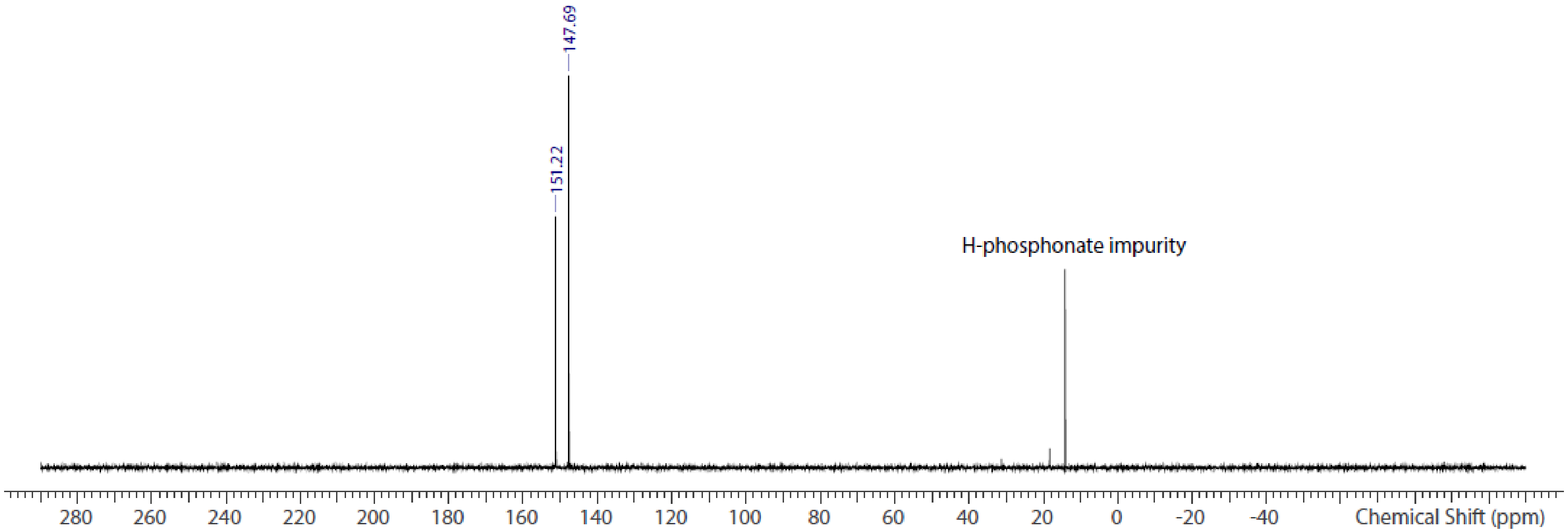


**Spectrum S24.** $^{31}P$ NMR spectrum of dx C phosphoramidite **41** ($CDCl_3$)

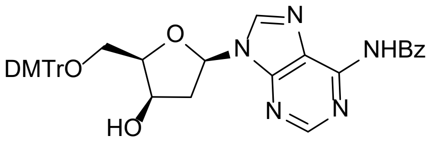


**44**

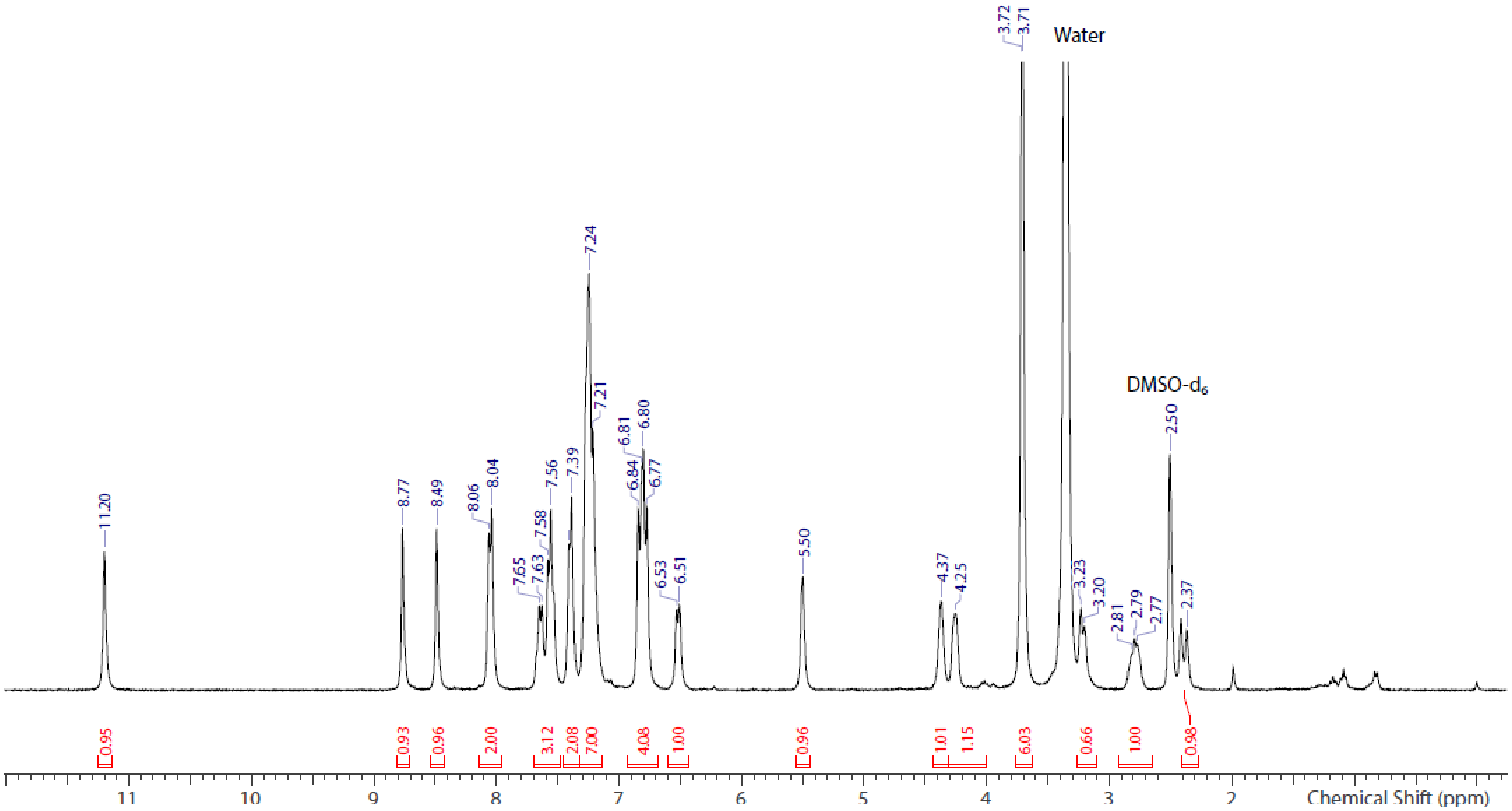


**Spectrum S25.** $^{1}$H NMR spectrum of dx A **44** (DMSO-$d_6$)

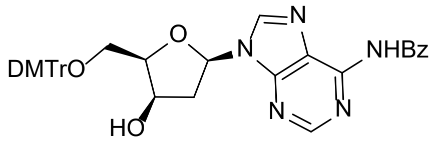


**44**

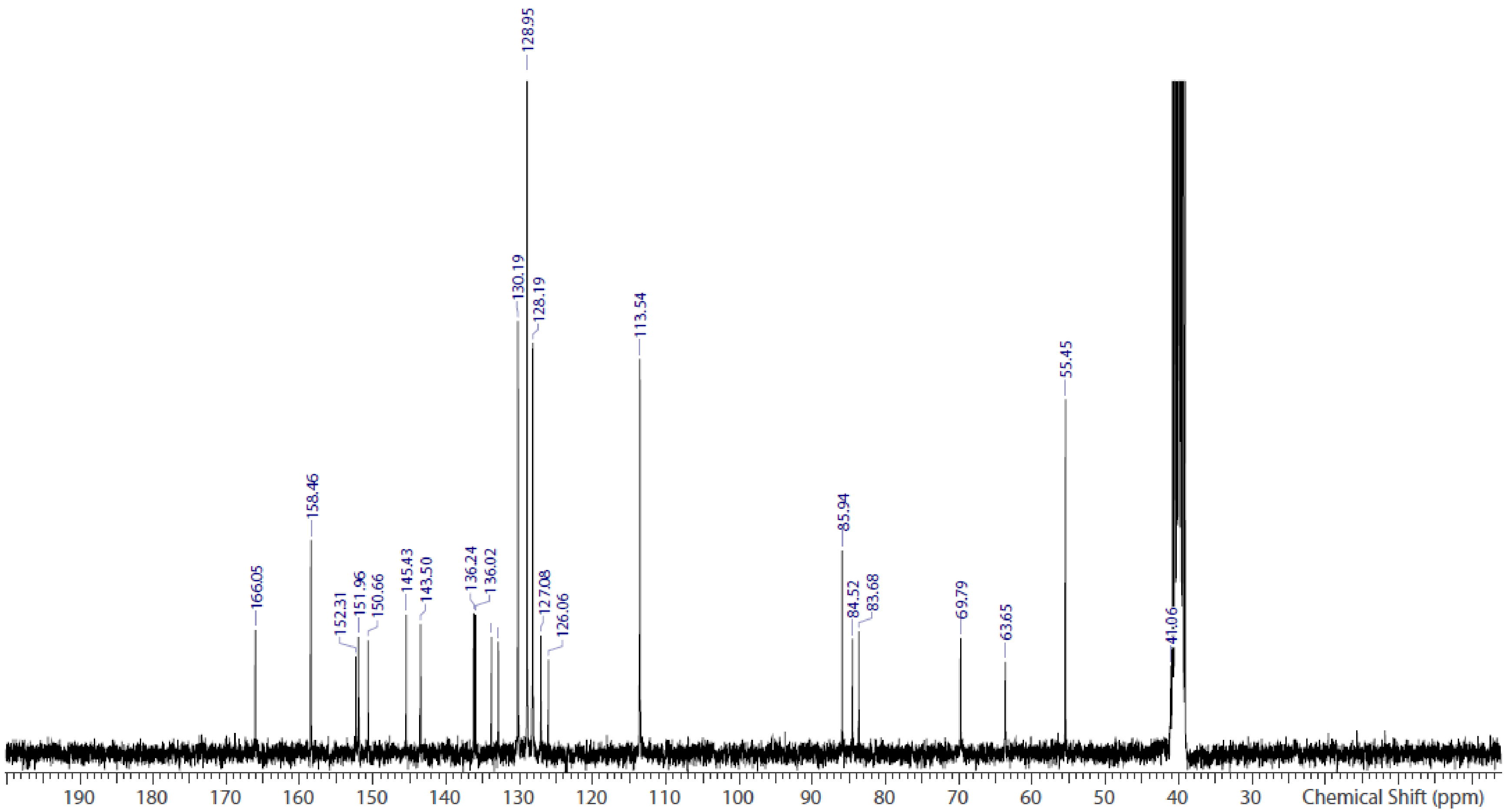


**Spectrum S26.** $^{13}C$ NMR spectrum of dx A **44** (DMSO-$d_6$)

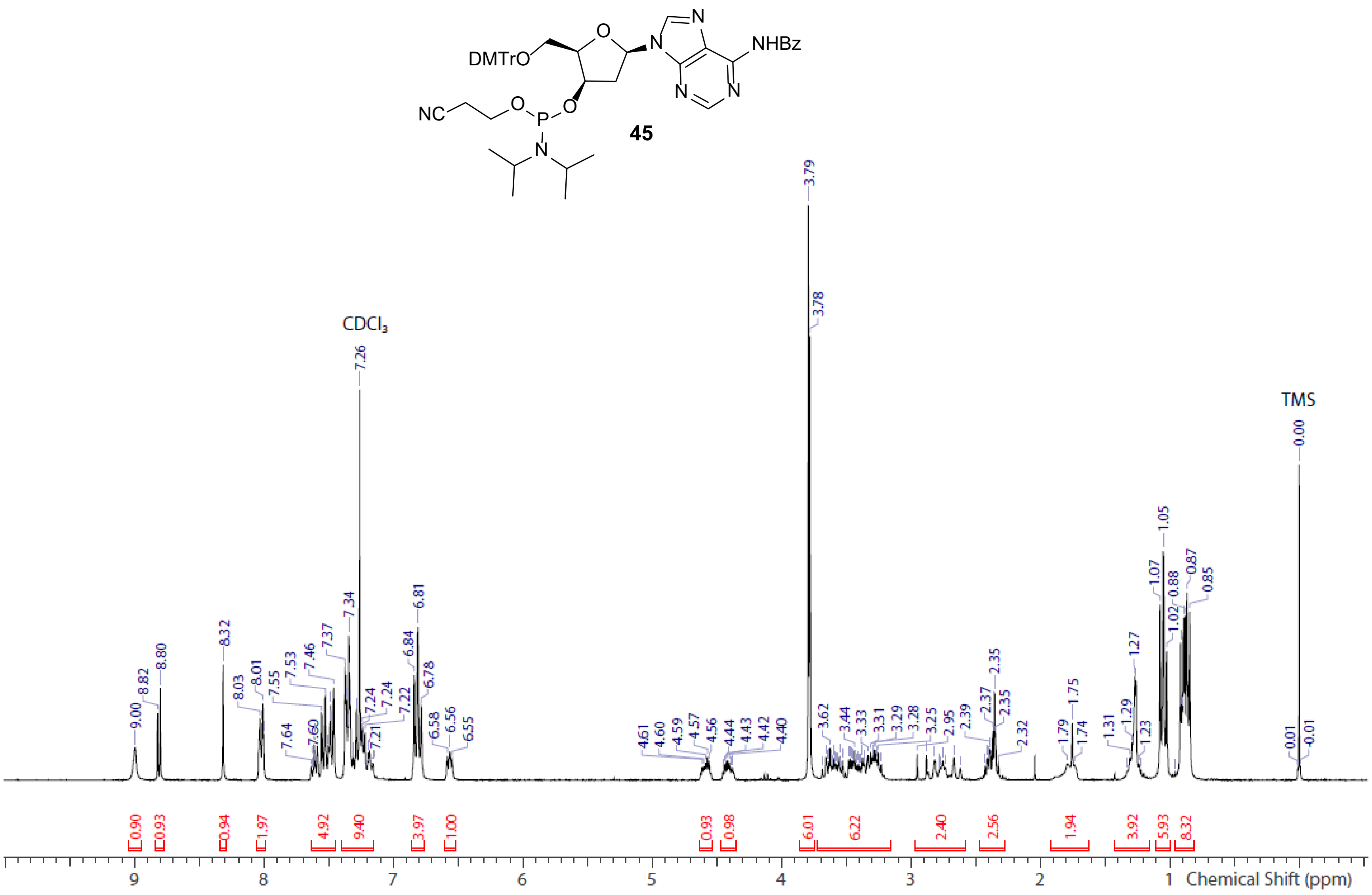


**Spectrum S27.** $^{1}$H NMR spectrum of dx A phosphoramidite **45** ($CDCl_3$)

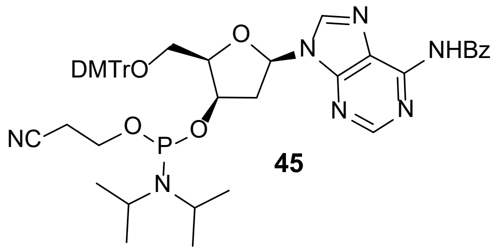


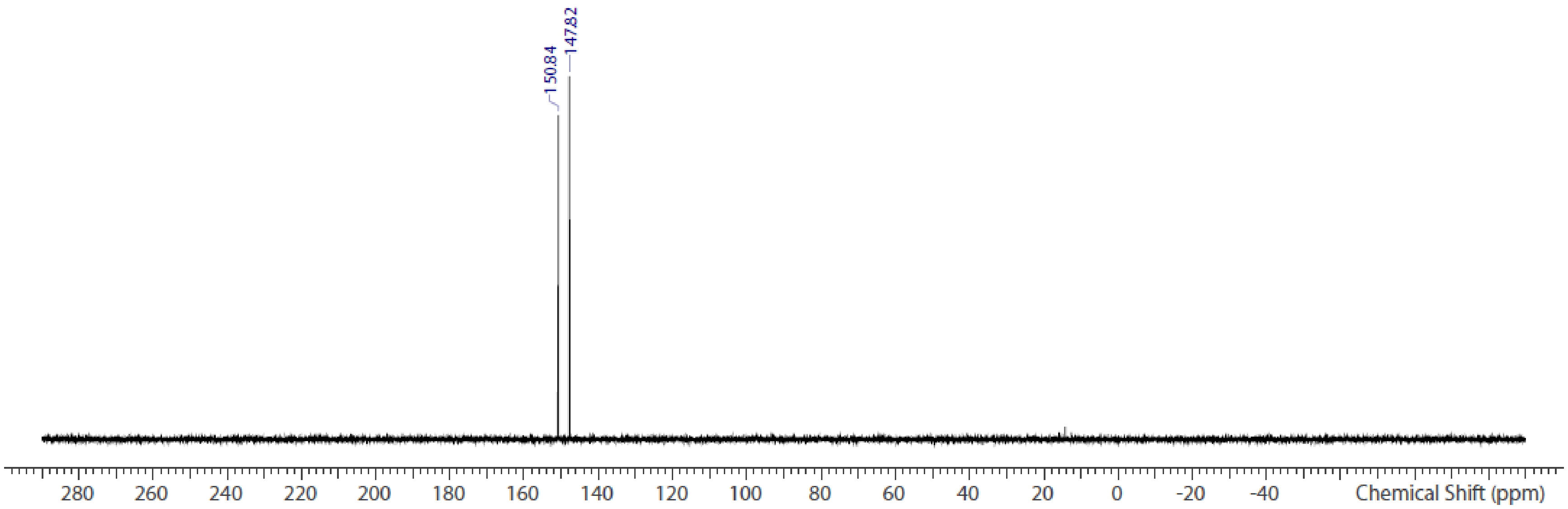


**Spectrum S28.** $^{31}P$ NMR spectrum of dx A phosphoramidite **45** ($CDCl_3$)

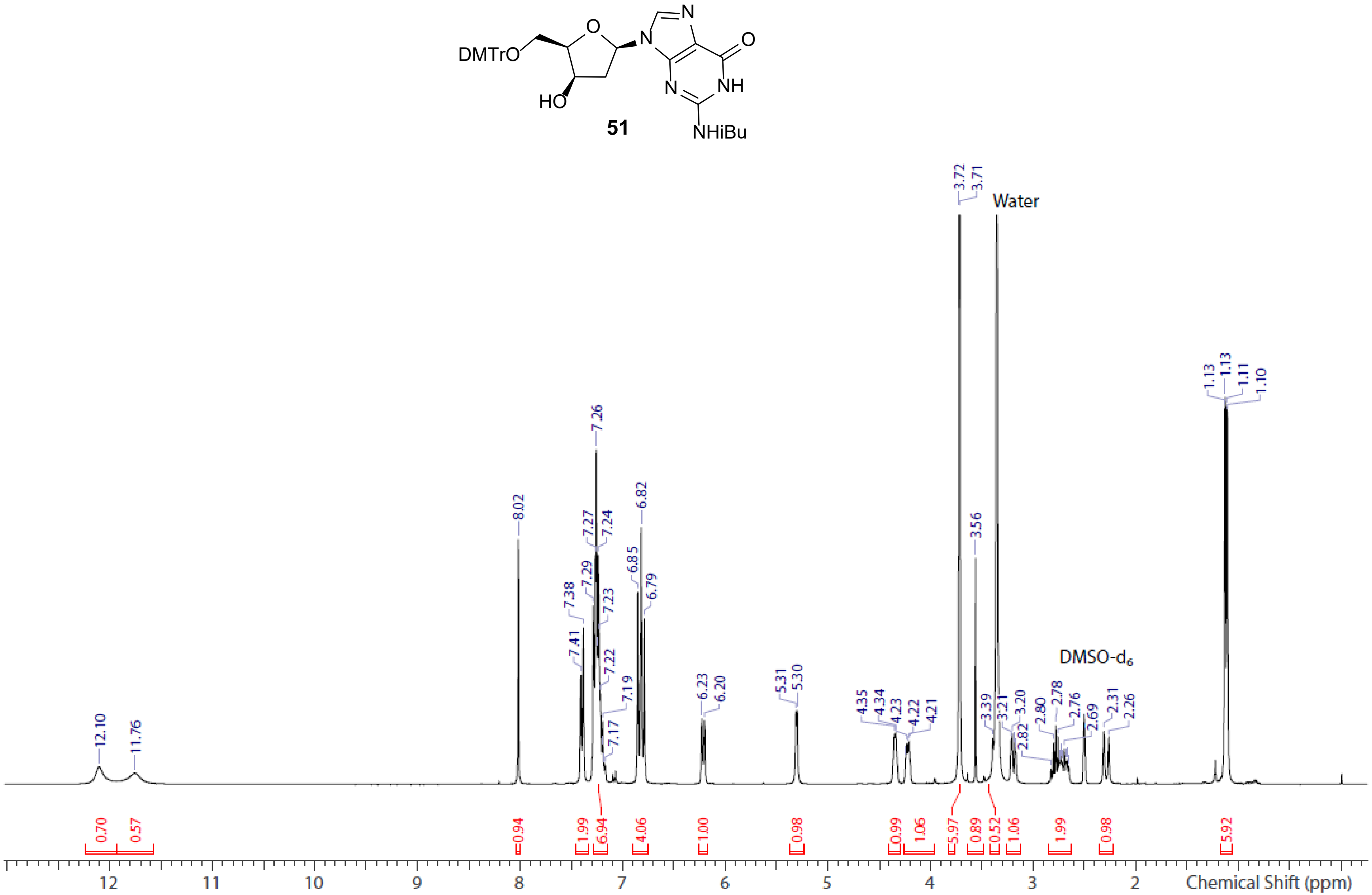


**Spectrum S29.** [1]H NMR spectrum of dx G **51** (DMSO-$d_6$)

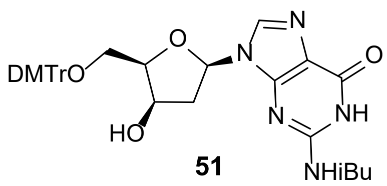


**51**

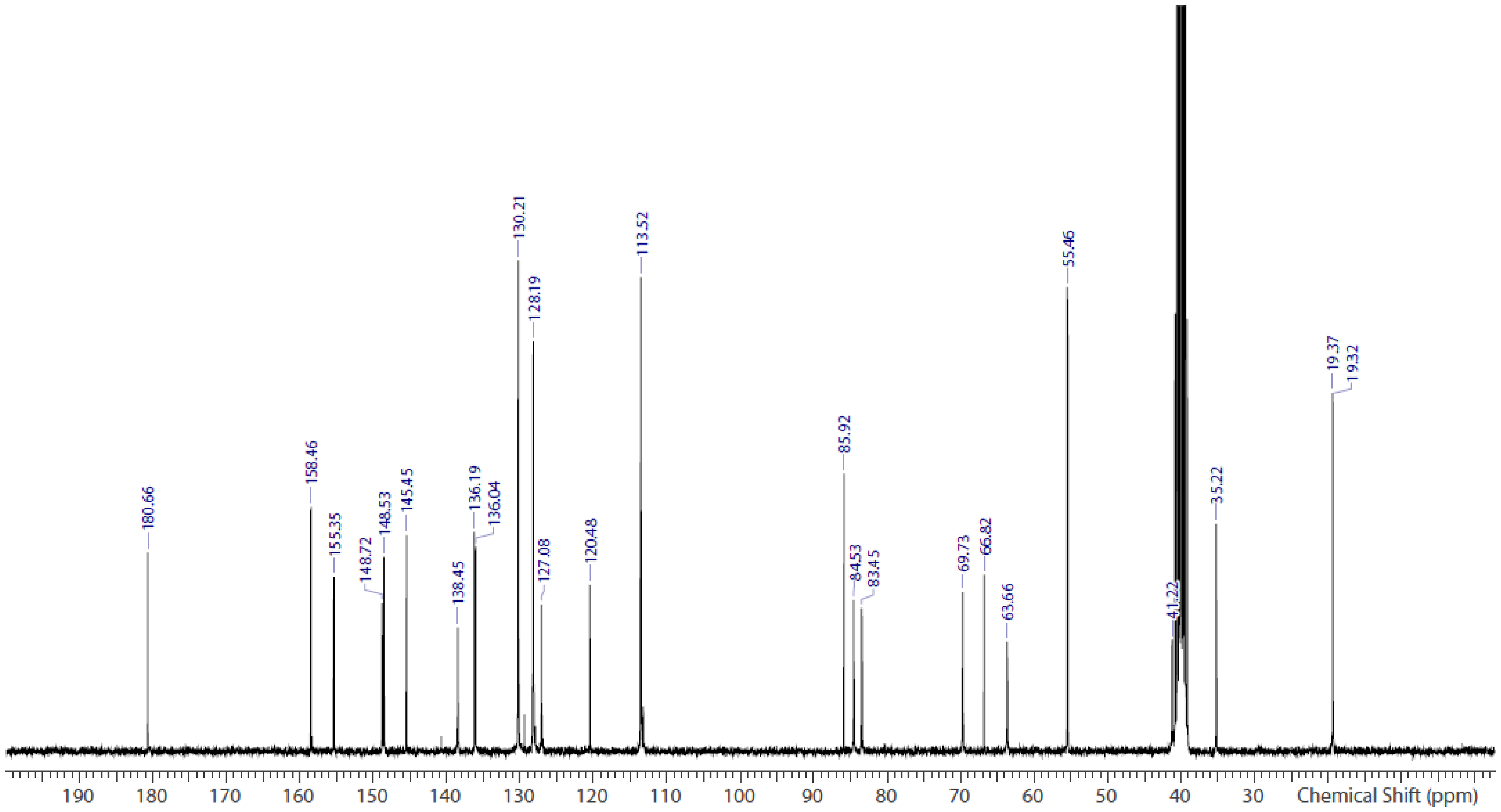


**Spectrum S30.** $^{13}$C NMR spectrum of dx G **51** (DMSO-$d_6$)

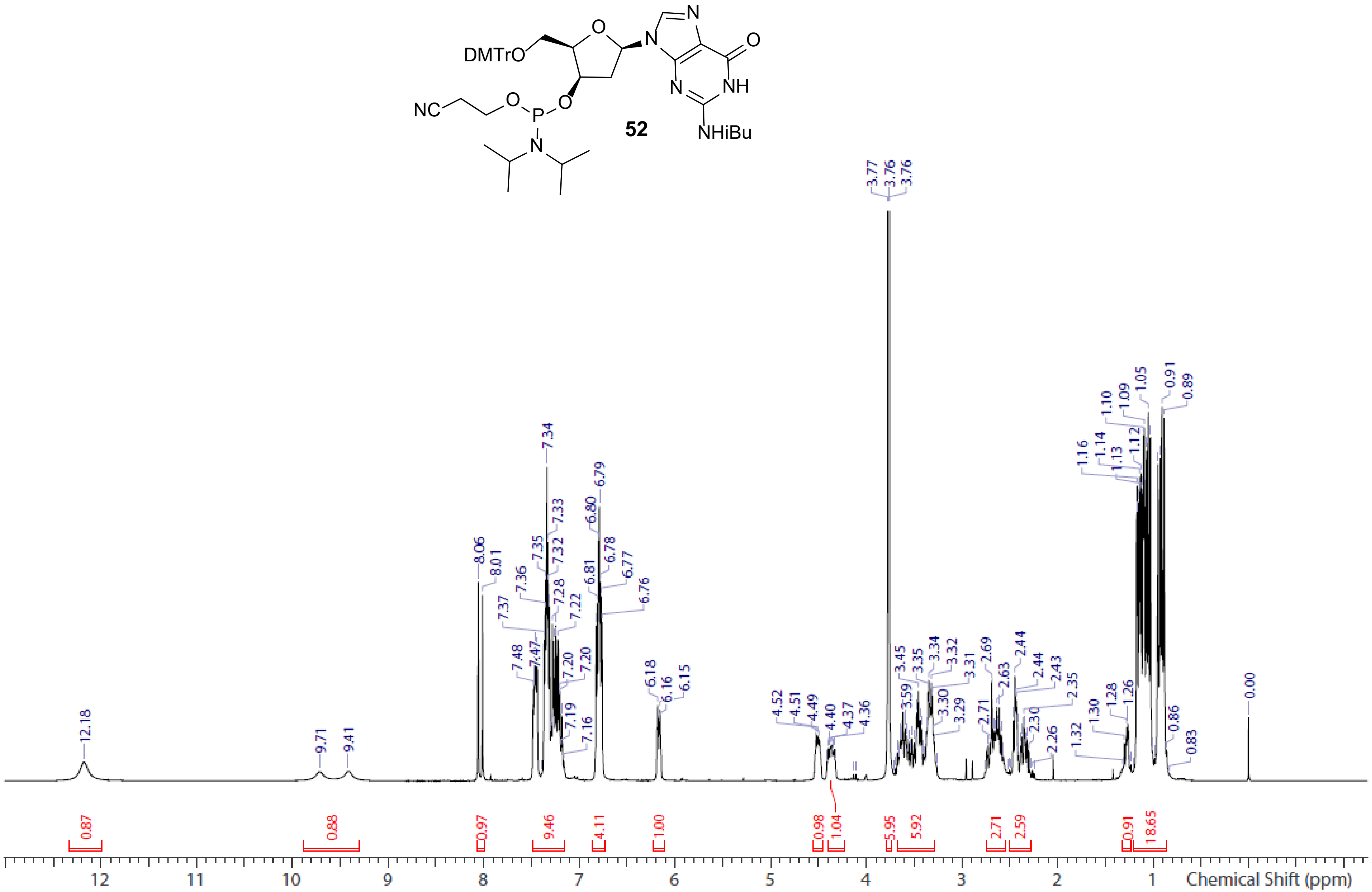


**Spectrum S31.** $^{1}$H NMR spectrum of dx G phosphoramidite **52** ($CDCl_3$)

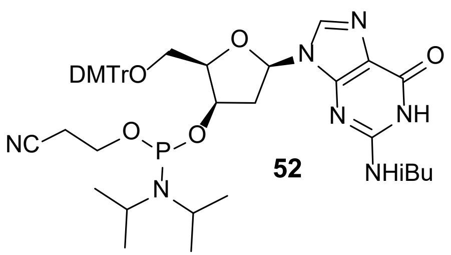


52

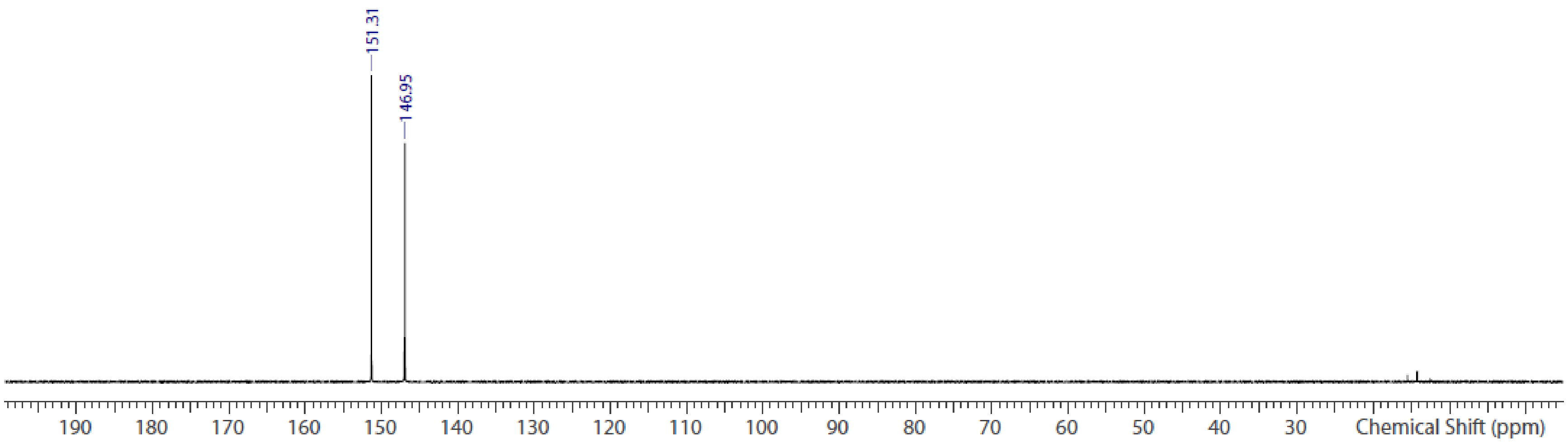


**Spectrum S32.** $^{31}$P NMR spectrum of dx G phosphoramidite **52** ($CDCl_3$)
